# Solvation dynamics-powered structure and function of multi-molecular cellular systems exemplified by non-equilibrium cereblon-degrader-CK1α ternary complex formation

**DOI:** 10.1101/2021.10.29.466510

**Authors:** Hongbin Wan, Vibhas Aravamuthan, Sarah Williams, Charles Wartchow, José S. Duca, Robert A. Pearlstein

**Author notes:** Corresponding author: Robert A. Pearlstein, Ph.D. Phone: -871-7293. Novartis Institutes for BioMedical Research, 5959 Horton Street, Suite 900, Emeryville, CA 94608.

## Abstract

Cellular functions are executed via a form of analog computing that is based on the switchable covalent and non-covalent states of multi-molecular fluxes (i.e., time-dependent species/state concentrations) operating in the non-linear dynamics regime. We and others have proposed that the non-covalent states and state transitions of aqueous fluxes are powered principally by the storage and release of potential energy to/from the anisotropic H-bond network of solvating water (which we refer to as the “solvation field”), which is a key tenet of a first principles theory on cellular structure and function (called Biodynamics) that we outlined previously. This energy is reflected in water occupancy as a function of solute surface position, which can be probed computationally using WATMD software. In our previous work, we used this approach to deduce the structural dynamics of the COVID main protease, including substrate binding-induced enzyme activation and dimerization, and product release-induced dimer dissociation. Here, we examine:

1) The general relationships between surface composition/topology and solvation field properties for both high and low molecular weight (HMW and LMW) solutes.
2) The general means by which structural dynamics are powered by solvation free energy, which we exemplify via binding between the E3 ligase CUL4A/RBX1/DDB1/CRBN, LMW degraders, and substrates. We propose that degraders organize the substrate binding surface of cereblon toward complementarity with native and neo substrates, thereby speeding the association rate constant and incrementally slowing the dissociation rate constant.
3) Structure-activity relationships (SAR) based on complementarity between the solvation fields of cognate protein-ligand partners exemplified via LMW degraders.

## Introduction

The equilibrium state distributions of biomolecular species present in the in vitro regime are widely assumed to extend to the native cellular regime. However, many cellular functions depend on time- and space-dependent variation in the concentrations/number densities of their constituent molecular and ionic species (which we refer to as “fluxes”), where equilibrium principles and metrics (e.g., ΔG, IC_50_, K_d_) break down. Molecular fluxes can be described symbolically as:

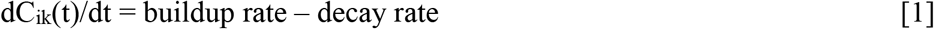

where C is the time-dependent concentration of a given molecular species i within a given compartment k that builds and decays at specific rates (which we refer to as “extrinsic rates”) that are distributed over one or more intramolecular, intermolecular, and covalently modified states according to their individual rates of buildup and decay (which we refer to as “intrinsic rates”).^1^ The most populated states are those exhibiting similar intrinsic and extrinsic buildup and decay rates.^1^ The equilibrium regime is the special case in which the species concentrations are fixed, dC(t)/dt = 0 and the buildup and decay rates are equal. We have developed a multi-scale first principles, physics-based theory called Biodynamics describing the general means by which cellular behaviors are conveyed by dynamic multi-molecular and multi-ionic fluxes operating in the non-linear dynamics (NLD) regime (where the behavior of the whole system >> Σ(behaviors of the parts)).^1, 3, 4^ Our theory carries major implications for:

1) Molecular biology, including protein-protein interactions and the relationship between mutation-induced protein dysfunction and cellular dysfunction/disease.
2) Cell biology, including the means by which cellular functions are conveyed by multi-flux systems.
3) Drug R&D, including structure-activity relationship (SAR) analysis of low molecular weight (LMW) analogs, pharmacokinetic-pharmacodynamic (PK-PD) analysis, and structure-based drug design.

In our previous work, we applied Biodynamics principles and WATMD software (a first principles method for probing water occupancy across solute surfaces) to qualitatively deduce the solvation free energy-powered non-covalent non-equilibrium intra- and intermolecular state transitions of the COVID main protease (M^pro^) and dynamic blockade of the human ether-a-go-go related gene potassium channel (hERG).^4, 5^ In this work, we:

1) Briefly review the basic properties of fluxes and the common architecture of multi-flux systems exemplified by the cereblon-mediated proteasomal protein degradation pathway (the sink for many protein fluxes).
2) Overview our ongoing progress in deciphering the general mechanisms by which solvation free energy is transduced into the intrinsic rates of non-covalent intra- and intermolecular state transitions (vis-à-vis the status quo paradigm) under aqueous non-equilibrium conditions, which we extend to protein folding and the low molecular weight (LMW) regime.
3) Expand the application of our theory and WATMD to deduce the solvation free energy-powered <u>non-equilibrium</u> binary and ternary states of cereblon (the substrate-recruiting subunit of the CUL4A/RBX1/DDB1/CRBN E3 ligase) bound to native and synthetic LMW degraders and neo substrates.

### The moving parts of cellular systems consist of molecular and ionic fluxes multiplexed into non-linear dynamic systems

Whereas molecules and ions constitute the tangible <u>moving parts</u> (i.e., fluxes) of cellular systems, the <u>intangible behaviors</u> of such systems <u>emerge</u> from NLD operation. Cellular systems respond to perturbations via acceleration or deceleration of specific fluxes and/or the state transitions thereof.^2^ Molecular fluxes are generated in a source → sink fashion powered by transducible covalent free energy sources (e.g., ATP) and undergo dynamic solvation-powered non-covalent intra- and intermolecular state transitions (including enzyme-substrate binding) throughout their lifetimes (Figure 1A). Protein fluxes, the workhorses of cellular function, are produced via transcription and translation and cleared via proteasomal, lysosomal and other degradation mechanisms (Figure 1B) (noting that fast fluxes depend on activation of pre-synthesized pro- protein or zymogen pools). The lifetimes of proteins that decay via proteasomal degradation are governed by the relative rates of poly-ubiquitylation versus deubiquitylation catalyzed by E3 ligases and deubiquitylases (DUBs), respectively (and possibly other mechanisms) that we address later in this work.

**Figure 1.**
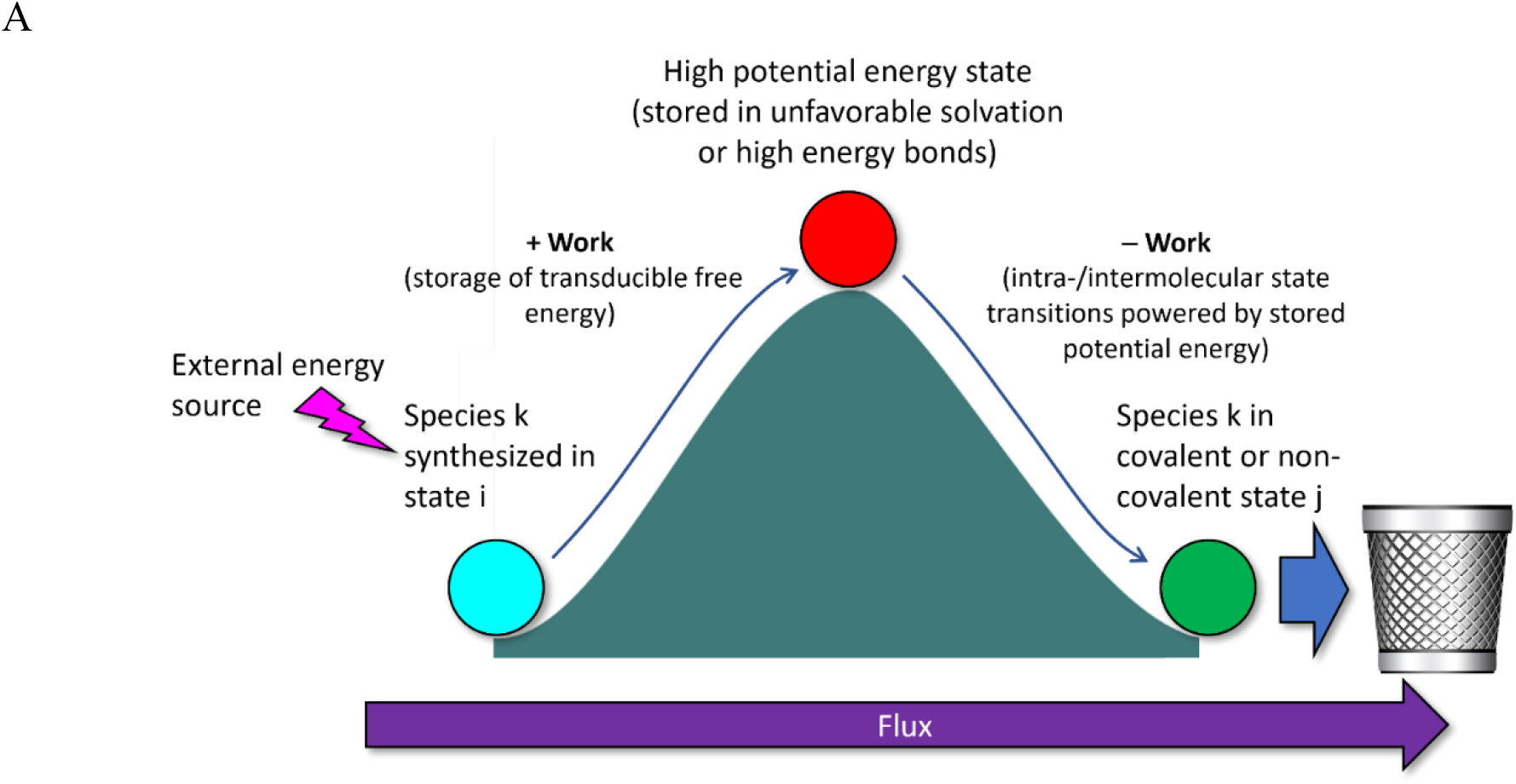
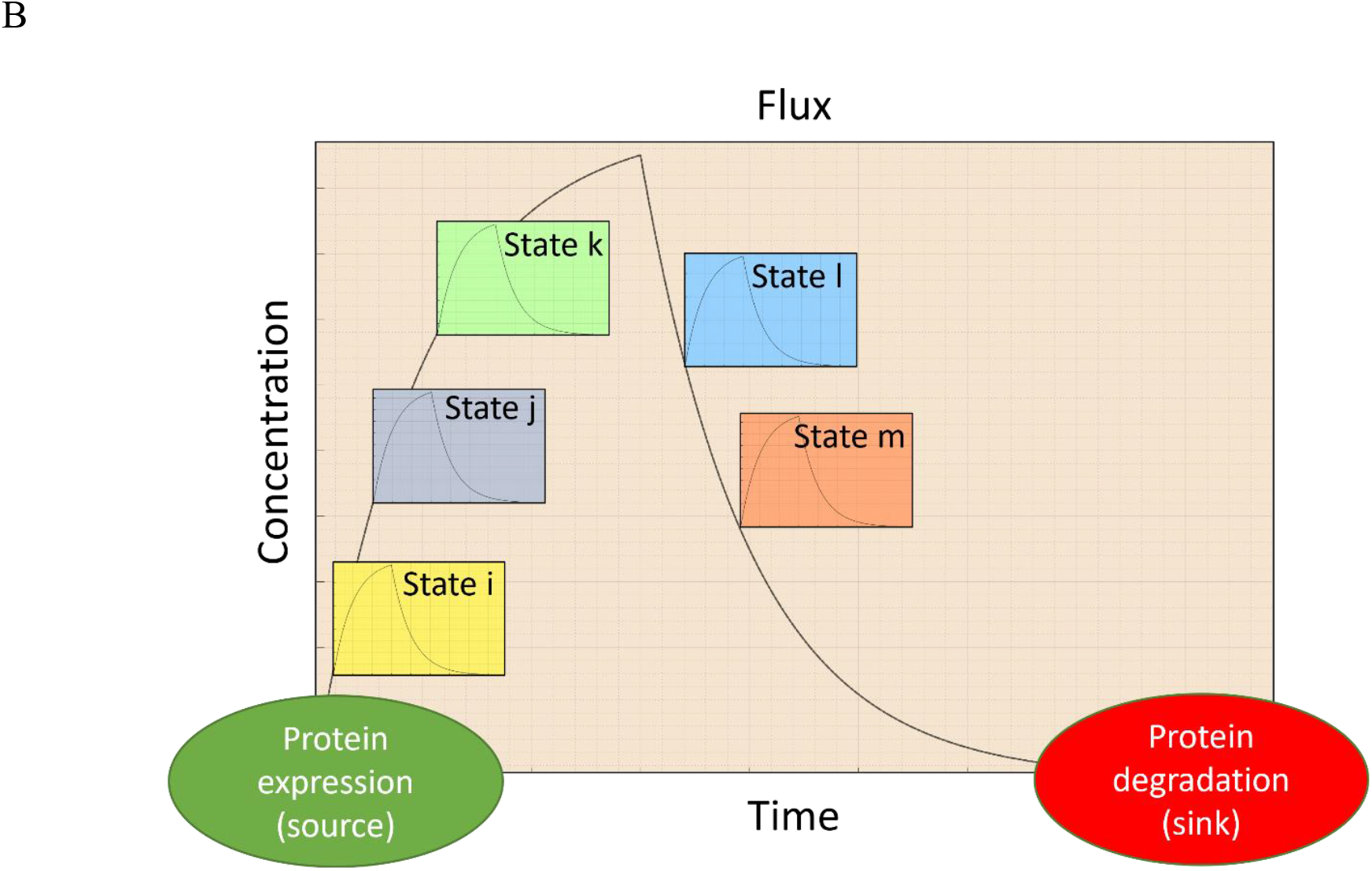
(A) Fluxes are powered by various forms of transducible potential and kinetic energy, beginning with synthesis or inward intercompartmental translocation, storage of transducible free energy forms (including unfavorable solvation and high energy bonds), state transitions, and degradation or outward intercompartmental translocation. (B) Protein fluxes build via expression and decay via degradation in a source-sink fashion and undergo a range of covalent and non-covalent solvation free energy-powered intra- and intermolecular state transitions that build and decay throughout their lifetimes.

The states of each flux are populated dynamically according to their rates of buildup and decay, the <u>physical form</u> of which we refer to as “molecular differential equations” (MDEs), and the canonical symbolic form of which is given in equation 1. Fluxes exist in one or more intra- and intermolecular states that build and decay over time, analogous to the flow of fluid to/from the fixtures of a plumbing system, which depend on the rates of entry and exit to/from the system (analogous to the extrinsic flux rates) and the rates of entry and exit to/from each fixture (analogous to the intrinsic flux rates).^2^ The non-covalent and enzyme-mediated covalent states of a given flux necessarily build and decay on the overall timescale of the cellular function in which they participate (e.g., the open, inactivated, and closed states of cardiac ion channels build and decay on the ∼350 ms timescale of the action potential^6, 7^). Cellular behaviors consist of the “physical solutions” of networked time- and space-dependent multi-molecular or multi-ionic fluxes (Figure 2), which we refer to as “physical math”.^2^ Cellular systems comprised of three or more multiplexed MDEs operate in the NLD regime, giving rise to emergent behaviors that are >> Σ(behaviors of the moving parts). NLD systems lack analytical solutions and must be solved computationally via symbolic mathematical simulations or physically via analog computing in the case of living cells (where the solutions/outputs consist of various dynamic behaviors). The “cellular operating system” (CellOS) is thus analog, wherein the hardware and software are one and the same. From this perspective, fluxes act as analog signal carriers, analogous to amplitude and frequency modulated (AM and FM) radio waves. Cellular behaviors are initiated by intra- and/or extracellular input signals that propagate through and switch the states of the corresponding network via alteration of one or more extrinsic and/or intrinsic flux rates.^2^ Abnormal genetic mutation-induced alterations of these rates constitutes the fundamental basis of cellular disease (e.g., slowed protein synthesis has been shown to play a role in Huntington’s disease^8^). Molecular and ionic fluxes build and decay <u>exponentially</u> in time (i.e., the solutions of both the physical and symbolic forms of the MDEs), which has crucial implications for cellular pathway architecture^1–3^ (Figure 3). Counter- balanced flux-anti-flux pairs that we refer to as “Yins” and “Yangs” are necessitated by such behavior.

**Figure 2.**
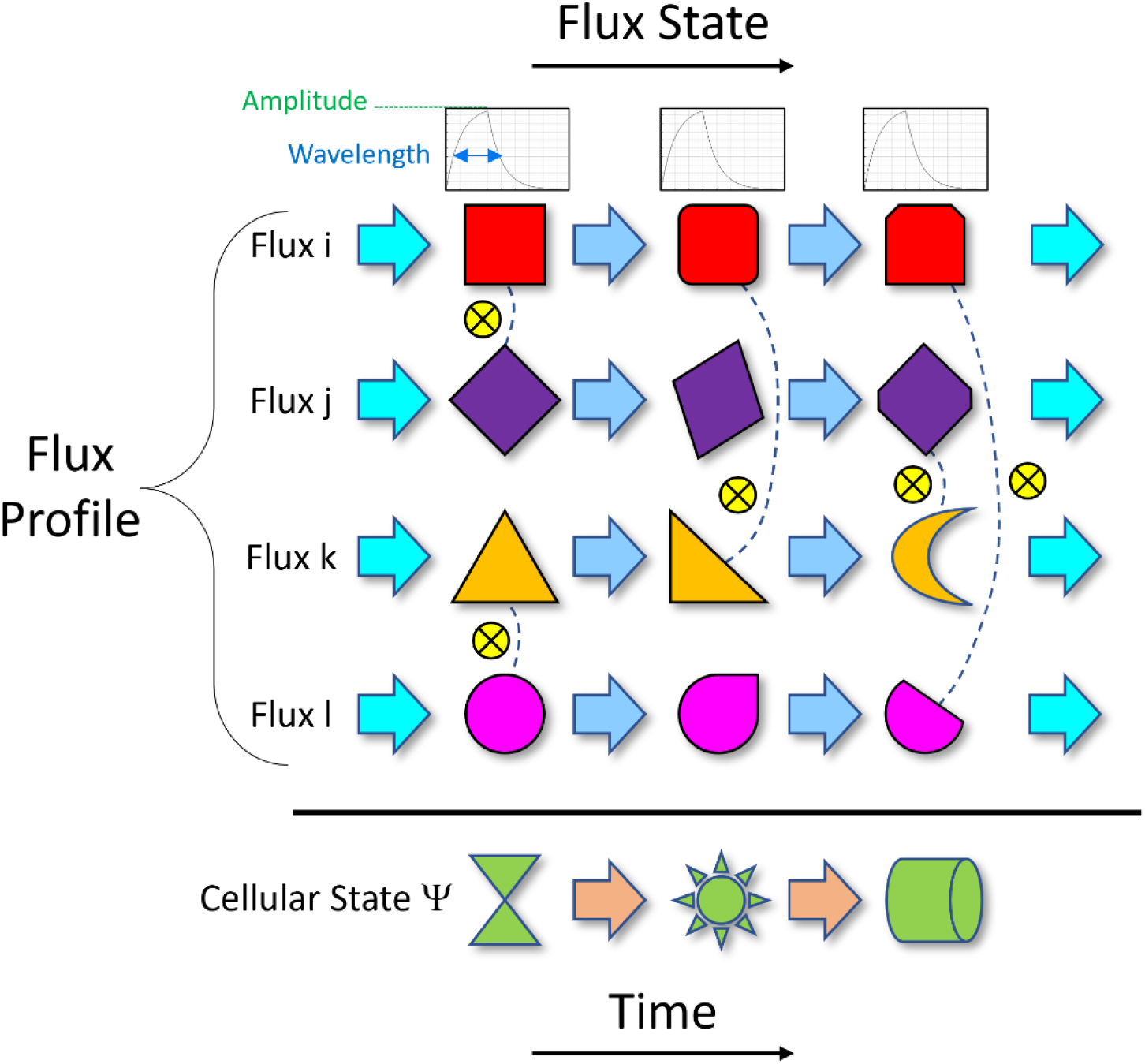
Time-dependent cellular states are defined by the dynamic state transitions of their constituent multi-flux systems, which are subject to a variety of transient perturbations/signals (typically initiated by stimulation of intracellular and extracellular receptors). Each participating flux (denoted by the geometric shapes along the time axis) encodes a signal in the amplitudes and wavelengths of its covalent and non-covalent intra- and intermolecular states, intrinsic and extrinsic rates, and state-specific inter-connections (analogous to a switchboard, denoted by dotted lines). Non-linear behaviors result from interdependence between the dynamic states and multiplexed configurations of each multi-flux system.

**Figure 3.**
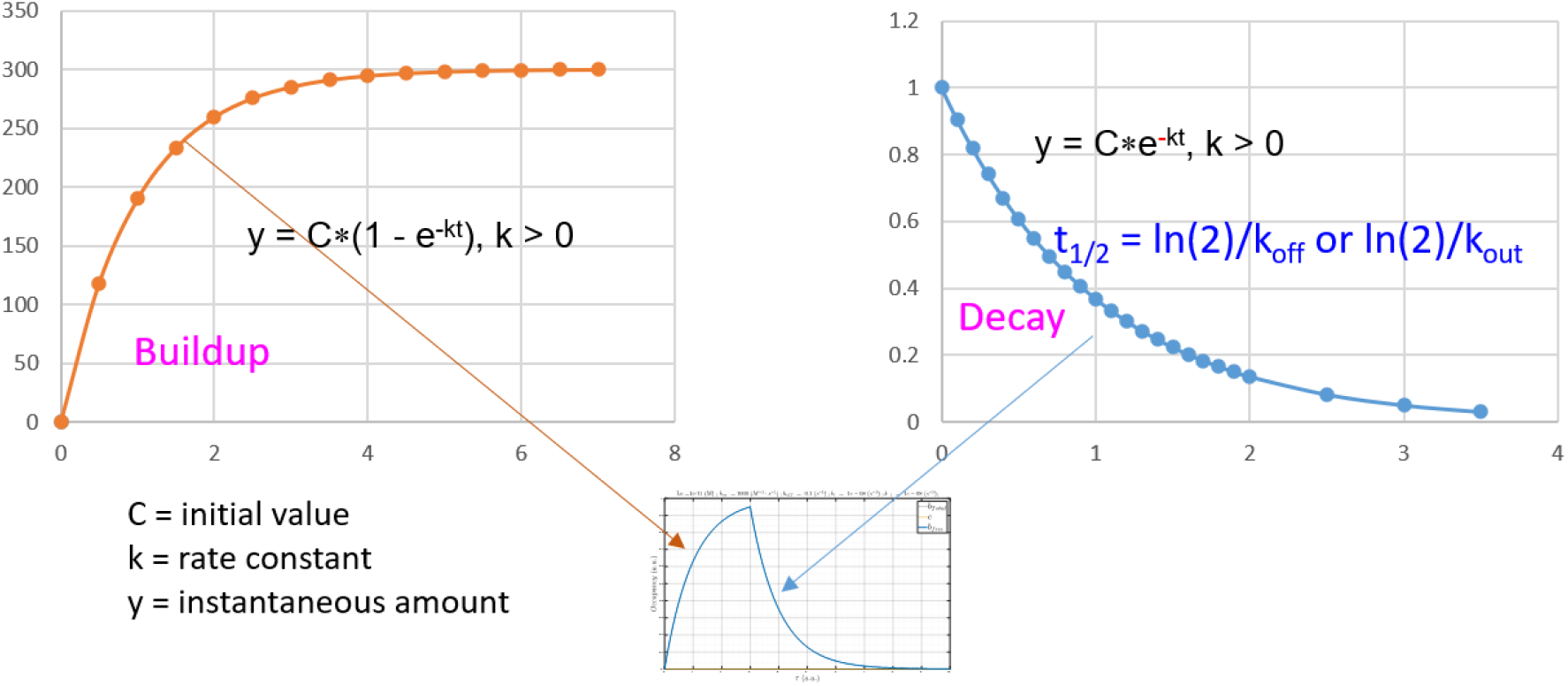
Fluxes and their populated states build and decay exponentially in time.

The instantaneous net rate of product formation in the case of enzyme Yin-Yang pairs (e.g., kinase-phosphatase, ligase-ubiquitinase) depends on the <u>relative</u> Yin and Yang activities, which operate in a dynamic push-pull fashion (a molecular tug-of-war) governed by the relative Yin and Yang flux rates/amplitudes, together with the phasing of the Yang buildups relative to the corresponding Yin-catalyzed products (Figure 4). Yang buildups, in turn, are driven by de novo expression or activation by downstream species (e.g., phosphatase activation by a downstream kinase), which we refer to as “Clocks”. The Yin-Yang balance is tipped in the forward and reverse directions by slow and fast Clocks, respectively. The states of each Yin-Yang-Clock module are governed by the rates of entry and exit to/from the available intra- and intermolecular states of the module.

**Figure 4.**
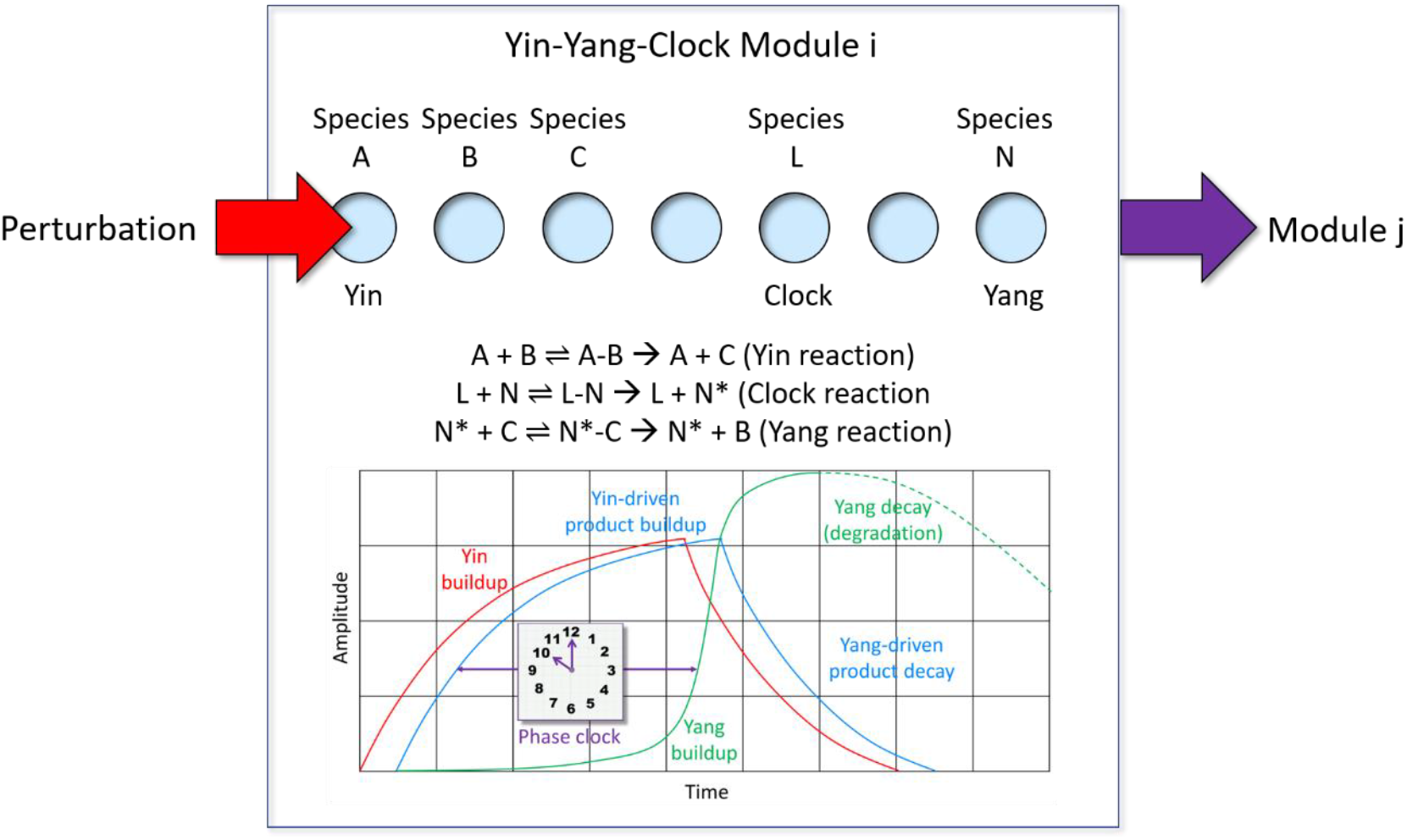
Cellular pathways are comprised of sets of sequential enzyme-catalyzed reactions (fluxes) among the participating species (blue filled circles, denoted as A, B, C, etc.). Pathways are putatively organized into sets of Yin-Yang-Clock modules (exemplified here for a hypothetical module i), in which the Yin, Yin product, Clock, and Yang, consist of species A, B, L, and N, respectively. The forward and reverse fluxes are maintained in an out-of-phase relationship via the Clock-dependent Yang buildup (green curve) that lags behind that of the corresponding Yin- catalyzed product (blue curve). The Yin-Yang balance is respectively tipped toward the Yin and Yang during the product buildup and decay phases.

### The status quo concept of structure-free energy relationships is challenged by Biodynamics theory

Molecular dynamics (MD) simulations, free energy perturbation (FEP), docking scores, and other force-field-based methods have been used extensively for studying the intra- and intermolecular structures and state transitions of proteins and other biomolecules.^9, 10, 11–18, 19–22^ However, the widely held view that non-covalent molecular structure and function are powered by van der Waals, electrostatic, H-bond, hydrophobic, hydrophilic, π-stacking, π-cation, and other types of interatomic energy contributions is being increasingly questioned by our findings^2–4, 23, 24^ and those of others.^25^ Mounting evidence suggests that, under both static equilibrium and dynamic non-equilibrium conditions, the energetically preferred sterically-accessible structural states of aqueous HMW and LMW biomolecular solutes and non-covalent HMW-LMW and HMW-HMW complexes are powered principally by solvation free energy, and in particular, solvation H-bond free energy relative to that of bulk solvent. Molecular structures reside within a Goldilocks zone of favorable and unfavorable free energy contributions comprised of:

1) Stabilizing/H-bond enriched solvation (“cold spots”), the free energy of which is significantly exceeded by that of bulk solvent.
2) Destabilizing/H-bond depleted solvation (“hot spots”), the free energy of which significantly exceeds that of bulk solvent.

Overly H-bond enriched states are non-rearrangeable (e.g., rigid protein structures), whereas overly H-bond depleted states are poorly or randomly populated (manifesting as disorder, and in extreme cases, precipitation or aggregation). Furthermore, the barriers to/from a given non-covalent state (denoted as 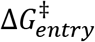 and 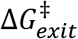, respectively) consist predominantly of the local desolvation and resolvation costs incurred respectively in entering and exiting that state (i.e., 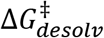 and 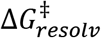), rather than ΔG per se.^3, 4, 23, 24, 26^ Interatomic contact energies are overestimated by empirical force-field (FF) and quantum mechanics (QM)-based methods relative to the H-bond free energy of liquid water, consistent with the vastly higher melting and boiling points of water <u>per unit molecular volume</u> compared with those of liquid organic substances.^3^ The predominant contribution of solvation free energy to ΔG is therefore underestimated in classical interatomic contact models, resulting in:

1) Poor structure <u>predictions</u> based on calculated free energies.
2) Inapplicability to the non-equilibrium cellular setting.
3) Incorrect or suboptimal <u>interpretation</u> of structure-free energy relationships derived from measured data (e.g., kinetic barriers and rate constants measured via surface plasmon resonance, ΔG measured via isothermal titration calorimetry, and biochemical assay data).

Pure liquid bulk water behaves as a high density, isotropic H-bond network to which each water molecule contributes an average of ∼3.5 moderately strong H-bonds.^3^ The H-bond network of liquid water resides within a Goldilocks zone of fluidity and strength, behaving as a quasi-solid when subjected to high frequency perturbations. Water minimizes its enthalpic and entropic free energy contributions when perturbed by HMW and LMW solutes, air, surfaces, cells, etc. relative to those of bulk solvent by:

1) Altering its rate of entry/exit to/from water-non-water interfaces and bulk solvent.
2) _2)_ Increased proton dissociation.^3^
3) Ordering into a wide range of non-isotropic two- and three-dimensional H-bond network geometries in conformance with the local water-solute interfacial geometry.
4) Driving non-covalent solute rearrangements (including protein folding and binding) toward the minimum <u>solvation</u> H-bond free energy, which is incorrectly attributed under the conventional wisdom to hydrophilic, hydrophobic, and other types of interatomic interactions.

Mounting evidence suggests that the <u>primary</u> free energy source powering solute structure and function consists of solvation free energy (water H-bond free energy, in particular), rather than interatomic contacts.^1, 8, 24–26^ A spectrum of free energies, ranging from favorable to unfavorable relative to that of bulk solvent, is stored in the H-bond network of solvating water as a function of local solute surface structure and composition (Figure 5), as follows:

1) **Strained solvation**: diffuse H-bond/<u>enthalpically depleted, entropically neutral</u> solvation at non-polar or single/weak H-bond donor- or acceptor-containing solute regions, corresponding to a wide range of local environment-dependent positive potential energies.
2) **Partially relaxed solvation**: bulk-like, entropically neutral solvation at moderately polar solute regions, corresponding to zero strain.
3) **Fully relaxed solvation**: H-bond/<u>enthalpically enriched, entropically depleted</u> solvation at multi-donor- and/or acceptor solute environments exhibiting a wide range of local environment-dependent negative/favorable potential energies.

**Figure 5.**
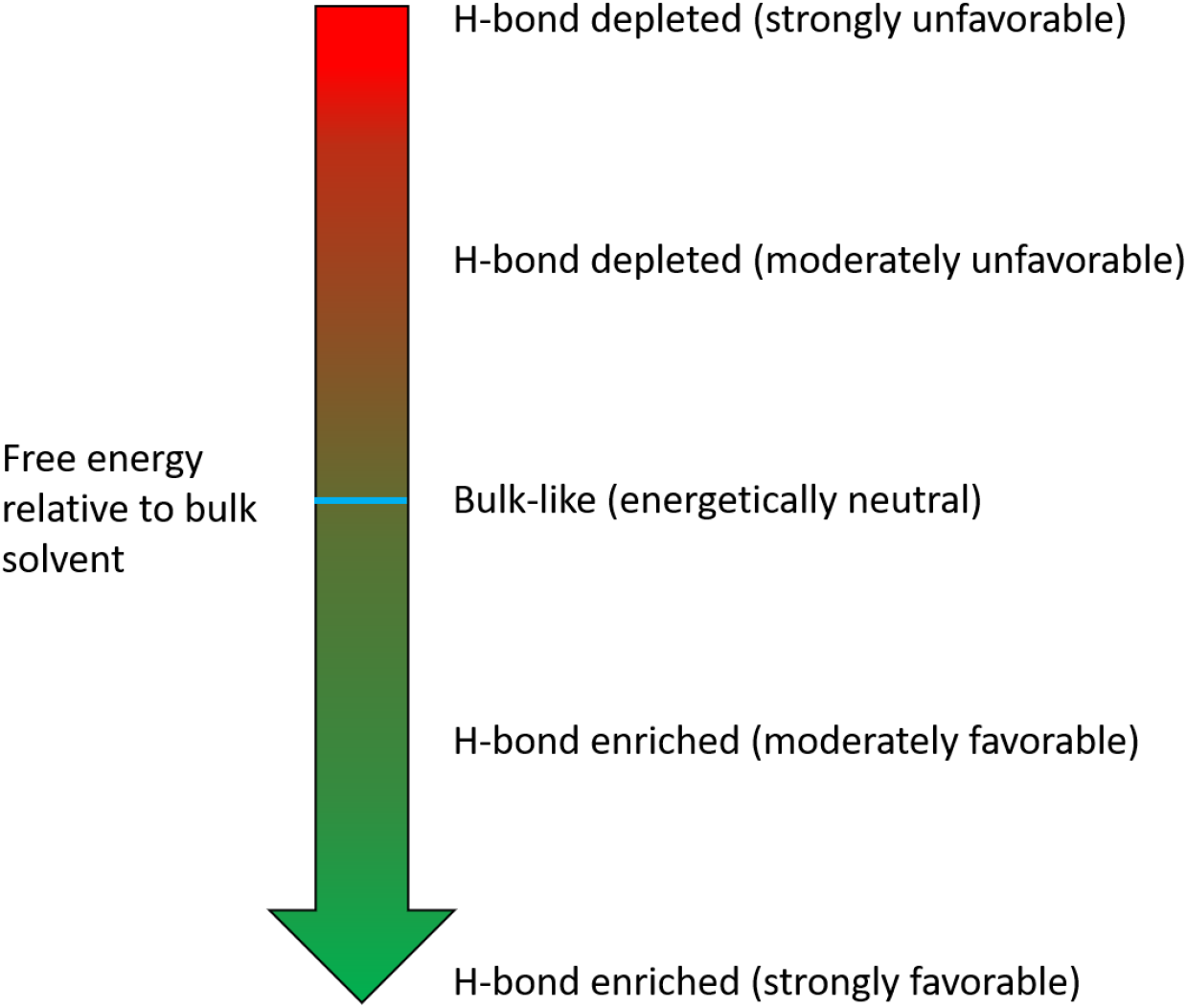
The free energy of bulk solvent is split into a spectrum of favorable and unfavorable contributions across the internal (buried) and external (exposed) surfaces of solutes relative to that of bulk solvent, corresponding to H-bond depleted/strained (potential energy > bulk solvent), bulk- like/relaxed (energetically neutral), and fully relaxed H-bond enriched (potential energy < bulk solvent) water-water-solute configurations. Potential energy stored in H-bond <u>depleted</u> solvation is released via the replacement of such water by <u>non-polar solute groups</u> that themselves incur little desolvation cost during intra- or inter-solute rearrangements.

The absolute free energy (*G^abs^_j_*) of a given non-covalent sterically-accessible state j consists predominantly of the H-bond enriched + depleted solvation free energy of that state relative to that of bulk solvent (i.e.,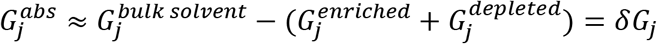). States in which 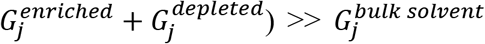 are unfavorable, and therefore poorly populated. The chemical structures of solutes (Figure 6, left) are effectively “imprinted” in their solvating water (Figure 6, right), loosely analogous to the manner in which heavenly bodies are imprinted in space-time, resulting in gravity. From this perspective, the water H-bond network equates to a field (henceforth referred to as the “solvation field”) that responds to the regional loss and gain of H-bonds across solute surfaces (noting that solvating water is capable of energetically resolving solute differences on the order of a single carbon atom)^3^.

**Figure 6.**
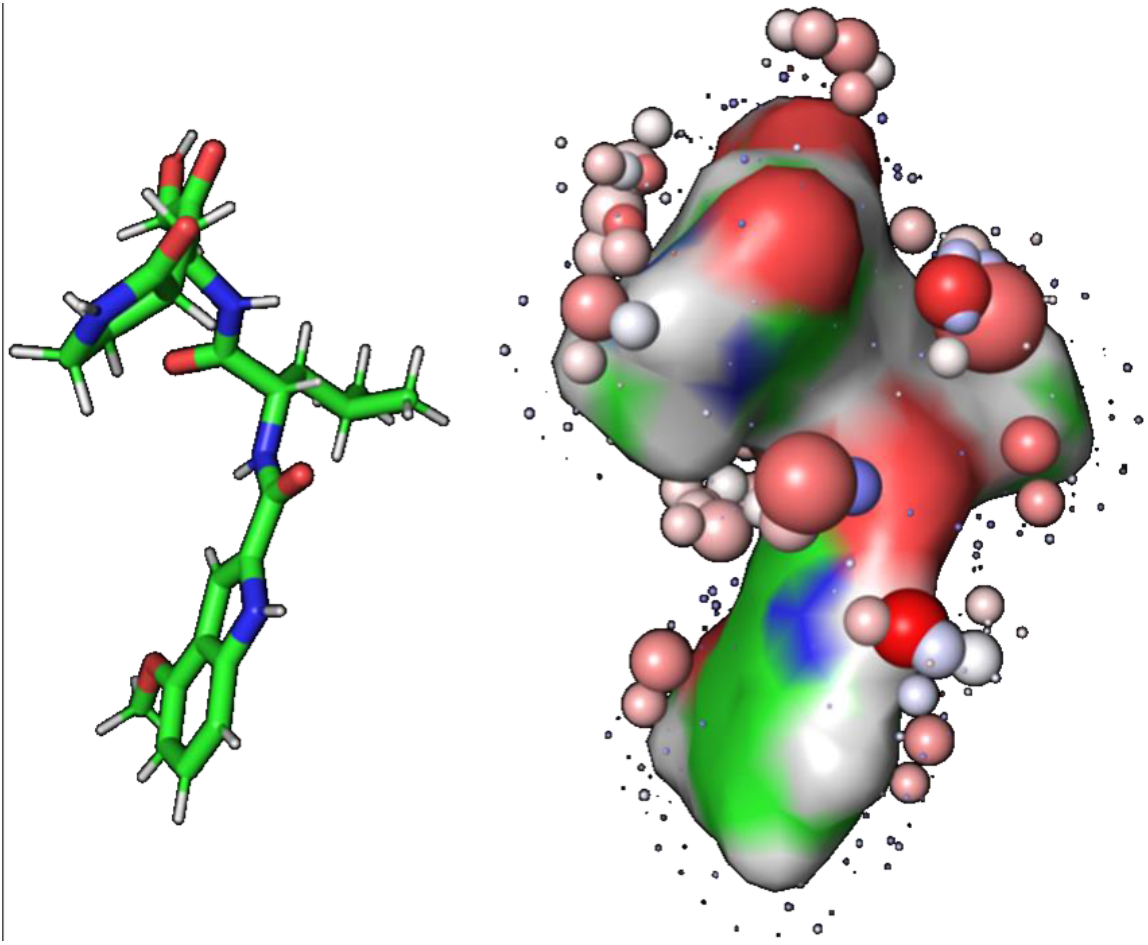
(Left) Non-covalent intra- and intermolecular energies are commonly interpreted on the basis of chemical structure and interatomic contacts. (Right) In reality, such energies are contributed largely by solute/state-specific solvation fields represented here by colored spheres (color-coding explained in Materials and methods) denoting H-bond depleted/low water occupancy (tiny spheres) and enriched/high water occupancy (large spheres) regions of the field. Solutes are driven toward their lowest free energy states corresponding to minimized H-bond depleted solvation and maximized H-bond enriched solvation content. Solubility and rearrangeability depend on a counter-balanced mixture of H-bond enriched and depleted solvation within a Goldilocks zone of total solvation free energy.

### Relaxation of the solvation field results from the replacement of interfacial H-bond depleted solvation by non-polar solute groups during intra- or inter-solute rearrangements

All non- covalent rearrangements are powered predominantly by the drive of each solvating water to achieve H-bond free energies ≤ that of bulk solvent (Figure 7). We proposed previously that “hydrophobicity” and “lipophilicity are misnomers for non-polar-non-polar interactions, which in practice, are less favorable than those between water and non-polar moieties.^3^ Sterically accessible solute states residing significantly below the Goldilocks zone of H-bond enriched solvation are non-rearrangeable, whereas those significantly above that of H-bond depleted solvation form slowly or not at all. H-bond enriched and depleted solvation “flavors” are comingled (non-randomly) and dynamically counter-balanced, as follows:

1) Overall solubility (neglecting dissolution) is promoted by H-bond enriched solvation residing largely on concave regions of solute surfaces.
2) Intra-/intermolecular rearrangeability (i.e., sensitivity to perturbations). Populated states reside within a Goldilocks zone of H-bond enriched versus depleted solvation. Such states form the fastest (i.e., have the lowest barriers or appropriate H-bond replacement groups), while achieving sufficient kinetic stability to perform their functions (e.g., enzyme-substrate complexes are sufficiently stable to turn over prior to substrate dissociation).

**Figure 7.**
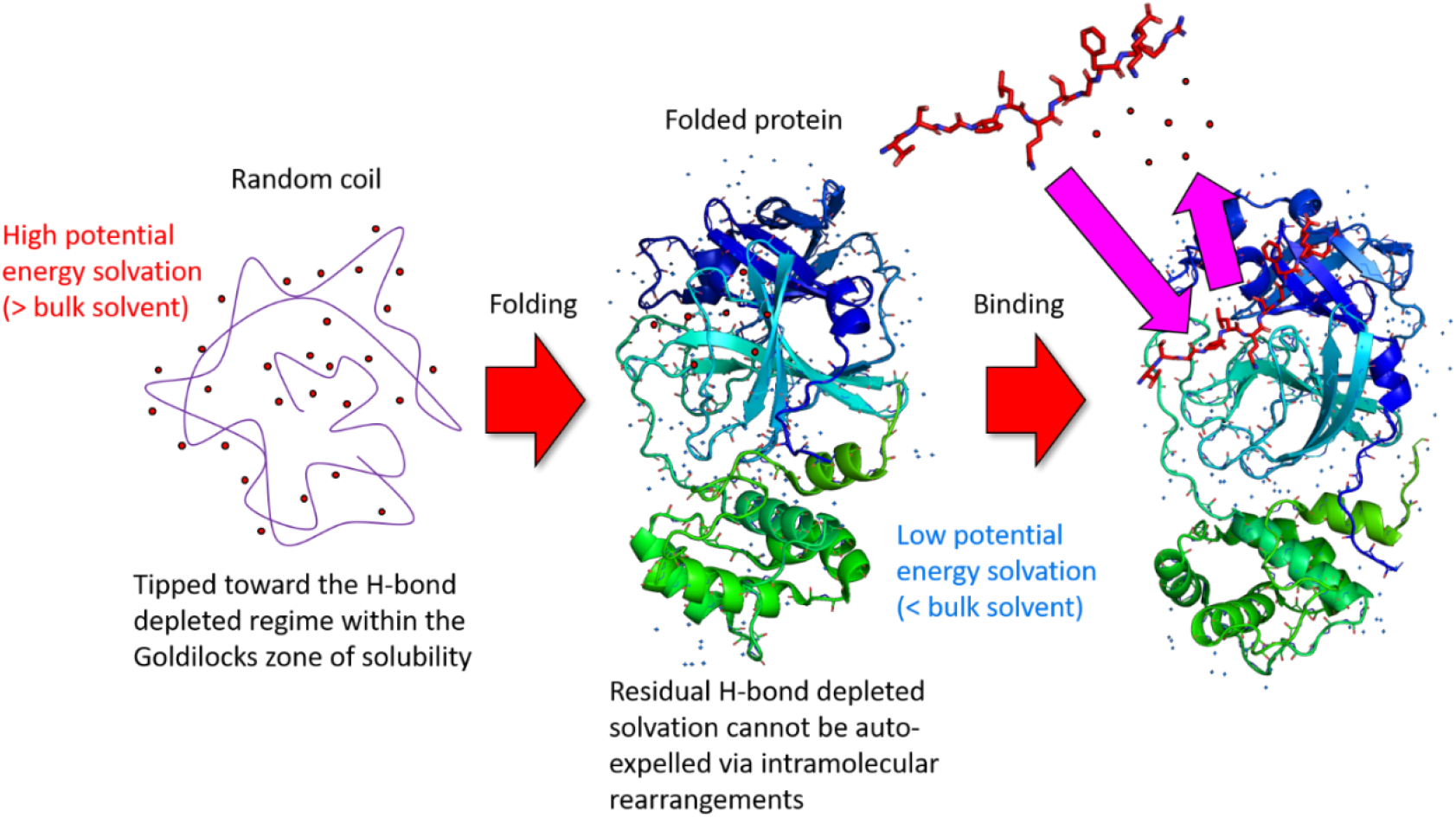
Cartoon depicting the auto-desolvation of high potential energy H-bond depleted solvation (denoted by red filled circles) (left) via folding of HMW and conformational rearrangements of LMW solutes (exemplified for the apo form of COVID M^pro^) (middle), and via non-covalent binding (right) (exemplified for the substrate-bound form of COVID M^pro^)^4^. Rearrangement-induced desolvation of H-bond enriched water (denoted by blue filled circles) typically incurs a loss of H-bond free energy, but is at best, a zero sum game of solute H-bond gains and water H-bond losses. Crystal waters correspond to bulk-like, H-bond enriched, or trapped solvation.

Continuous water exchanges between the solvation field and bulk solvent (Figure 8A), and possibly between H-bond depleted and enriched regions of the same field (Figure 8B), are driven by potential energy difference between these regions. Water transfer rates depend on the degree of enrichment or depletion of water-water H-bonds at each solute surface position, as follows:

1) Fully <u>exposed</u> water distributed around the external surfaces of LMW and HMW solutes, which is readily exchangeable with bulk solvent. Water occupancy is bulk-like at iso-bulk solvent H-bond positions (3.5 moderately strong H-bonds per water), high at H-bond enriched/energetically relaxed positions (3.5-4 strong H-bonds per water) corresponding to fast entry/slow exit, and low at H-bond depleted/energetically strained positions (< 3.5 weak H-bond per water) corresponding to slow entry/fast exit.
2) Partially exposed water occupying <u>buried</u> surface-connected channels that form transiently in HMW solutes, which exchanges slowly with bulk solvent due to impeded diffusion within the channels. Such channels appear in time-averaged molecular dynamics trajectories and some experimentally determined structures.
3) Water <u>trapped</u> within transient highly strained surface-disconnected buried cavities (“bubbles”) within HMW solutes, which is non-exchangeable with bulk solvent. Such volumes likewise appear in time-averaged MD trajectories and some experimentally determined structures.

**Figure 8.**
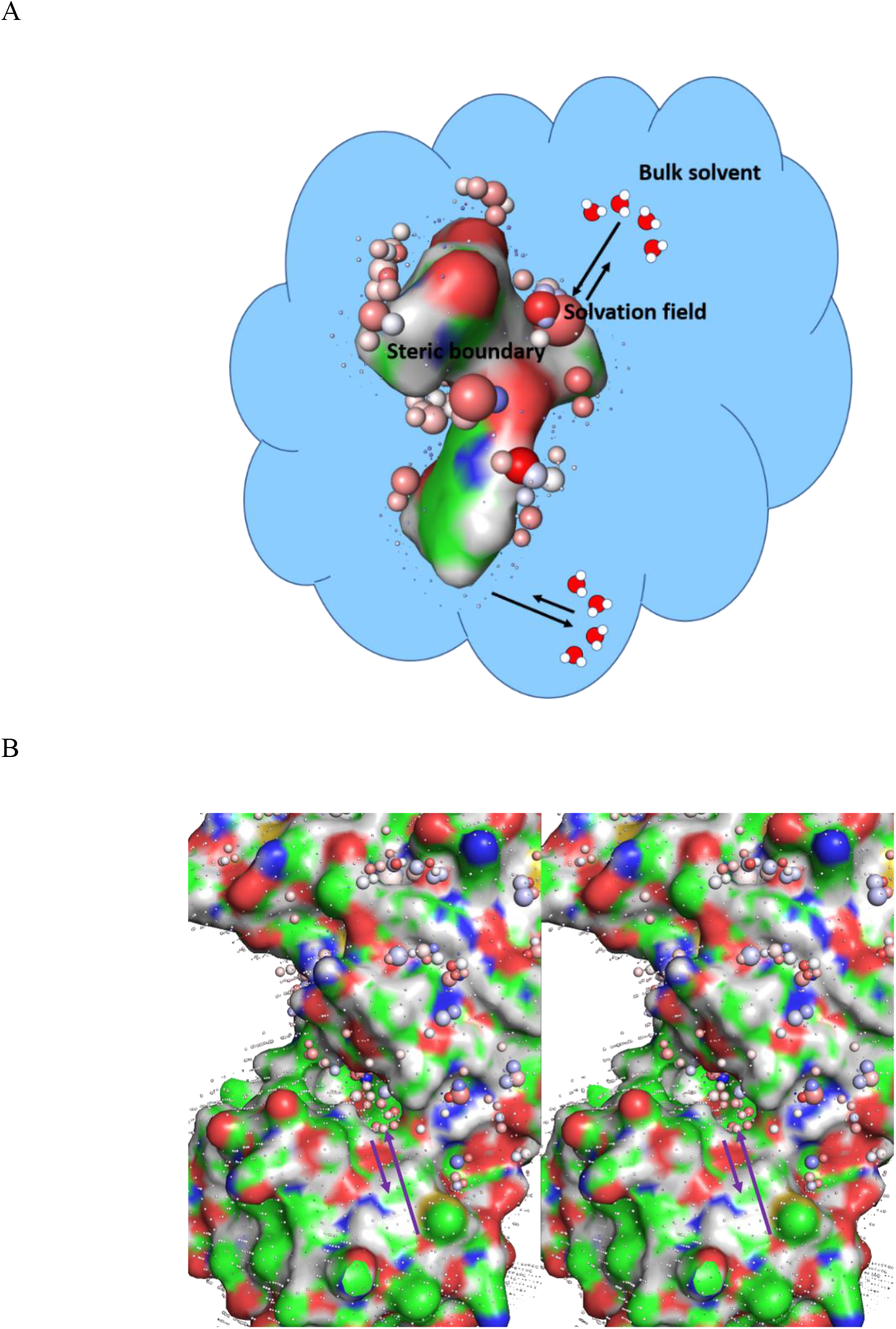
Cartoons showing the hypothetical exchanges between bulk solvent and H-bond enriched/high water occupancy (denoted by large spheres) and depleted/low water occupancy (denoted by small spheres) regions of the solvation field for LMW and HMW solutes (color-coded by atom type). The rates of water exchange to/from solvation and bulk solvent are governed by free energy barriers that are proportional to the loss of interfacial H-bond free energy relative to that of bulk solvent. (A) The solvation field of the COVID M^pro^ inhibitor PF-00835231 overlaid on a volume of bulk solvent (blue cloud). Fast water transfer <u>from</u> bulk solvent/slow transfer <u>to</u> bulk solvent occurs at H-bond enriched positions of the solvation field, whereas slow water transfer <u>from</u> bulk solvent/fast transfer <u>to</u> bulk solvent occurs at H-bond depleted positions. (B) Fast water transfer laterally across the surfaces of HMW solutes (exemplified here by COVID M^pro^)^4^ may likewise occur between H-bond depleted and H-bond enriched positions (stereo view of the M^pro^ surface color-coded by atom type).

We previously reported a qualitative first principles computational method called WATMD for predicting the distribution of H-bond depleted, enriched, and bulk-like solvation across internal and external solute surfaces relative to that of bulk solvent based on all-atom, explicit water MD simulations referred to in this work as “solvation dynamics” (SD) simulations to emphasize the focus on water exchanges versus molecular state transitions.^3, 4^ We used this approach to deduce the inhibition and functional state transitions of COVID M^pro^ based on a set of crystal structures in which the putative functional states were captured.^4^

### From equilibrium ΔG in vitro to non-equilibrium ΔG^‡^ in living cells

Under <u>equilibrium</u> conditions, the sterically-accessible configurations of HMW and LMW solutes are distributed <u>in a stationary fashion</u> in accordance with 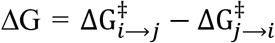) (Figure 9A). However, under <u>non-equilibrium</u> conditions in cells (the basis of signal-carrying molecular and ionic fluxes), the configurations are distributed <u>transiently</u> in proportion to the separate 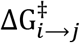 and 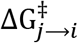 entry/exit barriers to/from each state (i.e., the intrinsic rates) (Figure 9B). The most populated states of a given species consist of those that build the fastest and/or decay the slowest <u>relative to the rates of buildup and decay of the species</u> itself (i.e., the extrinsic rates).^1^ We proposed more than a decade ago that:^23^

**Figure 9.**
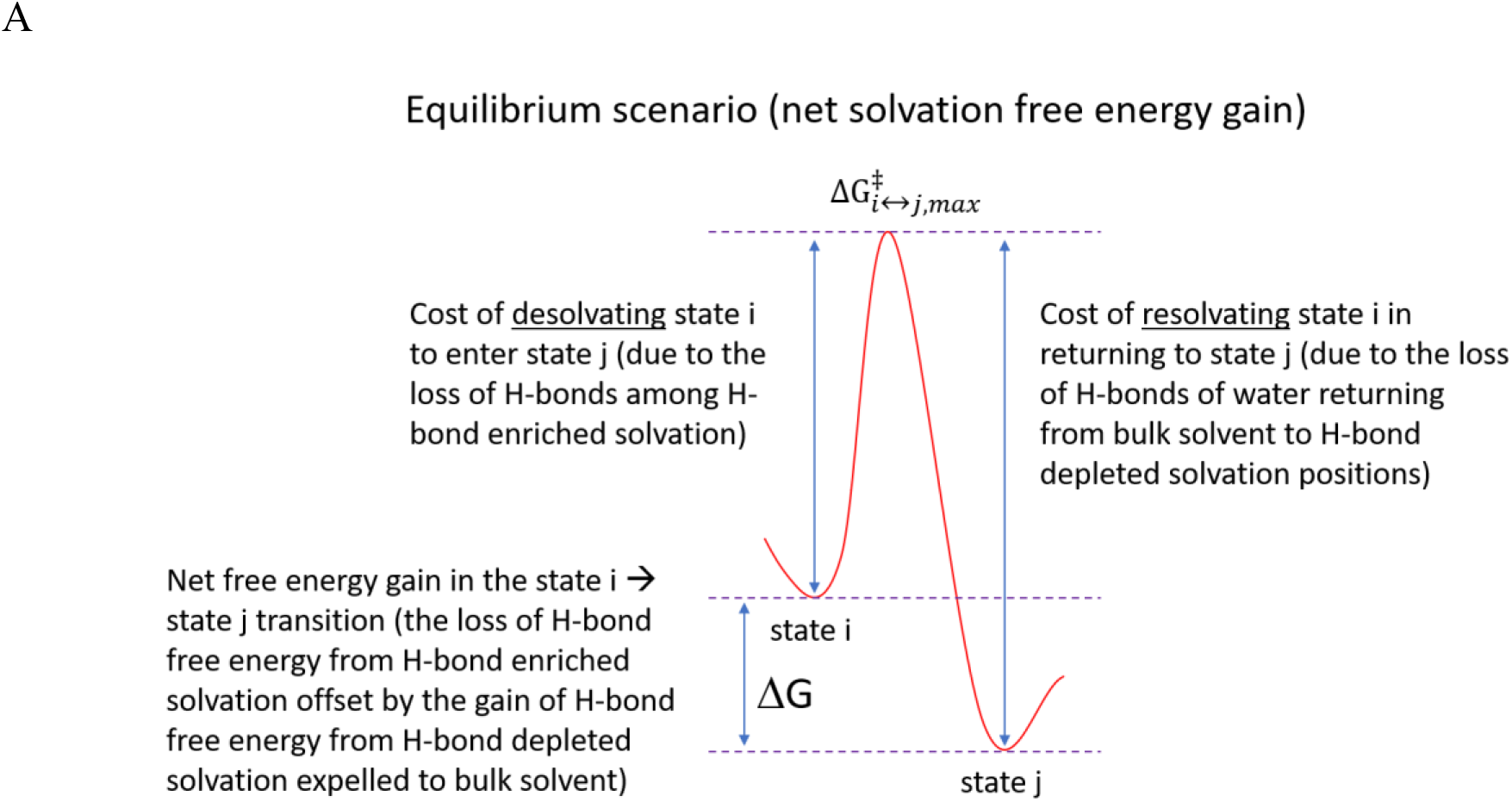
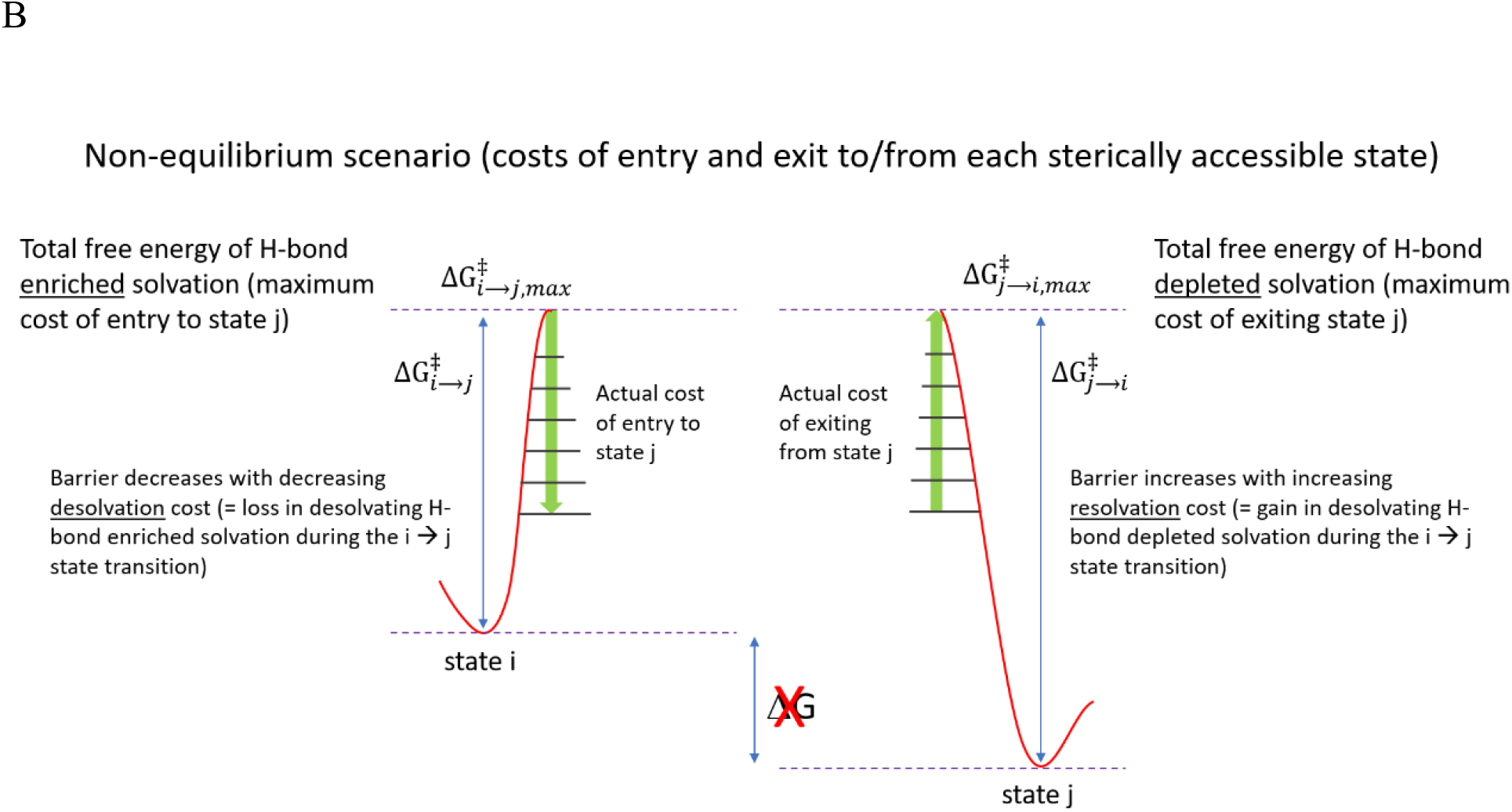
The sterically-accessible intra- and intermolecular structural states of a given solute equilibrate between states i and j via the expulsion of H-bond enriched and depleted solvation to bulk solvent upon dissolution of the solute at t = 0. (A) The equilibrium states at t = ∞ are distributed according to the <u>net</u> free energy of favorable H-bond enriched plus unfavorable depleted solvation in each state (ΔG_i↔j_). (B) Prior to equilibration, the states are distributed solely based on their rates of entry and exit (proportional to 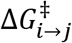 and 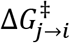, respectively), which in turn, are contributed principally by the cost of expelling H-bond enriched and depleted solvation to bulk solvent. As such, the cost of desolvating H-bond enriched solvation during the i → j state transition equates to the cost of accessing H-bond depleted solvation, which together, convey specificity (where the H-bond replacements are tuned for the cognate partners) and kinetic stability (where the rate of decay of state j is proportional to the cost of re-solvating H-bond depleted solvation positions that were desolvated during the i → j transition).

1) 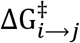 equates principally to the <u>desolvation</u> cost of H-bond enriched solvation, which decreases as the H-bonds of such water are replaced increasingly optimally by intra-solute H-bonds during the i → j state transition (Figure 9B). We define 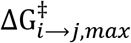 as the maximum cost of expelling H-bond enriched solvation in the complete absence of intra- or inter-solute water H-bond replacements, and 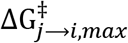 as the maximum cost of resolvating H-bond depleted positions during the i → j state transition. An additional cost is incurred during the i → j state transition when H-bond depleted solvation is generated de novo in state j due to the addition of solvent exposed non-polar groups and/or the loss of solvent exposed polar groups.
2) 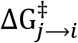 equates principally to the <u>resolvation</u> cost of H-bond depleted solvation positions from which water was expelled during entry to state j. We define 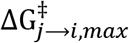 as the maximum possible cost of resolvating H-bond depleted solvation positions if all such positions were expelled during the i → j state transition. An additional cost is incurred in the j → i state transition when H-bond enriched solvation is generated de novo in state j.

The energetically accessible states of a given species exhibit low 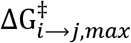 and/or high 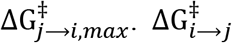 decreases as the H-bonds of H-bond enriched solvation are increasingly replaced by iso-energetic intra- or inter-solute H-bonds (the stringency of which grows with increasing magnitude of 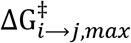). Net free energy <u>gains</u> due to such replacements are unlikely due to:

1) Entropy-enthalpy compensation, equating to the entropic cost accompanying large gains in H-bond enthalpy.
2) The typically lower free energy of optimal <u>water-solute</u> H-bonds that promotes H-bond enriched solvation compared with <u>intra- and -inter-solute</u> H-bonds (noting that increasing propensity of solute-solute H-bonds is mirrored by increasing propensity of water-solute H-bonds, resulting in a zero sum game at best).

<u>Intramolecular</u> transitions between states i and j (with the rate constant denoted as k_in_) are guided by the cost of desolvating H-bond enriched solvation in cis, as follows:

1) **k_in_ ≈ 0 (s^-^**^1^**) (collisions)**: high desolvation cost of H-bond enriched solvation within the rearrangement interface during entry to state j <u>in the absence of intra-solute replacements</u> for H-bonds lost during expulsion of H-bond enriched water to bulk solvent. The solvation fields across the interface effectively <u>bounce apart</u>.
2) **Maximal k_in_ (s^-^**^1^): low cost desolvation of H-bond enriched solvation within the rearrangement interface during entry to state j <u>in the presence of intra-solute replacements</u>.

The solvation fields across the interface <u>collapse</u> due to the expulsion of H-bond enriched solvating water to bulk solvent, thereby allowing access to H-bond depleted solvation positions by non-polar/weakly polar solute groups.

1) 3) **0 < k_in_ < maximal**: a range of desolvation costs of H-bond enriched solvation within the rearrangement interface during entry to state j associated with <u>partial</u> replacement of the H-bonds that were lost during water transfer to bulk solvent.

Intramolecular states are <u>kinetically stabilized</u> (with the rate constant denoted as k_out_) by H-bond depleted solvation, as follows (Figure 10A):

1) **Maximal k_out_ (s^-^**^1^): corresponding to no resolvation cost at H-bond depleted positions during exit from state j due to the lack of such positions or non-polar/weakly polar solute groups that are capable of replacing them at a low desolvation cost.
2) **0 < k_out_ < 1 (s^-^**^1^): corresponding to low-high resolvation costs at H-bond depleted positions during exit from state j.

Entry to state j is slowed by the de novo generation of H-bond <u>depleted</u> solvation in state j in excess of that in state i (which destabilizes state j), whereas exit from state j is slowed by the de novo generation of H-bond <u>enriched</u> solvation in state j in excess of that in state i (which stabilizes state j). In the absence of de novo H-bond enriched solvation, the lifetime of state j is determined principally by the resolvation cost of returning to state i or another state. <u>Intermolecular</u> state transitions are powered in a similar fashion as their intramolecular counterparts, except that desolvation of the interface occurs in <u>trans</u> (Figure 10B), and k_in_ and k_out_ are denoted as k_on_ and k_off_ (or k_1_ and k_-1_ in the case of enzymes), respectively. Intra- and intermolecular state transitions frequently occur in combination, reflecting the merger of the solvation fields of the binding partners (which are not simply the sum of the original fields of the partners due to non-linearity of the merging process).

**Figure 10.**
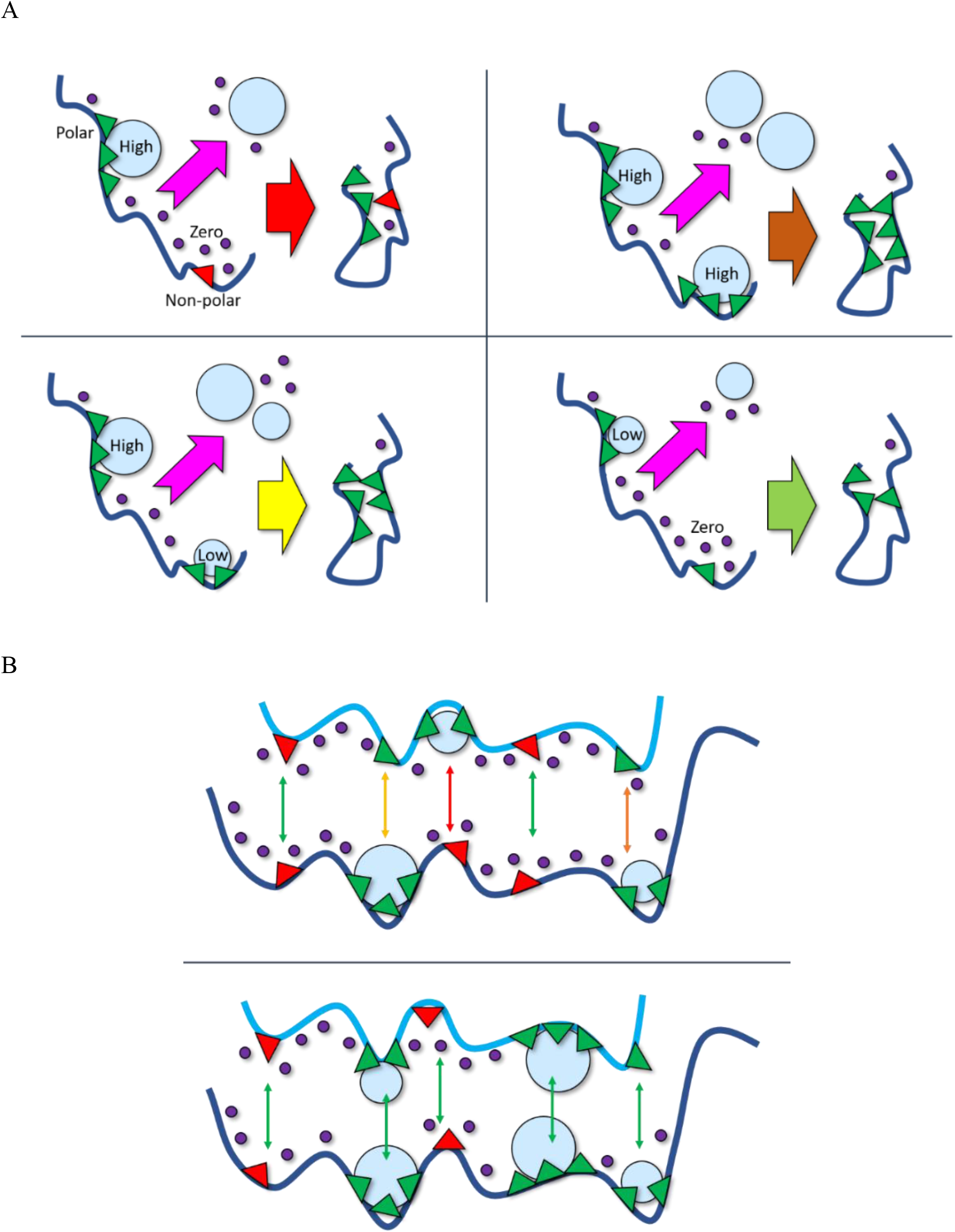
(A) The distribution of sterically accessible <u>intramolecular</u> states under both static equilibrium and dynamic non-equilibrium conditions is governed by the interfacial desolvation cost (enthalpic cost) and resolvation cost (enthalpic cost + entropic cost) incurred during entry to and exit from each state, respectively (noting that desolvation is entropically favorable and resolvation is entropically unfavorable, where entry and exit equate to the gain and loss of interfacial interatomic contacts, respectively). The rate of entry is proportional to the cost of <u>desolvating</u> H-bond <u>enriched</u> positions in cis, where slow entry equates to repulsion between unreplaced H-bonds of the interfacial solvating water. The rate of exit is proportional to the cost of <u>resolvating</u> H-bond <u>depleted</u> positions within the dissociating interface. We showed previously that little or no <u>resolvation</u> free energy cost corresponds to state lifetimes on the order of μs^27^. **H-bond enriched solvation thus serves as a “gatekeeper” for accessing H-bond depleted solvation**, relegating intermediate states to ultrashort lifetimes. The spectrum of desolvation costs is governed case-by-case by: 1) the magnitude of the excess H-bond free energy of H-bond enriched solvation relative to that of bulk solvent (denoted as Δ*G^enriched^*); and 2) the magnitude of the excess H-bond free energy of bulk solvent relative to that of H-bond depleted solvation (denoted as Δ*G^depleted^*). Four hypothetical scenarios are exemplified for a single pair-wise intramolecular desolvation reaction. Top left: a high Δ*G^enriched^* solvating water (light blue filled circle) at a <u>polar</u> solute position (green triangle) and two high Δ*G^depleted^* solvating water molecules (purple filled circles) at <u>non-polar</u> solute positions (red triangles). Top right: two high Δ*G^enriched^* solvating waters. Bottom left: high and low Δ*G^enriched^* solvating waters. Bottom right: a high Δ*G^enriched^* solvating water and a high Δ*G^depleted^* solvating water located at a weakly <u>polar</u> solute position. The positions of H-bond enriched solvation on polypeptide chains evolved to guide the folding process, whereas depleted positions evolved to stabilize folding-relevant sub-states along the folding pathway. (B) Same as A, except for the <u>intermolecular</u> case. Top: non-cognate partners collide, rather than associate, due to the high desolvation cost of H-bond enriched solvation within the binding interface. Bottom: cognate partners associate due to optimal solvation field complementarity, corresponding to lower desolvation cost of H-bond enriched solvation, and higher resolvation cost at H-bond depleted solute positions.

The most fundamental structure-free energy relationships therefore consist of structure-Δ*G^depleted^* and structure-Δ*G^enriched^*, rather than structure-ΔG relationships (which may be derived from the solvation free energy contributions under equilibrium conditions). The general mechanisms by which solvation free energy is transduced into the conformational distributions of HMW and LMW species, protein folding, and binding between HMW-LMW (including drug-target) and HMW- HMW species <u>under non-equilibrium conditions</u> are further explored and exemplified in the remainder of this work.

### Solvation-powered protein folding, structure, and function

Proteins fold and undergo specific intra- and intermolecular state transitions throughout their lifetimes. Non-linearity of multi-protein fluxes necessitates consideration of protein structure and function in a holistic fashion. However, for the sake of simplicity, we consider protein folding in isolation, as a non-linear stepwise desolvation-resolvation-powered process (Figure 11A). Folding begins near the upper boundary of the Goldilocks zone of H-bond depleted solvation free energy in the unfolded/random coil state (in which the protein is fully soluble), and proceeds toward the lower boundary throughout the folding process via the stepwise auto-expulsion of H-bond depleted solvation (nature’s form of energy minimization). Under non-equilibrium conditions, folding follows a strictly kinetic/dynamic (versus thermodynamic) pathway, in which the most populated states: 1) form the fastest; and 2) persist the longest. The timescale of protein synthesis/folding (or activation in the case of fast cellular responses), rearrangement (including binding), and deactivation or degradation is necessarily ≤ that of the corresponding function. The protein folding rate is governed by:

1) The rate of entry into each sub-state along the folding pathway, where the fastest rates (corresponding to the lowest cost of <u>desolvating</u> H-bond enriched solvation via optimal intra-protein replacement of the H-bonds of this water) reside along the evolutionarily- determined folding pathway.
2) The rate of exit from each substate along the folding pathway. Rate-limiting steps during folding are circumvented by short-lived sub-states. Exit rates slow as the random coil state condenses to secondary structure elements (SSEs) and tertiary structure powered by the cumulative replacement of H-bond depleted solvation by non-polar or weakly polar groups. The <u>resolvation</u> cost of total or partial unfolding or exit to a different folded substate determines the lifetimes of the folded states, respectively.

**Figure 11.**
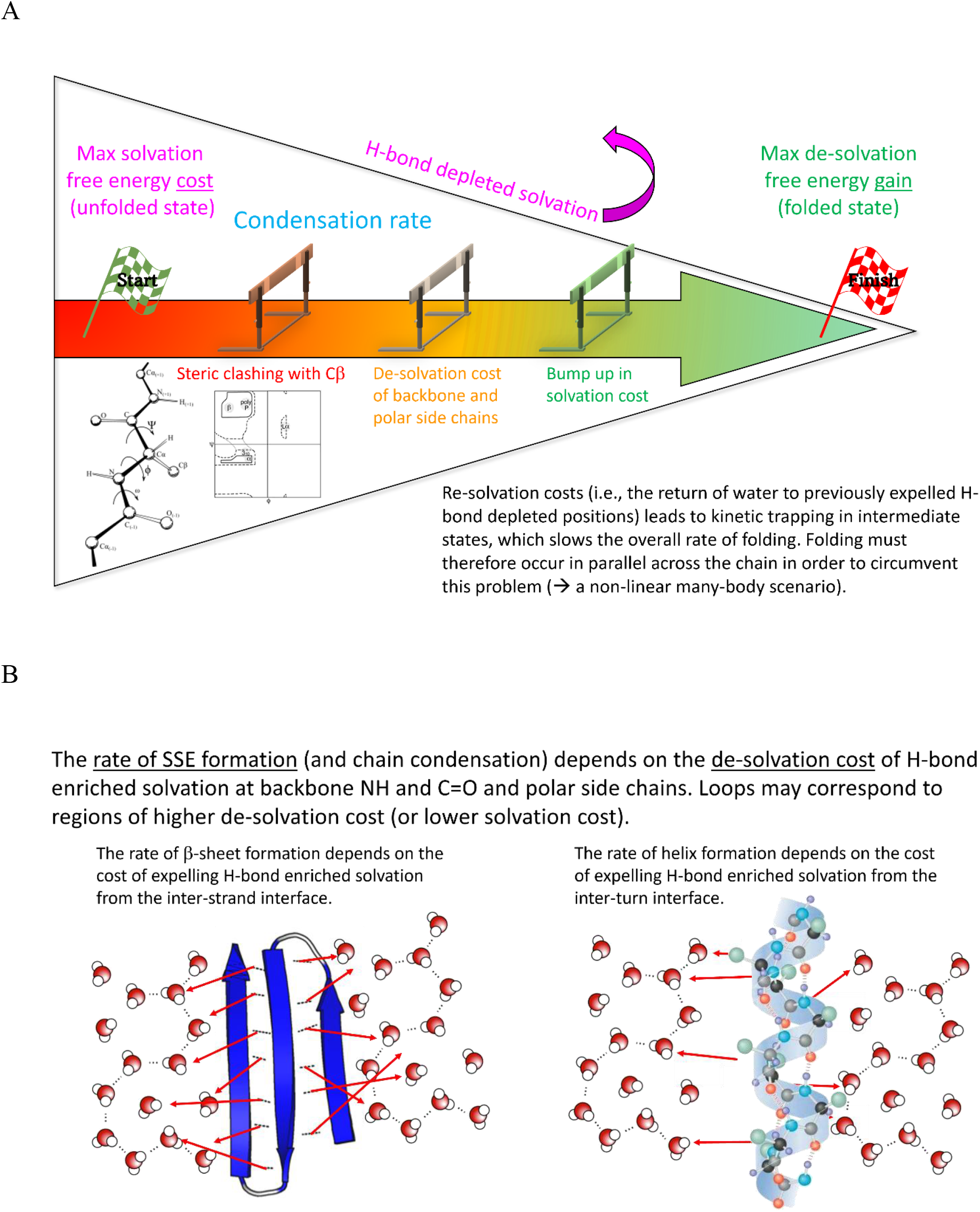
(A) Protein folding is driven by the step-wise desolvation of H-bond depleted solvation, from the maximal H-bond depleted solvation free energy in the unfolded state (i.e., release of the positive potential energy stored in such solvation) to the minimum in the folded state (noting that rearrangeability of the folded state depends on residual H-bond depleted solvation in the folded state within a Goldilocks zone of global stability). The folding pathway is guided by H-bond enriched solvation, in which the desolvation entry cost is lowest at each correct sub-state of the pathway compared with all other sterically accessible substates. (B) The fastest folding trajectories (necessarily corresponding to functionally folded proteins) consist of those in which H-bond enriched solvation is expelled by intra-protein, or in some cases intermolecular, H-bond replacements.

Folding is impeded by:

1) Excessively <u>favorable</u> solvation free energy in the unfolded state due to high <u>polar</u> amino acid content, which instead promotes extended, maximally solvent-exposed structural states.
2) Excessively <u>unfavorable</u> solvation in the unfolded state due to highly <u>non-polar</u> amino acid content, which promotes poor solubility, and in extreme cases aggregation/precipitation.
3) The absence of SSE formation in disordered regions (including loops and the N- and C-termini), in which auto-desolvation of H-bond depleted solvation is hampered due to:

a) Sterically constrained intra-protein H-bonding.
b) High desolvation cost of H-bond enriched solvation.

Disorder manifests as the protein’s attempt to alternately expel H-bond depleted solvation in multiple regions (commonly referred to as “conformational frustration”).

Rearrangeability of the <u>folded state</u> depends on a Goldilocks zone of counter-balanced <u>residual</u> H-bond depleted and enriched solvation, as follows:

1) Residual H-bond <u>depleted</u> solvation powers intra- and intermolecular rearrangements.
2) Residual H-bond <u>enriched</u> solvation stabilizes the folded state and counter-balances the destabilizing effect of H-bond depleted solvation, thereby limiting state transitions to those in which the H-bonds of expelled H-bond enriched solvation are replaced optimally by intra- or intermolecular H-bonds.

Intra-protein H-bonds present in experimentally observed protein structures represent the final cumulative replacements of H-bond enriched solvation desolvated throughout the folding process. **As such, intra-protein H-bonds correspond to positions at which H-bond enriched solvation “gatekeepers” controlling co-localization of the participating donors/acceptors have been optimally replaced** (Figure 11B), but do not contribute directly to the stability or persistence of the folded structure. It is further apparent that each intramolecular interface from which H-bond depleted solvation was expelled during the folding process decays as the interface resolvates (the lifetime of which depends on local resolvation costs), and reforms at a rate governed by the local desolvation costs. This putatively coincides with the transient formation of buried solvation (apparent in many of our time-averaged SD trajectories) that is capable of storing and releasing potential energy (as we demonstrated previously for COVID M^pro^,^4^ and as we demonstrate for cereblon below).

Sterically allowed ϕ-ψ backbone torsion angles, which are constrained largely by their common C_β_-backbone interactions, are similar for all <u>isolated</u> amino acids except Gly and Pro. Preferences for particular ϕ-ψ angles in polypeptide chains are therefore determined largely by side chain-side chain and side chain-backbone interactions between flanking and remote residues, as follows:

1) Side chain-specific and backbone solvation fields in the unfolded state.
2) SSE-specific solvation fields.
3) Side chain-side chain and side chain-backbone packing (i.e., the extent of packing- mediated backbone and side chain desolvation).
4) Side chain-SSE, backbone-SSE, and SSE-SSE packing (i.e., the extent of packing- mediated desolvation of secondary structure elements).

SSE formation is the means by which local polypeptide regions are condensed into compact substructures, analogous to the winding of a garden hose around a reel (helices) or the folding of a firehose (parallel and anti-parallel β-sheets) (Figure 12A). α-helices and β-sheets undergo further packing to form helical bundles and α/β structures (Figure 12B).

**Figure 12.**
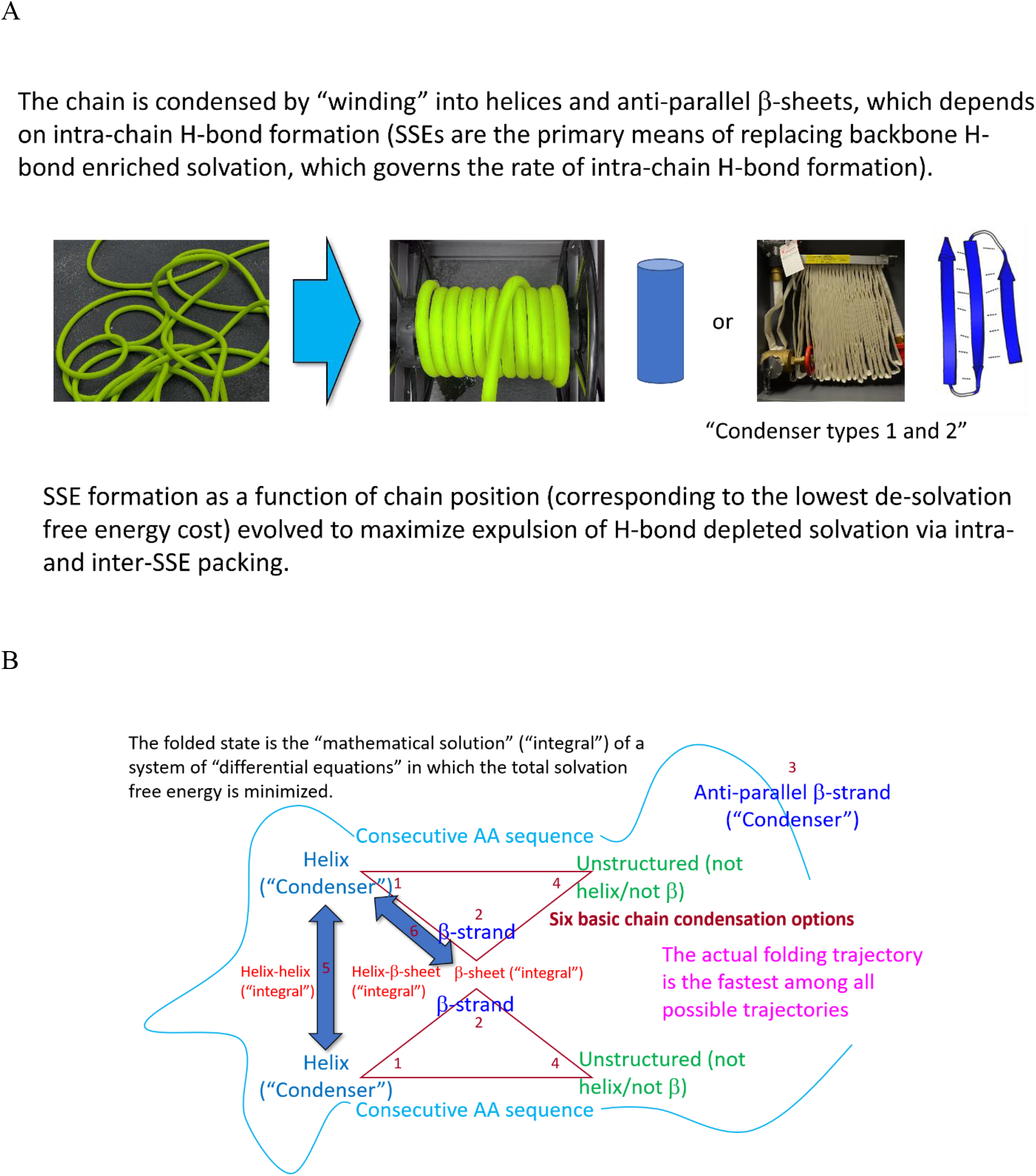
(A) Polypeptide chains condense into compact substructures via helix and β-sheet formation. (B) The chain further condenses via packing between SSEs.

Each folding step depends on the <u>mutual</u> relationship between the transient protein geometry and corresponding solvation field and many-body interactions between the remote quasi-independent parts of the chain (e.g., β-sheet, α/β, and helical bundle formation), resulting in non-linearity of the folding process. Whereas isolated SSEs can be loosely predicted based on sequence-structure relationship models, first principles prediction of the SSEs within tertiary structures is hampered by the non-linearity of their formation (Figure 13) (noting that data models like AlphaFold^28^ are useful for predicting <u>folded structures</u> rather than <u>folding pathways</u> per se).

**Figure 13.**
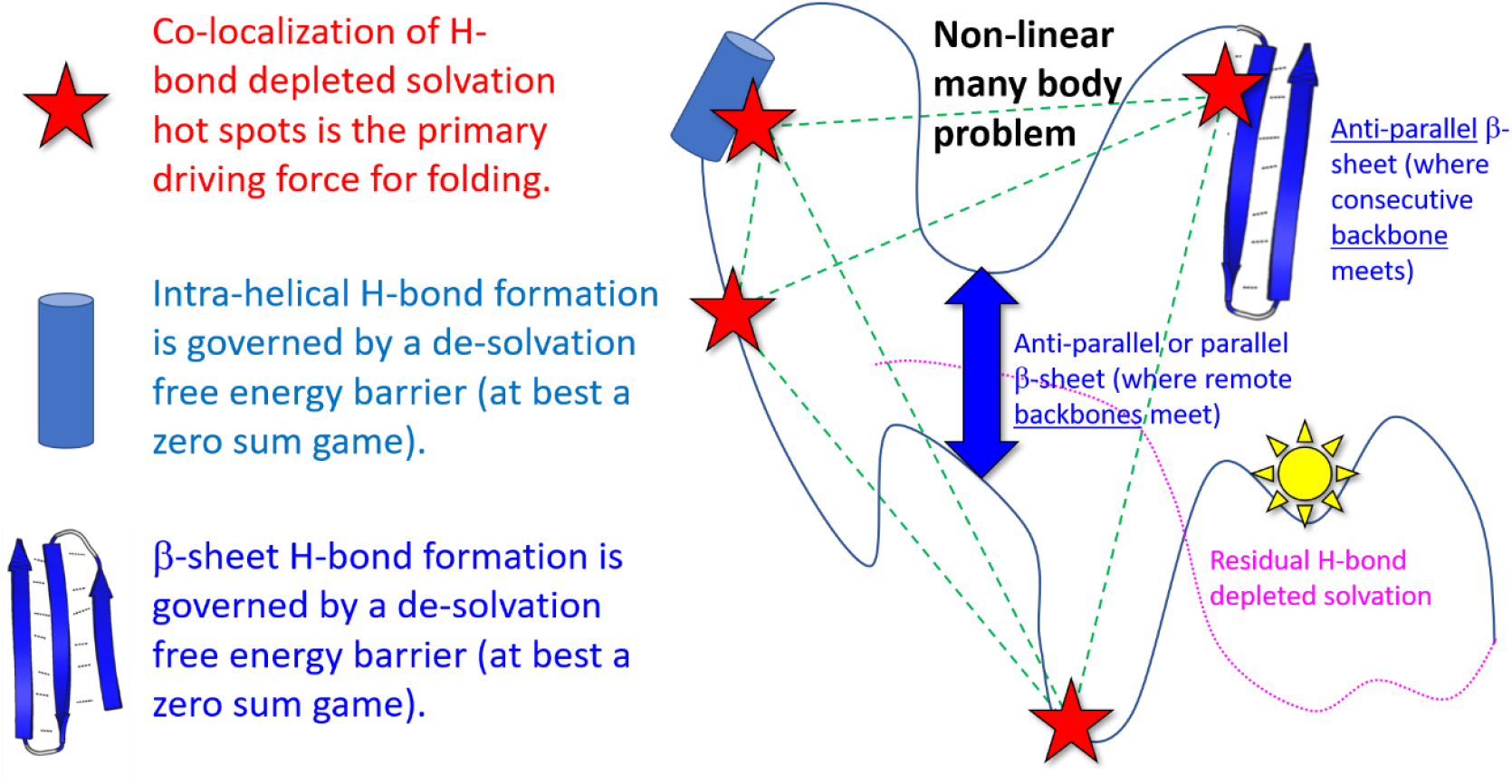
Each substructural element of a protein undergoing folding depends on all other elements, and vice versa, consistent with NLD behavior (i.e., the folded state of a protein is > the Σ(states of the constituent residues)).

Protein folding and the state transitions of folded proteins are both powered and facilitated by solvation, as follows:

1) Protein fluxes operate in a non-equilibrium source (synthesis/translocation)-sink (degradation translocation) fashion, in which the distribution of populated states is tipped transiently to one or more specific states in response to perturbations/signals.
2) States build and decay on the basis of local packing and unpacking of SSEs and/or unstructured regions (e.g., surface loops or buried regions) according to their desolvation and resolvation costs, respectively.
3) The external surfaces of proteins are solvated, as are rearrangeable <u>buried</u> SSEs and non-structured elements (NSEs), the latter in the form of transient internal water-filled cavities/pockets/channels (which we refer to as “internal solvation”) (Figure 14).
4) Internal solvation may be trapped (i.e., non-exchangeable with bulk solvent or other buried cavities), impeded (i.e., exchange more slowly with bulk solvent than external solvation), or freely exchangeable (i.e., exchange at a similar rate with bulk solvent). Internal solvation exists within a Goldilocks zone of enthalpic and entropic stability (i.e., overtly unstable and stable cavities/pockets/channels do not form in the first place, and not rearrange, respectively). The upper boundary of the Goldilocks zone consists of fully trapped water that is partially stabilized by water-water and water-protein H-bonds (at an entropic cost). The lower boundary consists of water that exchanges freely with bulk solvent via surface openings.
5) Internal NSEs rearrange transiently into water-filled cavities/pockets/channels, resulting in trapping or expulsion of the resident solvation to bulk solvent. The rearranged structural elements are necessarily resolvated by newly formed cavities/channels. **As such, buried water-filled cavities/pockets/channels are signatures of rearrangeable structural elements**.
6) Protein states build and decay at rates governed by the local interfacial resolvation and desolvation costs of their rearrangeable internal structural elements (where the rate of buildup exceeds the rate of decay and vice versa, respectively).
7) Desolvation of deep concave pockets via ligand association may depend on buried water channels that connect the binding interface to bulk solvent, or between the ligand and binding site lumen when the ligand volume is sufficiently less than the pocket volume.

**Figure 14.**
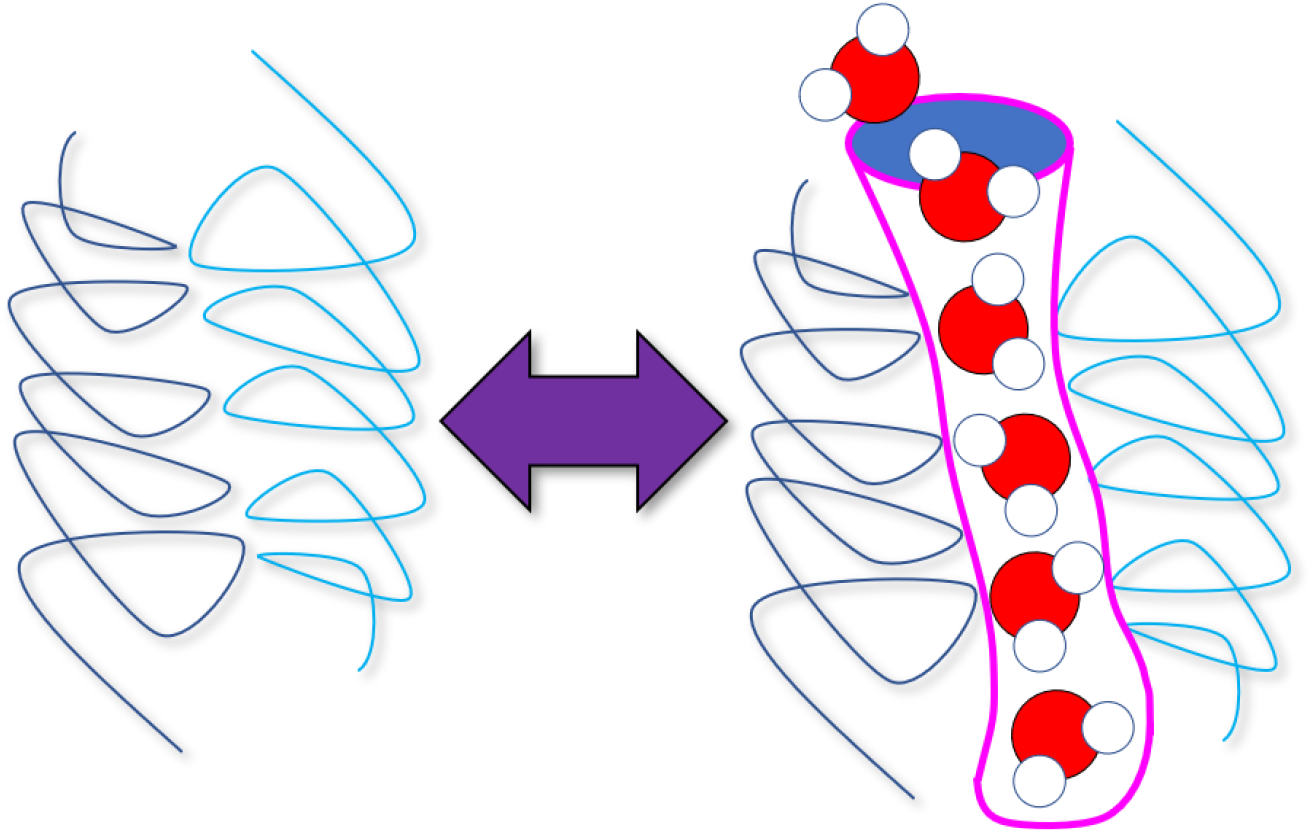
Intra-protein rearrangements are powered by the release of high energy solvation comprised of trapped/impeded water occupying transiently formed buried channels and cavities that we refer to as “internal solvation”.^4^

### The solvation field is the target of many post-translational modifications

The solvation fields of many solutes are modulated by post-transitional modifications (e.g., phosphorylation/dephosphorylation, lysine acetylation/deacetylation, glycosylation), in which the solvation free energy and desolvation costs of the targeted solute groups are increased or decreased to varying degrees. Electrostatic interactions between charged groups that are separated spatially by ≥ the Debye length are heavily screened by solvating water, the reason that co-localized positively charged side chains are commonly observed on solvent exposed protein surfaces. Rather, charge-charge (as well as charge-neutral) interactions are governed principally by the additional cost of desolvating the stronger water-solute H-bonds arising in such cases, compared with that of neutral groups. Extended conformations of targeted groups and the local substructures to which they are attached result from increased H-bond propensity and solvent exposure (affected substructures are effectively dragged into bulk solvent). Furthermore, the overall molecular density/radius of gyration is putatively governed by counter-balancing between the drive toward the maximal resolvation cost stabilizing the folded state (an inwardly-directed force) and maximal solvation of charged and polar surface groups (an outwardly-directed force). As such, the energy balance to/from folded versus extended solvent exposed conformations of the affected group(s) (e.g., the activation loop of protein kinases in the unphosphorylated and phosphorylated states) is tipped toward one state or the other via reversible post-translational modifications, contrary to the widely assumed electrostatic switching mechanism.

### Overview of E3 ligase-mediated protein degradation

Protein degradation serves as the ultimate sink for protein fluxes (the “terminal Yang”), many of which are degraded via the proteasomal pathway. Proteasome-mediated protein degradation in humans is initiated by poly-ubiquitylation of targeted proteins by a broad family of E3 ligases, including cullin RING ubiquitin complexes assembled from the family members of cullin (Cul), RING-Box (RBX), adapter (e.g., DDB), and substrate recruitment (e.g., DCAF) subunits.^29^ Cereblon, which consists of separate N- and C-terminal (CULT) domains, serves as the DCAF subunit of the CUL4A-DDB1-RBX1-CRBN E3 ligase.^29^ Targeted substrates of this ligase bind exclusively to binary complexes comprised of the CULT domain of cereblon and LMW compounds (commonly referred to as “degraders”). The function(s), native degrader(s) and substrate(s) of the CUL4A-DDB1-RBX1/CRBN E3 ligase are poorly understood. Glutamine synthase has been proposed as a native substrate, and uridine as a native degrader (suggesting a possible role of this ligase in repairing DNA containing mis-incorporated uracil bases).^30–32^ Cereblon is also the target of drugs (e.g., thalidomide and analogs thereof) that recruit specific non-native (“neo”) substrates (e.g., CK1α and BRD4) that are poly-ubiquitylated by the CUL4A- DDB1-RBX1-CRBN E3 ligase and subsequently degraded (commonly referred to as “targeted protein degradation” (TPD)). Neo substrates are recruited by two classes of LMW compounds referred to in this work as Type I degraders (widely known as “glues” or IMiDs) and Type II degraders (widely known as “bifunctionals” containing cereblon- and substrate-binding ends). Type I degraders bind in an order-dependent fashion, first to cereblon, followed by substrate binding to the binary cereblon-degrader complex, whereas Type II degraders form binary complexes with cereblon and their substrate targets independently, such that only the fraction of degrader bound to both proteins simultaneously (one per end) is catalytically viable and the remaining fraction is dead. Neo substrates can be classified, as:

1) Those mimicking the native substrate-cereblon footprint (Figures 15A and B), which includes the degrader pocket of cereblon (denoted in this work as pocket L) and a β-turn on the target protein (recruited by Type I degraders).
2) Those exhibiting smaller footprints that bypass the degrader pocket (Figures 15C and D) (recruited by Type II degraders). We hypothesize that the k_-1_ (the dissociation rate constant) of neo substrates is slowed by Type II degraders, which otherwise lack sufficient H-bond depleted and/or de novo H-bond enriched solvation residing on the substrate binding surface.

**Figure 15.**
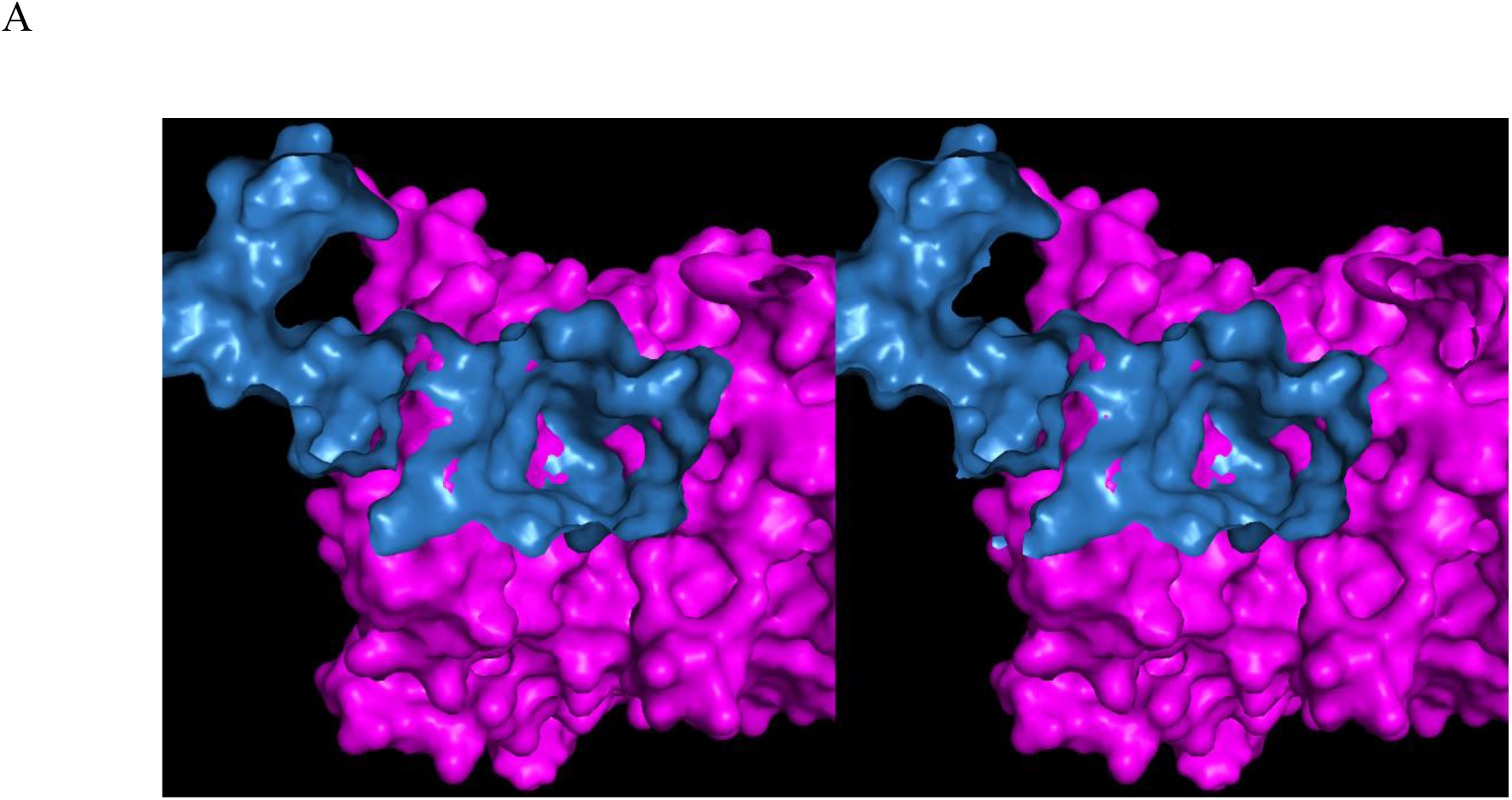
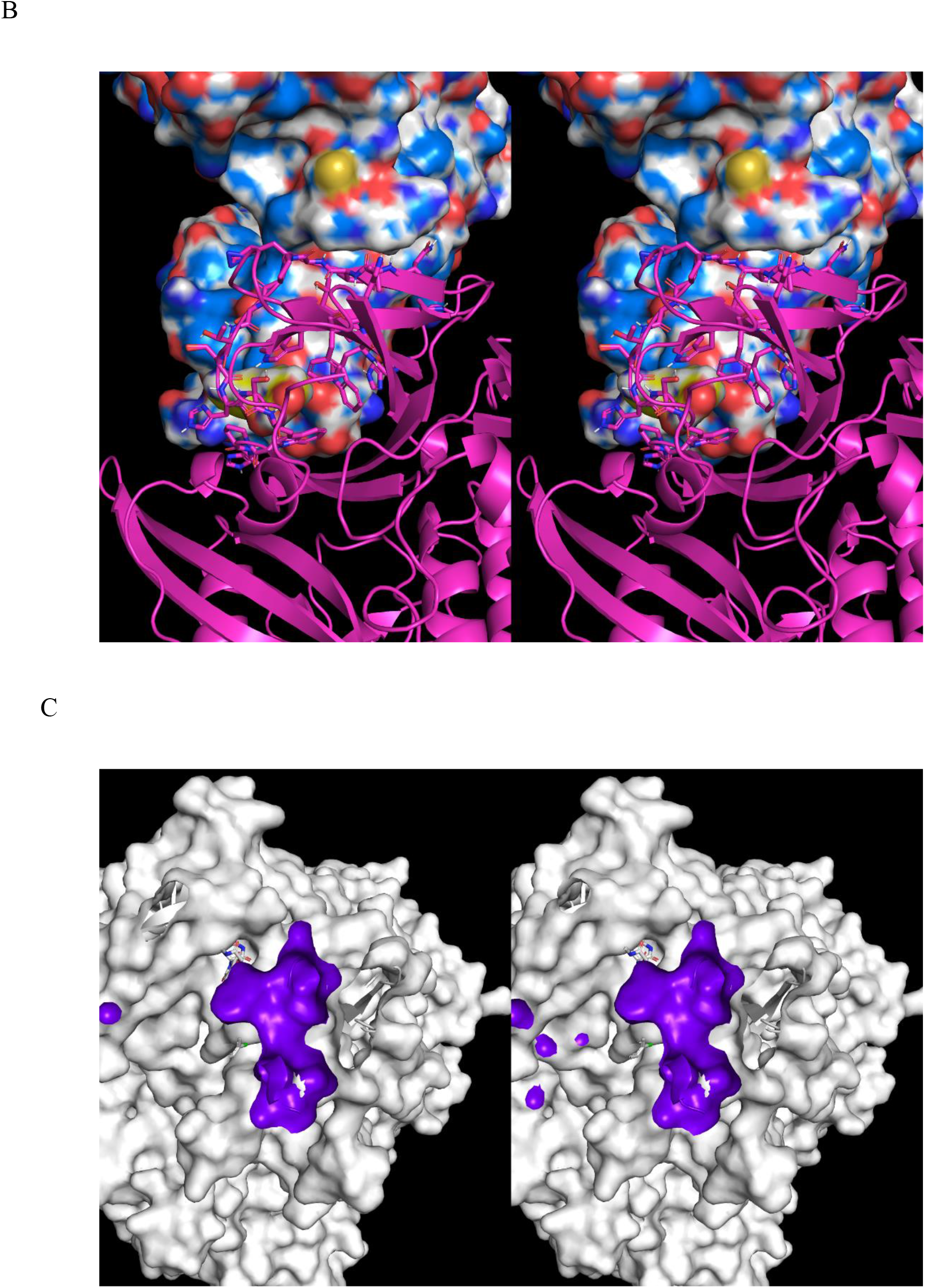
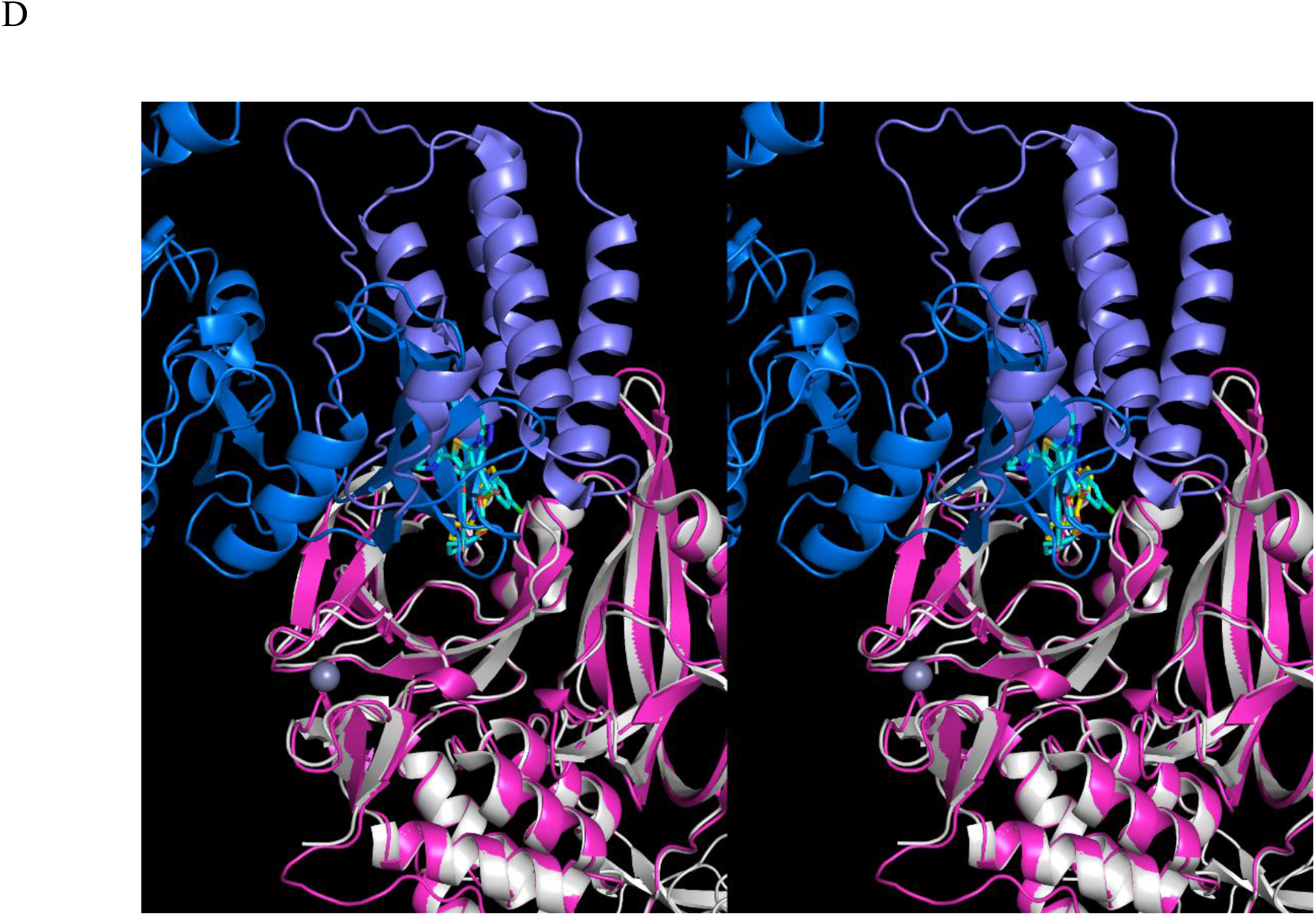
(A) Stereo view of the approximate contact footprint of CK1α (blue) on cereblon (magenta) in the ternary cereblon-lenalidomide-CK1α complex. (B) The saddle-shaped CK1α- cereblon interface in the ternary structure looking from the bottom of the degrader pocket in cereblon (magenta ribbon) toward CK1α (blue surface). The β-hairpin of CK1α fits within the circular β-sheet containing the degrader binding site (visible as a cluster of aromatic side chains). (C) Stereo view of the approximate contact footprint of BRD4 on cereblon in 6BN7, which is far smaller than that of CK1α. Unlike CK1α, pocket L resides outside of the BRD4-cereblon interface. (D) Stereo view of CK1α-cereblon from the ternary complex (blue and magenta, respectively) overlaid on BRD4-cereblon (purple and white, respectively) (PDB code = 6BN7). BRD4 and CK1α bind adjacent to and opposite to the N-terminal domain of cereblon, respectively. The Type II bifunctional degrader dBET23 plays a different role than the Type I degrader lenalidomide in recruiting substrates. Type II degraders are predicted in this work to speed the substrate k_1_ and slow the k_-1_ by augmenting the small cereblon-substrate footprint, whereas Type I degraders are predicted to primarily speed the substrate k_1_.

TPD is a proven clinical strategy, and the structural features governing recruitment are known for several target proteins.^33–38^ However, the mechanistic, kinetic, and dynamic aspects of recruitment and degradation in the native cellular context addressed in this work are poorly understood, and typically treated using equilibrium models^39^ that inconsistent with time-dependent substrate degradation under native conditions.

## Materials and methods

We used a top-down approach to study TPD using multi-scale Biodynamics principles. First, we explored the generic TPD pathways for Type I and II degraders using ordinary differential equation-based models of the constituent multi-flux systems (noting that we used these models to deduce the optimal kinetic relationships among the species but did not simulate the time-dependent behaviors of the pathways). We then used WATMD-calculated solvation fields to deduce the solvation-powered structural dynamics governing non-equilibrium binary cereblon-degrader and ternary cereblon-degrader-substrate complex formation, including large-scale rotation of the C-terminal (CULT) domain relative to the N-terminal domain observed in 5FQD versus 6H0F (which we speculate facilitates access of the E2-ubiquitin (Ub) subunit to the terminal Ub of the growing poly-Ub chain during processive ubiquitylation). We calculated the solvation fields of cereblon (focused on the degrader and substrate binding sites), LMW degraders, and the neo substrate CK1α (focused on the cereblon binding site) separately and in combination using the structures summarized below and in Tables 1 and 2:

1) Cereblon +/- DDB1 (denoted as structures A_closed and A_closed’, respectively).
2) Cereblon-CK1α, from which lenalidomide was removed (denoted as structure B).
3) Cereblon-lenalidomide, from which CK1α was removed (denoted as structure C).
4) Cereblon-lenalidomide-CK1α (the original 5FQD structure, denoted as structure D).
5) CK1α, from which cereblon and lenalidomide were removed (denoted as structure E).
6) Cereblon + DDB1 (derived from 6H0F), the C-terminal (CULT) domain of which is rotated relative to the N-terminal domain (denoted as structure A_open).
7) Mouse cereblon, in which the N-terminal domain is absent.

**Table 1.** The HMW solutes (proteins) included in our study (noting that 5FQD is the human ortholog of the protein complex).

| Name | PDB code | Reference code |
| --- | --- | --- |
| Apo-like* cereblon (closed, -DDB1) | Time-averaged (from 5FQD<sup>40</sup>) | A_closed |
| Apo-like cereblon (closed, +DDB1) | Time-averaged (from 5FQD) | A_closed' |
| Apo-like cereblon (closed, +DDB1 with restraints applied in SD simulation) | Time-averaged (from 5FQD) | A_closed'' |
| Apo-like cereblon (open, +DDB1) | Time-averaged (from 6H0F<sup>33</sup>) | A_open |
| Cereblon-CK1α (closed) | Time-averaged (from 5FQD) | B |
| Cereblon-lenalidomide (closed, -DDB1) | Time-averaged (from 5FQD) | C |
| Cereblon-lenalidomide (closed, +DDB1) | Time-averaged (from 5FQD) | C' |
| Cereblon-lenalidomide (closed, -DDB1, -N-terminal peptide) | Time-averaged (from 5FQD) | C'' |
| Cereblon-lenalidomide-CK1α (closed) | 5FQD (chains B and C) | D |
| CK1α | Time-averaged (from 5FQD) | E |
| Cereblon-lenalidomide-CK1α (closed) | Time-averaged (from 5FQD) | F |
| Cereblon-pomalidomide-IKZF1 (open) | 6H0F | G |
| Apo cereblon CULT (mouse)<sup>#</sup> | Time-averaged (from 3WX2<sup>35</sup>) | H |
| Lenalidomide-cereblon CULT (Ms-CI4) | 4V30<sup>30</sup> | I |
| BRD4-cereblon (closed) | 6BN7<sup>41</sup> | J |
| Uridine-cereblon docking model | Time-averaged (from 5FQD) | K |

\*Apo-like cereblon refers to the substrate and degrader unbounded state of the protein modeled from 5FQD. <sup>#</sup>Mouse cereblon (which lacks the N-terminal domain present in the human protein) is the only experimentally determined apo cereblon structure reported as of this writing.

**Table 2.** The LMW solutes included in our study.

| Name | Chemical structure | PDB code |
| --- | --- | --- |
| Lenalidomide | https://pubchem.ncbi.nlm.nih.gov/compound/Lenalidomide | Extracted from 5FQD |
| Thalidomide | https://pubchem.ncbi.nlm.nih.gov/compound/Thalidomide | Extracted from 4CI1<sup>42</sup> |
| Uridine | https://pubchem.ncbi.nlm.nih.gov/compound/Uridine | Modeled |

The software and methods are summarized briefly here, and in detail in our previous work.^4^ The simulations were performed using AMBER 20 (University of California, San Francisco) in explicit TIP3P water to simulate the solvation dynamics (which we refer to as “SD simulations” due to the focus on water) of both the LMW and HMW structures and complexes. We used WATMD V9 was to analyze the dynamic water exchanges between bulk solvent and a three-dimensional grid of 1 Å^3^ voxels proximal to the solute surfaces. The accumulated occupancy of water oxygen (O) and hydrogen (H) atoms within each voxel was calculated over the last 10 ns of a 100 ns trajectory (Figure 16). Such occupancies are governed by the water exchange rates to/from each voxel, including water donor versus acceptor preferences, as described in the Introduction. H-bond <u>depleted</u> solvation manifests as ultra-low occupancy voxels (denoted as ULOVs) represented by tiny white spheres exhibiting little or no preference for water H or O. Such solvation resides in the left tail of the Gaussian distribution of water occupancies reported in our previous work.^4^ H-bond <u>enriched</u> solvation manifests as high occupancy voxels residing in the right tail of the Gaussian distribution (denoted as HOVs) represented by larger spheres (the radii of which are proportional to water occupancy) exhibiting a range of preferences for water H or O, including:

1) High preference for H (bright blue).
2) Moderate preference for H (light blue/purple).
3) No preference for either (white).
4) Moderate preference for O (pink/light red).
5) High preference for O (bright red).

**Figure 16.**
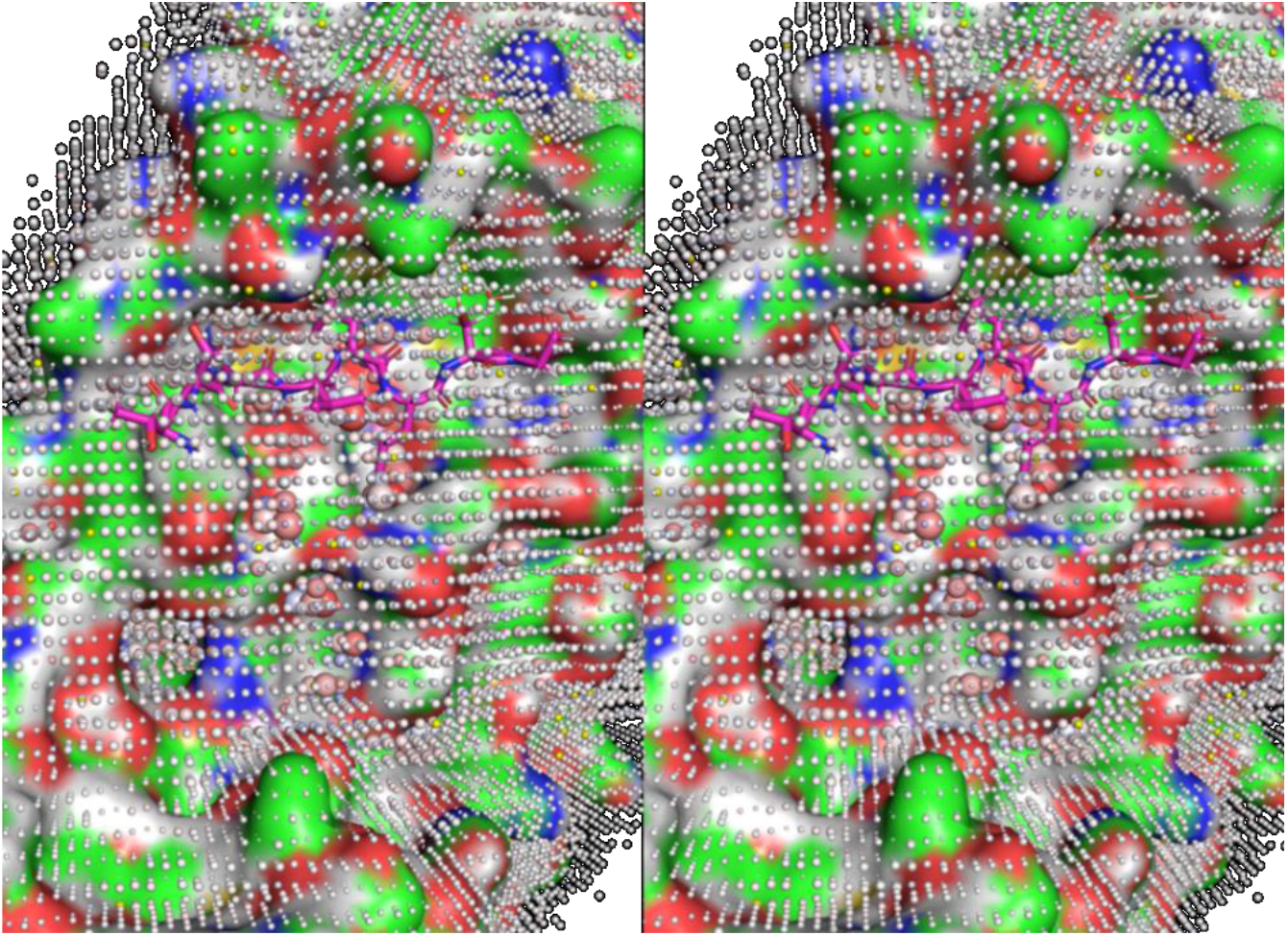
Stereo view of predicted water occupancies within a three-dimensional grid of voxels around a solute surface exemplified for COVID M^pro^ (surface colored by atom type)^4^ generated using PyMol 2.4.2 (Schrodinger, LLC) (noting that all WATMD results reported in this work were generated using PyMol). Bulk-like solvation (centered around the mean of the Gaussian curves shown in reference ^4^) is represented by small white spheres, H-bond enriched solvation by larger spheres, and H-bond depleted solvation by tiny spheres (non-discernable in this figure). Bulk solvent, residing above the outermost voxel layer shown, was omitted from the drawing (and all other drawings in this work) for the sake of clarity. The spheres are color-coded according to the preference of the corresponding voxel for water H or O in shades of blue and red, respectively (see text). Water occupancy and H versus O preference within each voxel are determined by the local the solute environment.

The solvation fields were superimposed on the corresponding time-averaged solute structures (generated by WATMD) and visualized using PyMol 2.4.2 (Schrodinger, LLC).

### HMW SD simulation protocol

Briefly, the structures listed in Table 1 were prepared using the default settings of the PPREP tool in Maestro 12.7 (Schrodinger, LLC). The structures were then solvated in explicit TIP3P water using AMBER tools, followed by a 100 ns SD simulation using AMBER 20: PMEMD CUDA and the ff14SB force-field. WATMD calculations were performed on the last 10 ns (40,000 frames) of each trajectory. All of the resulting structures and voxel grids were aligned to a common reference frame consisting of cereblon or CK1α in structure F (see Table 1) using the align function of PyMol.

### LMW SD simulation protocol

The LMW structures listed in Table 2 were extracted from crystal structures or modeled using the Build function of Maestro (as noted in the table) and pre-processed with GAFF in ANTECHAMBER (charges generated using the AM1-BCC method). The structures were solvated in explicit TIP3P water using AMBER tools, followed by 100 ns restrained SD simulation using AMBER 20 (ff99SB force-field). WATMD calculations were performed on the last 10 ns (40,000 frames) of each trajectory. All of the resulting structures and voxel grids were aligned to a common reference frame consisting of lenalidomide in structure F (see Table 1) using the align function of PyMol.

### Uridine docking protocol

We built the uridine structure de novo using Maestro, followed by minimization in MacroModel (default settings). We then docked the structure in the degrader pocket of cereblon taken from 5FQD using the Glide XP tool of Maestro (default settings), which resulted in a single pose that we used as input for LMW WATMD calculations.

## Results

### The general dependence of ΔGenriched and ΔGdepleted on solute surface composition, topology, and curvature

Here, we summarize the <u>general learnings</u> from our previous WATMD calculations on COVID M^pro4^ and LMW hERG blockers,^5^ together with cereblon and the degrader drugs studied in this work. Preferences among the sterically-accessible conformational states of a given solute are determined principally by solvation H-bond free energy (versus interatomic energy contributions), which varies spatially across solute surfaces, as follows:

1) **Type, number, and distribution** of polar/charged acceptor/donor groups versus non-polar groups across non-buried surface regions.
2) **H-bond propensity (charge, polarity)** of solute donor/acceptor groups relative to that of water-water H-bond groups.
3) **Regional concavity** (<u>depth below</u> the mean surface elevation) versus convexity (<u>height above</u> the mean surface elevation) across the surface. Optimization of the solvating water-water-solute H-bond network geometry depends on ordering-dependent adaptation to the solute surface geometry, resulting in varying degrees of entropic loss.
4) **Regional surface curvature magnitude**, which determines the requisite degree of geometric H-bond network adaptation and ordering.
5) **Solvent accessibility** of each donor/acceptor group, as reflected in the <u>local</u> solvent exposed surface area, which ranges between uni- and omni-directional at high versus low exposed surface regions, respectively.

The aforementioned characteristics of solvent exposed surfaces differ fundamentally between LMW and HMW solutes due to the significantly greater folding/side chain packing ability of the latter relative to the former, together with the lack of buried surfaces in LMW solutes, as follows:

1) LMW solute surfaces are contributed largely by <u>individual</u> loosely packed/unpacked

substituents/R-groups, whose surfaces range from planar (e.g., the core of aromatic rings) to convex (e.g., acyclic groups and the edges of aromatic rings) to concave (e.g., inter- substituent connections). We refer to LMW surface geometries as <u>first order</u> convexity/concavity (Figure 17A). Single strongly polar groups projecting from convex surfaces into solvent promote G_enriched_ < (or <<) G_bulk solvent_ (Figure 17B), whereas non-polar convex surfaces may promote G_enriched_ > (or >>) G_bulk solvent_ (depending on the magnitude of the surface curvature and degree of entropic loss due to additional ordering at such regions) (Figure 17C).

1) HMW solute surfaces consist largely of the outer boundaries of their packed side chains and backbones, which we refer to as <u>second order</u> convexity (Figure 18A) and concavity (Figure 18B). Co-localization of remote H-bond groups in the monomer sequence may occur in <u>concave</u> surface regions due to folding. H-bond enriched solvation is often promoted by such environments (Figure 18C). High <u>convex</u> surface curvature in proteins maps largely to loops and β-turns. Buried water channels/cavities may additionally arise in folded structures.
2) Foldability/packability of HMW solutes enables far greater solvation-powered rearrangeability of surface convexity/concavity, curvature, and polar/non-polar surface composition compared with LMW solutes.

**Figure 17.**
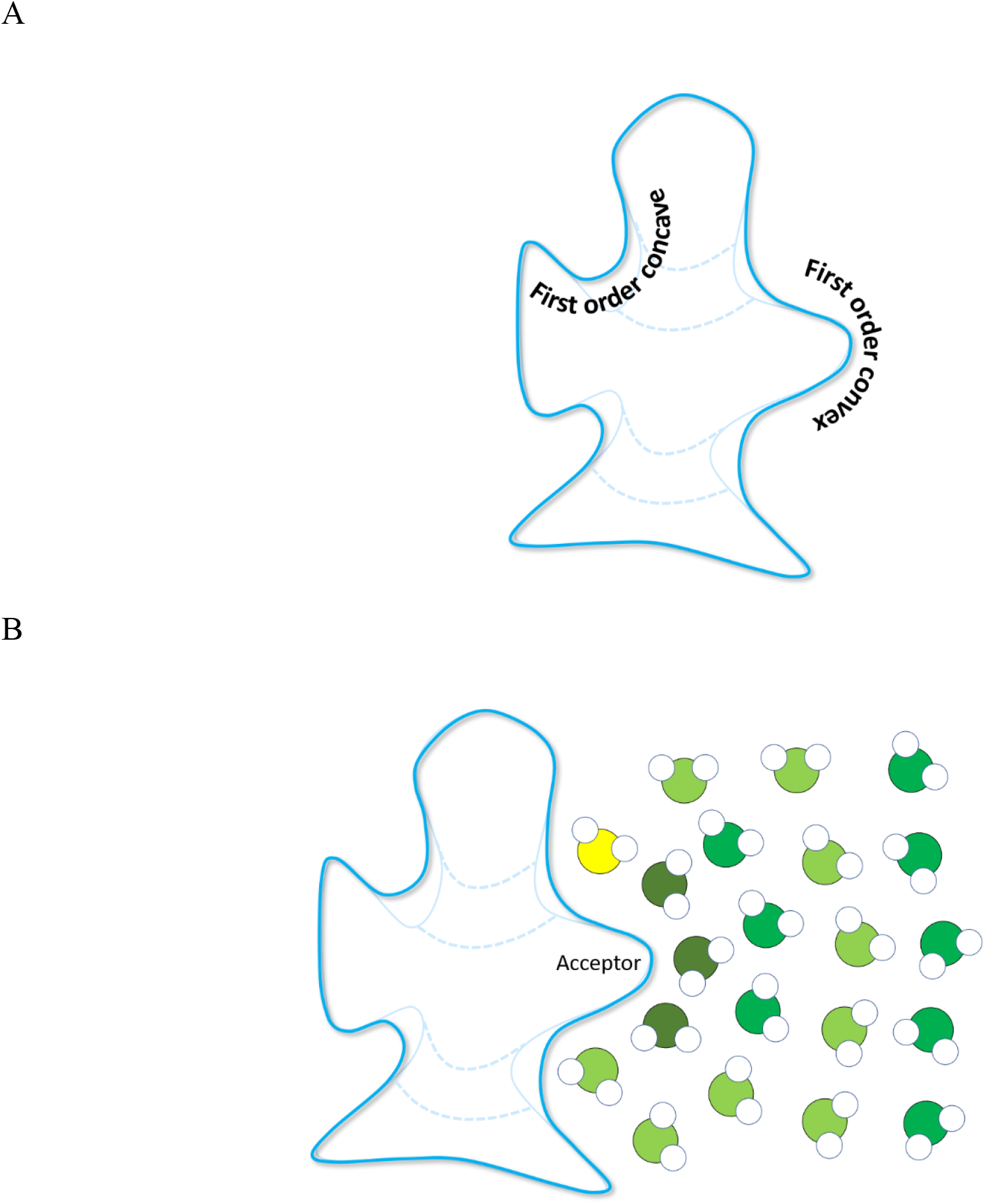
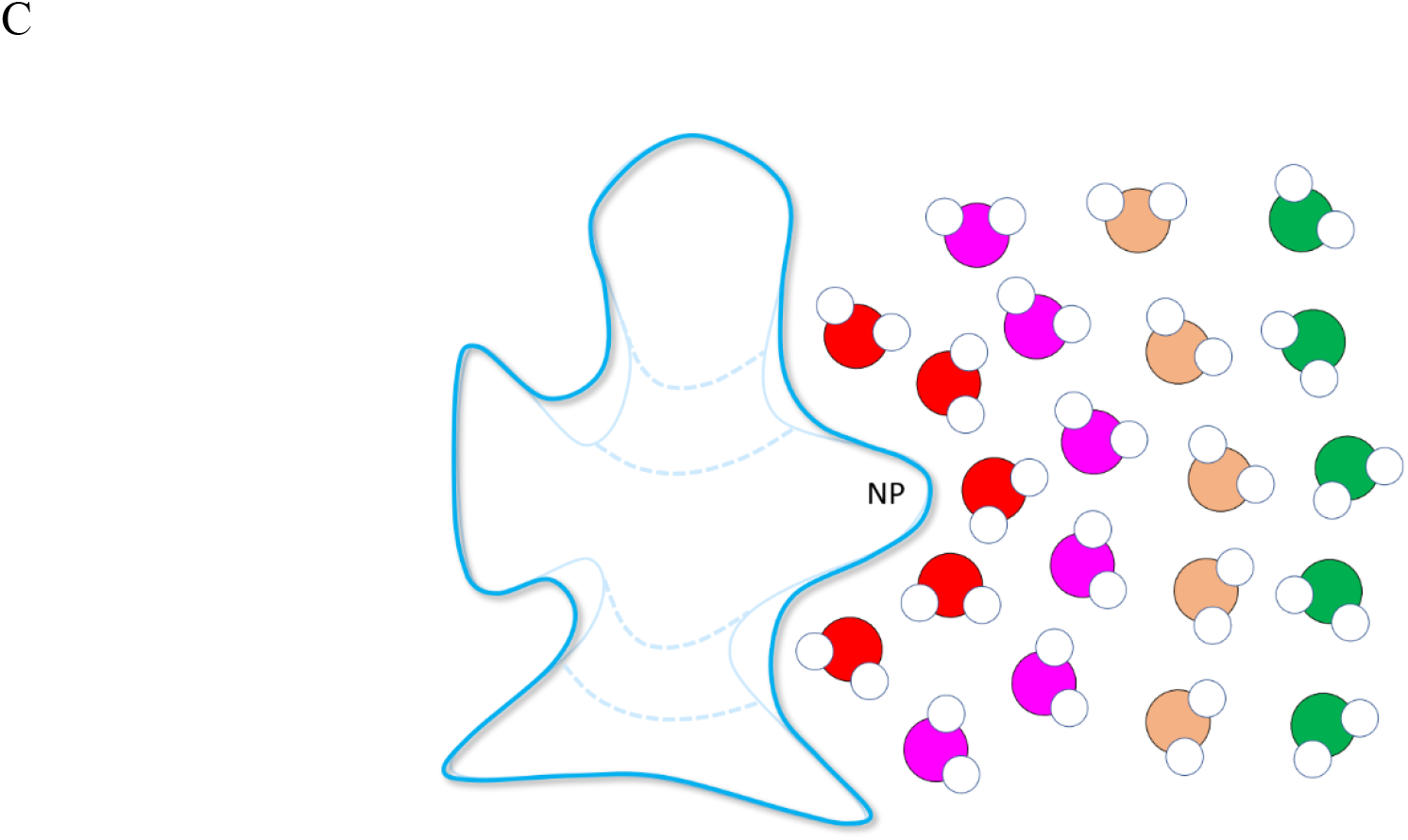
Hypothetical cartoons exemplifying the general properties of the non-isotropic H-bond fields of <u>LMW solutes</u>, which are strongly influenced by solute surface properties, including convexity/concavity, local curvature, and polar/non-polar composition. Solvation fields seek to minimize their free energy in the presence of dissolved solutes (which may reside above or below that of bulk solvent) via intra- and/or inter-solute rearrangements corresponding to optimal solute surface curvature, convexity/concavity, and polar/non-polar group exposure. Depending on the local polar/non-polar surface composition, high surface curvature may result in severe distortion of the water H-bond network geometry, accompanied by the loss or gain of free energy (noting that entropic losses are expected at highly distorted H-bond network regions due to additional ordering in such regions). (A) The surfaces of LMW solutes are contributed largely by <u>individual</u> (i.e., non-packed) chemical groups, which exhibit varying degrees of concave and convex curvature (which we refer to as first order curvature), ranging from planar to severely curved (e.g., at the core versus edges of aromatic substituents, respectively). Convex and concave surfaces map largely to substituents and inter-substituent connections, respectively. (B) Polar convex surfaces incur a tradeoff between favorable H-bond enrichment and unfavorable warping of the water H-bond field. (C) Non-polar convex surfaces (denoted as NP) of LMW solutes induce varying degrees of strain on the H-bond field, which are both H-bond depleted and warped.

**Figure 18.**
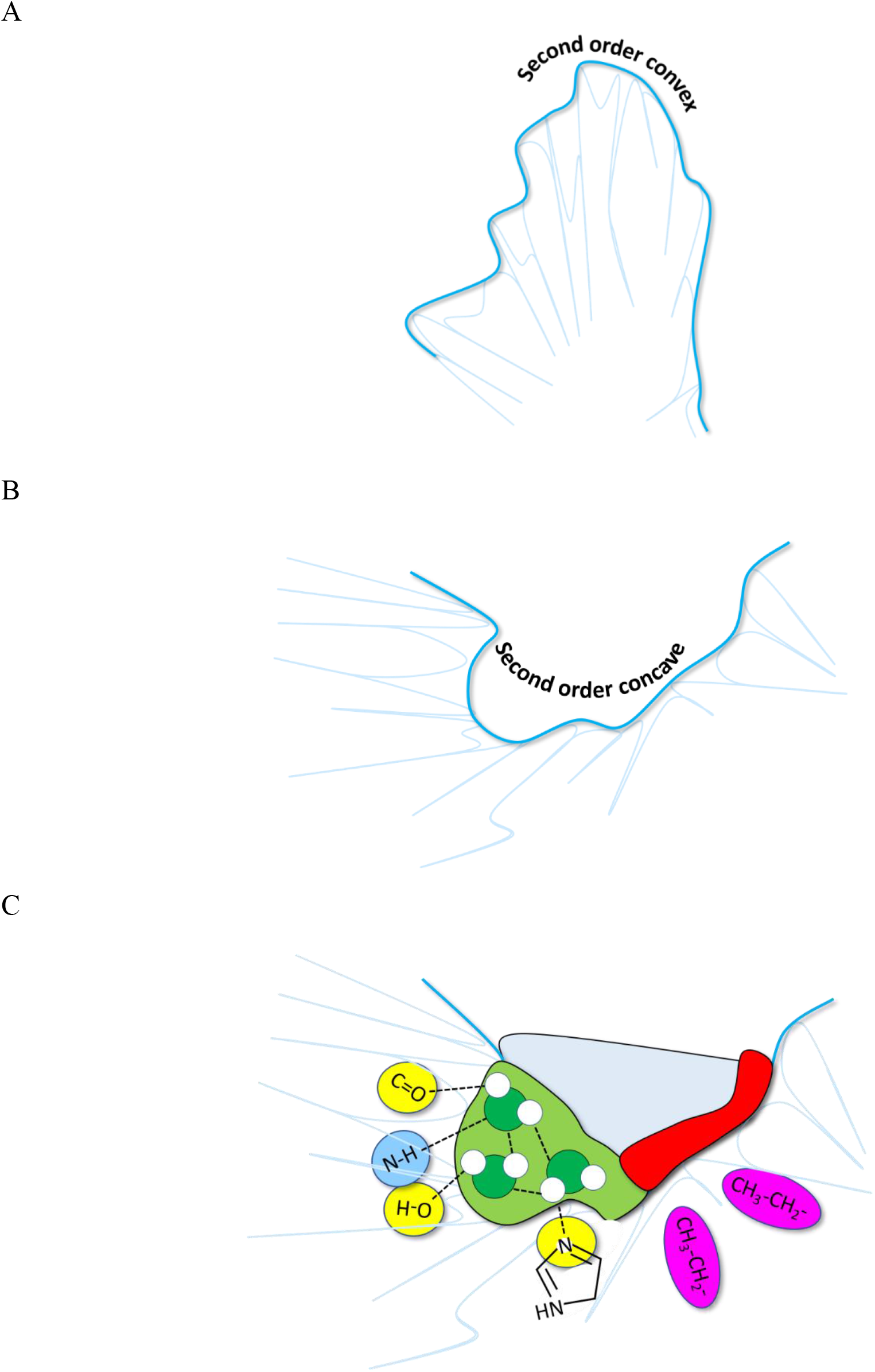
As for LMW solutes, the non-isotropic solvation fields of HMW solutes are strongly influenced by local surface curvature and polar/non-polar composition. The surfaces of <u>HMW solutes</u> are contributed largely by <u>packed</u> side chain and backbone moieties, where curvature is a collective property of the packed moieties (which we refer to as second order curvature). (A) Highly curved <u>convex</u> surfaces are typically associated with loops and β-turns. (B) Highly curved <u>concave</u> surfaces typically map to the interfaces between secondary structure elements and domain interfaces. (C) Folded concave surfaces may result in co-localization of remote H-bond groups in the monomer sequence, promoting H-bond enriched solvation of such regions (light green background) interspersed with H-bond depleted solvation (red background).

### Case study: A top-down analysis of E3 ligase-mediated protein degradation

We set about to qualitatively analyze:

1) Proteasomal protein degradation via a symbolic mathematical model of a representative subset of the pathway in terms of:

a) The extrinsic and intrinsic rates of assembly of the substrate-degrader-bound E3 complex, buildup of the poly-ubiquitylated substrate species and decay thereof via deubiquitylation (the Yang of degradation) and proteasomal degradation (the Yin of degradation) (noting that Clocks were omitted for the sake of simplicity).
b) The optimal binding kinetics of Type I and II degraders needed to achieve the maximum rate of ternary complex formation (yielding the degradation rate >> rate of substrate binding to its downstream effector) at the lowest possible degrader concentration.
2) The following solvation properties that we used to deduce the functionally important non-equilibrium state transitions of cereblon, including binary and ternary complex formation:

a) Complementarity between the solvation fields of cereblon and Type I degraders governing binary complex formation analyzed using WATMD (a solvation field perspective on structure-activity relationships).
b) The role of cereblon-Type I degrader complexes (e.g., cereblon-lenalidomide) in promoting substrate binding.
c) Complementarity between the binary degrader-cereblon solvation field and that of CK1α, which governs both ternary complex formation and the lifetime thereof (chosen based on the availability of a published crystal structure of the ternary complex).
d) The solvation field of cereblon-degrader-CKα vis-à-vis stability of the ternary complex.
e) The role of buried water channels in powering the intra- and intermolecular state transitions of cereblon.
f) Rotation of the CULT domain relative to the N-terminal domain (denoted as the “open” state) (6H0F).

### Analysis of proteasome-mediated protein degradation pathway dynamics

The cereblon-mediated protein degradation pathway examined in this work is outlined schematically in Figure 19 and formulated as a symbolic mathematical model provided in the Supplementary Information. The model includes native and non-native (“neo”) substrate recruitment by endogenous LMW degraders and drugs.

**Figure 19.**
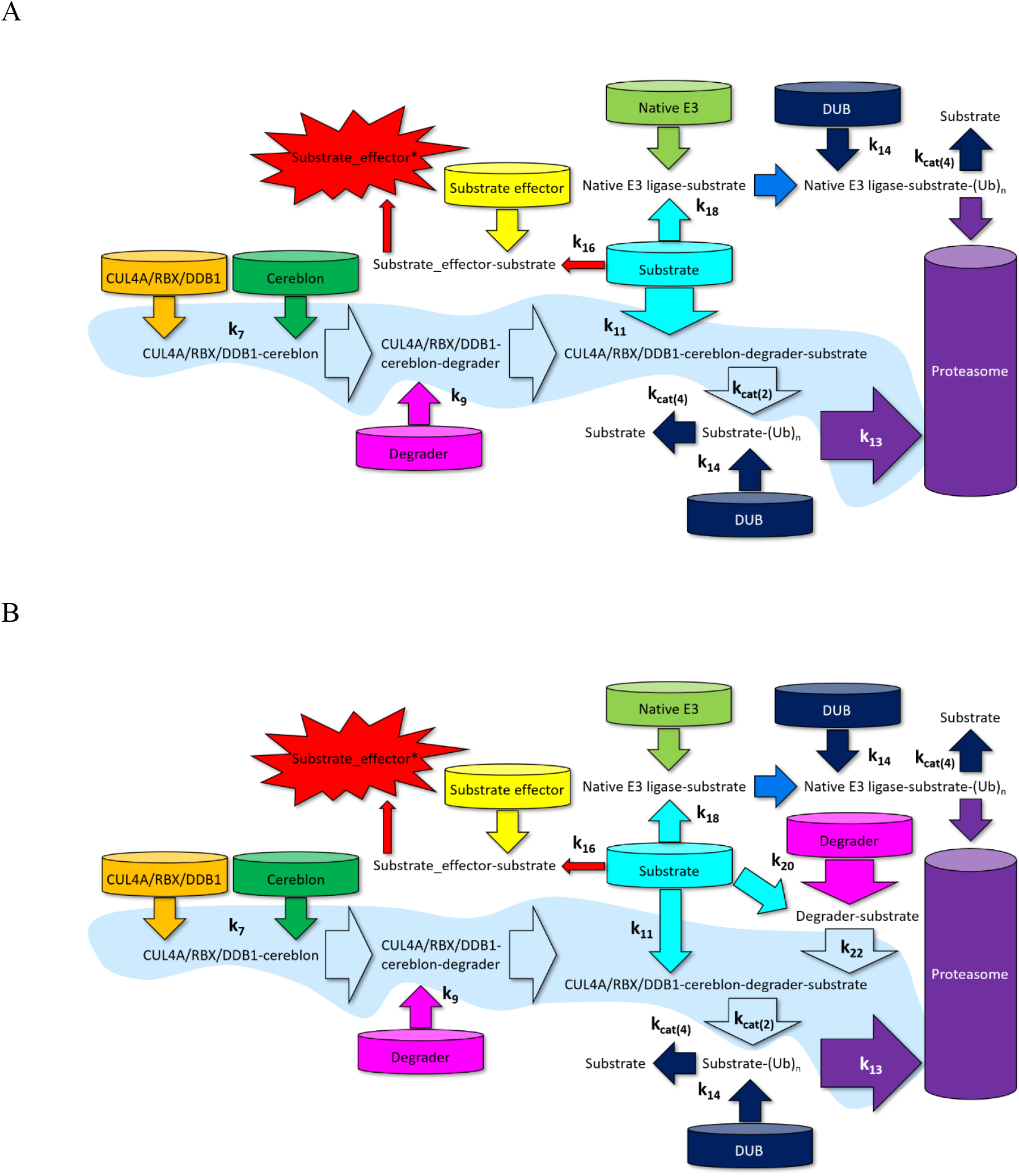
Cartoon of the cereblon-mediated degradation pathway schema detailed in the text (including DUB counter-balancing), with the main trunk highlighted in light blue. Some reactions and rate constants have been condensed and Clocks omitted for the sake of simplicity. (A) Type I degraders (e.g., lenalidomide, pomalidomide) and their targeted substrates (e.g., CK1α) and downstream effector of the target substrate. The substrate degradation rate (k_cat(4)_) is optimally >> the rate of downstream substrate-mediated effector activation (k_16_) and the rate of binary cereblon- degrader formation (k_9_) is optimally ≈ the rate of cereblon buildup (k_7_). (B) Type II degraders (e.g., dBET23) and their targeted substrates (e.g., BRD4). The rate of degrader-substrate binding (k_20_) is ideally > the rate of degrader-cereblon binding (k_9_), which promotes substrate inhibition and degradation, while limiting cereblon inhibition.

**Achieving efficacious cereblon-degrader-substrate dynamic occupancy (corresponding to a threshold inhibition of the downstream neo substrate effectors) at the lowest possible degrader concentration depends on kinetic tuning between the degrader and the overall pathway**, and in particular:

#### For Type I degraders (Figure 19A)

1) Degradation rate (k_13_) >> substrate + substrate_effector k_on_ (k_16_).
2) Substrate + CUL4A/RBX1/DDB1-cereblon-degrader k_1_ (k_11_) ≈ [substrate](t).
3) Fast degrader + CUL4A/RBX1/DDB1-cereblon k_on_ (k_9_).
4) CUL4A/RBX1/DDB1-cereblon-[degrader] k_off_ < CUL4A/RBX1/DDB1-cereblon- degrader + substrate on-rate (k_1_ · [substrate](t)).
5) Fast degradation via the native ligase (i.e., fast k_18_).

#### For Type II degraders (Figure 19B)

1) Degradation rate (k_13_) >> substrate + substrate_effector k_on_ (k_16_).
2) Degrader + substrate k_on_ (k_20_) ≈ [substrate](t).
3) CUL4A/RBX1/DDB1-cereblon-[degrader-substrate] k_off_ < CUL4A/RBX1/DDB1- cereblon-degrader-substrate k_cat_.
4) Degrader + substrate k_on_ (k_20_) > degrader + CUL4A/RBX1/DDB1-cereblon k_on_ (k_9_), given that CUL4A/RBX1/DDB1-cereblon is inhibited in the latter case, and the substrate is both inhibited and degraded in the former.
5) CUL4A/RBX1/DDB1 + degrader-substrate k_on_ (k_22_) > CUL4A/RBX1/DDB1 + degrader k_on_ (k_9_).
6) k_9_ is rate-limiting when slower than k_20_ but must be balanced against k_16_.
7) Fast ternary complex formation in both directions (i.e., fast k_22_ and k_11_).
8) Degradation is faster than deubiquitylation (i.e., k_13_ >> k_14_).
9) Fast degradation via the native ligase (i.e., fast k_18_).

### Thalidomide-cereblon-CK1α complex formation is precluded by steric incompatibility

CK1α is recruited by lenalidomide-cereblon, but not thalidomide-cereblon, which can be explained largely on the basis of steric incompatibility between the phthalimide ring of thalidomide and substrate β-hairpin.^43^ The backbones and side chains of thalidomide- and lenalidomide-bound cereblon are well-aligned, ruling out the possibility of binding mode differences per se. Rather, the phthalimide ring of thalidomide is rotated about the glutarimide nitrogen (N) toward the β- hairpin of CK1α (Figure 20A), stemming largely from puckering differences in the glutarimide ring in the presence of a second unique carbonyl oxygen (O) (noting that the glutarimide ring binding geometry is highly similar in both cases) (Figure 20B). The phthalimide ring position in lenalidomide is well-packed with the β-hairpin of CK1α (Figure 20C), which clashes in thalidomide (Figure 20D).

**Figure 20.**
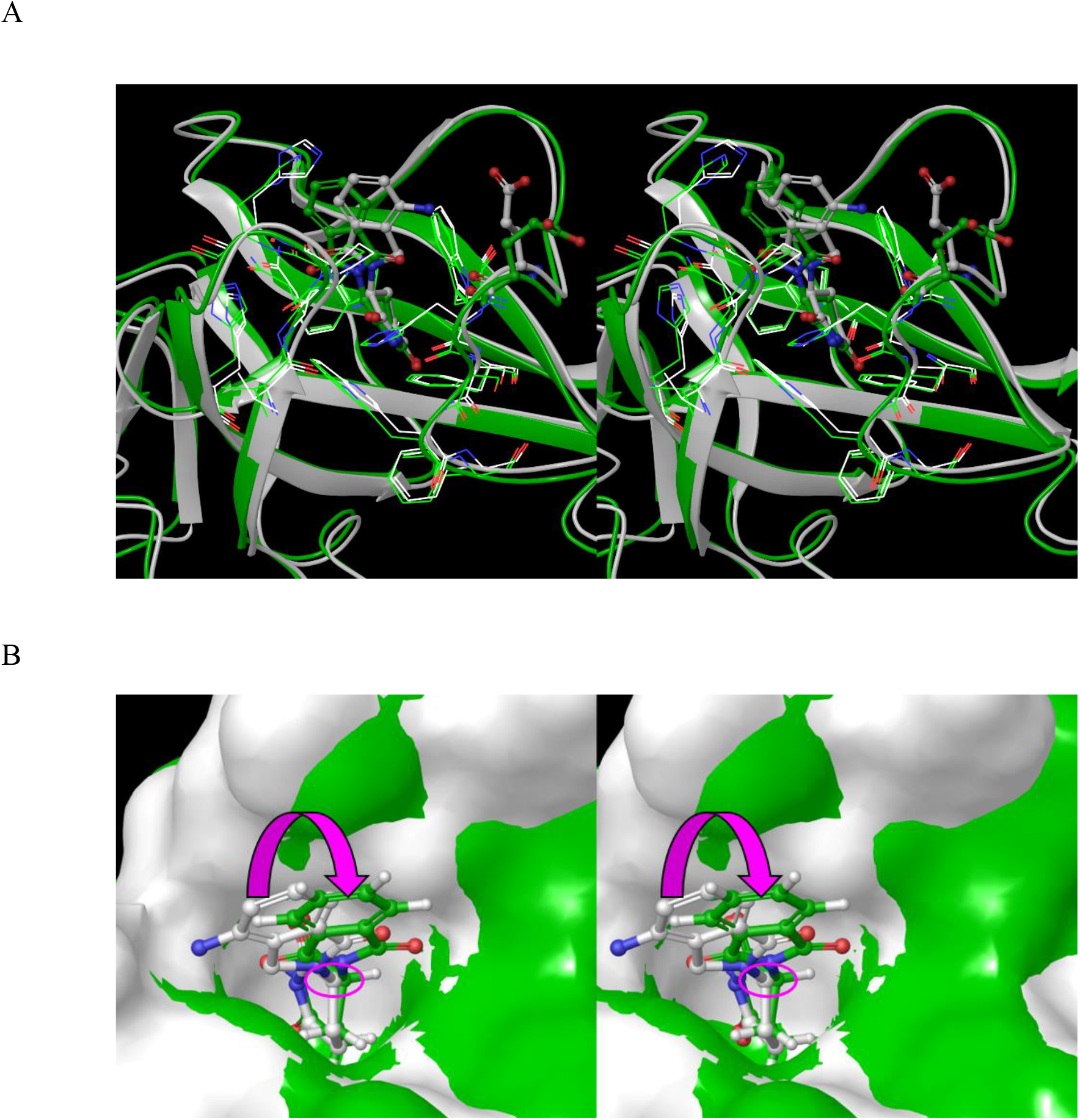
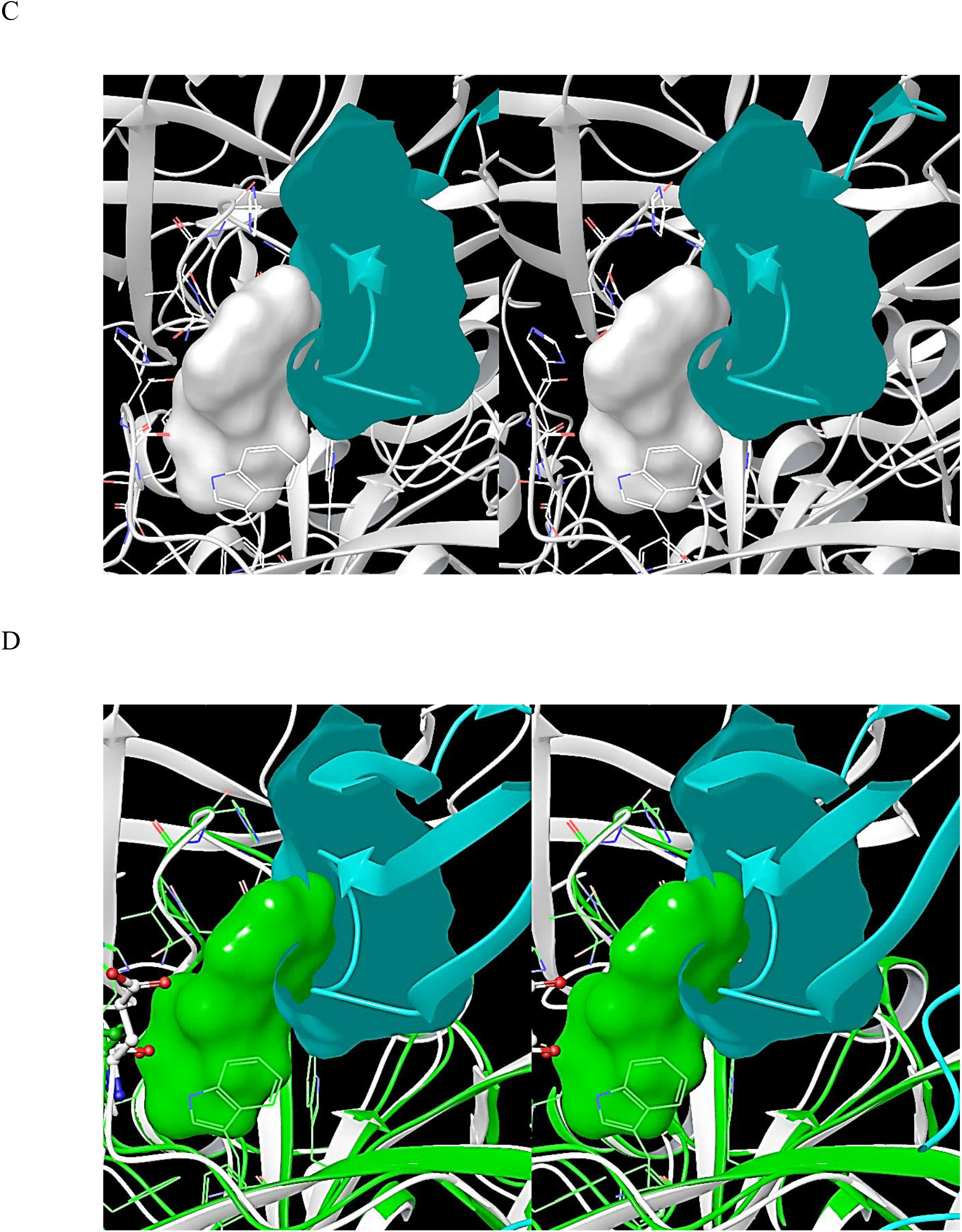
Stereo views of cereblon-thalidomide (S isomer) extracted from PDB code = 4CI1 (green) overlaid on cereblon-lenalidomide-CK1α (S isomer) extracted from 5FQD (white). (A) A large rotation in the phthalimide group position between the two structures is apparent, resulting largely from puckering differences in the glutarimide ring. (B) Overlay of thalidomide and lenalidomide based on the CULT domain of cereblon, demonstrating high conservation of the glutarimide ring binding mode, despite rotation of the phthalimide group (magenta arrow) about the ring N (magenta circle). Binding of thalidomide to cereblon is accommodated by the openness of the substrate-facing side of the ligand pocket in the absence of CK1α. (C) The phthalimide ring position in lenalidomide is well-packed with the β-hairpin of CK1α, which explains the selectivity of this compound for this substrate. (D) The phthalimide ring position in thalidomide is sterically incompatible with the β-hairpin of CK1α (teal blue) which precludes cereblon-thalidomide-CK1α ternary complex formation.

### The structural dynamics of cereblon and its binding interfaces

Next, we set about to study the structural dynamics of cereblon based on the 100 ns SD simulations described in Materials and methods, the prerequisite for WATMD solvation field predictions (noting that only high frequency protein rearrangements occur on this timescale). Our preference is to use experimental rather than predicted structures for this purpose, given that solvation-dependent free energy contributions are grossly underweighted in conventional MD simulations and other force-field-based approaches, as noted above. However, local rearrangements may be predicted from conventional MD simulations on a case-by-case basis in the absence of experimental structures. Our study focused on the mechanism by which cereblon is switched to its substrate binding-competent state by bound degraders (focused qualitatively on the intrinsic rates), the possible scenarios of which consist of either increaed substrate k_1_ via stabilization of the binding-competent form of the substrate binding surface, or decreased k_-1_ via degrader-contributed augmentation of the H-bond depleted solvation of the substrate binding surface. The small fractional increase in the substrate-binding surface area and H-bond depleted solvation free energy contributed by bound degraders is inconsistent with the total absence of substrate binding in the degrader-unbound state (Figure 21) (the H-bond depleted solvation of unbound and bound lenalidomide and thalidomide is addressed in a later section).

**Figure 21.**
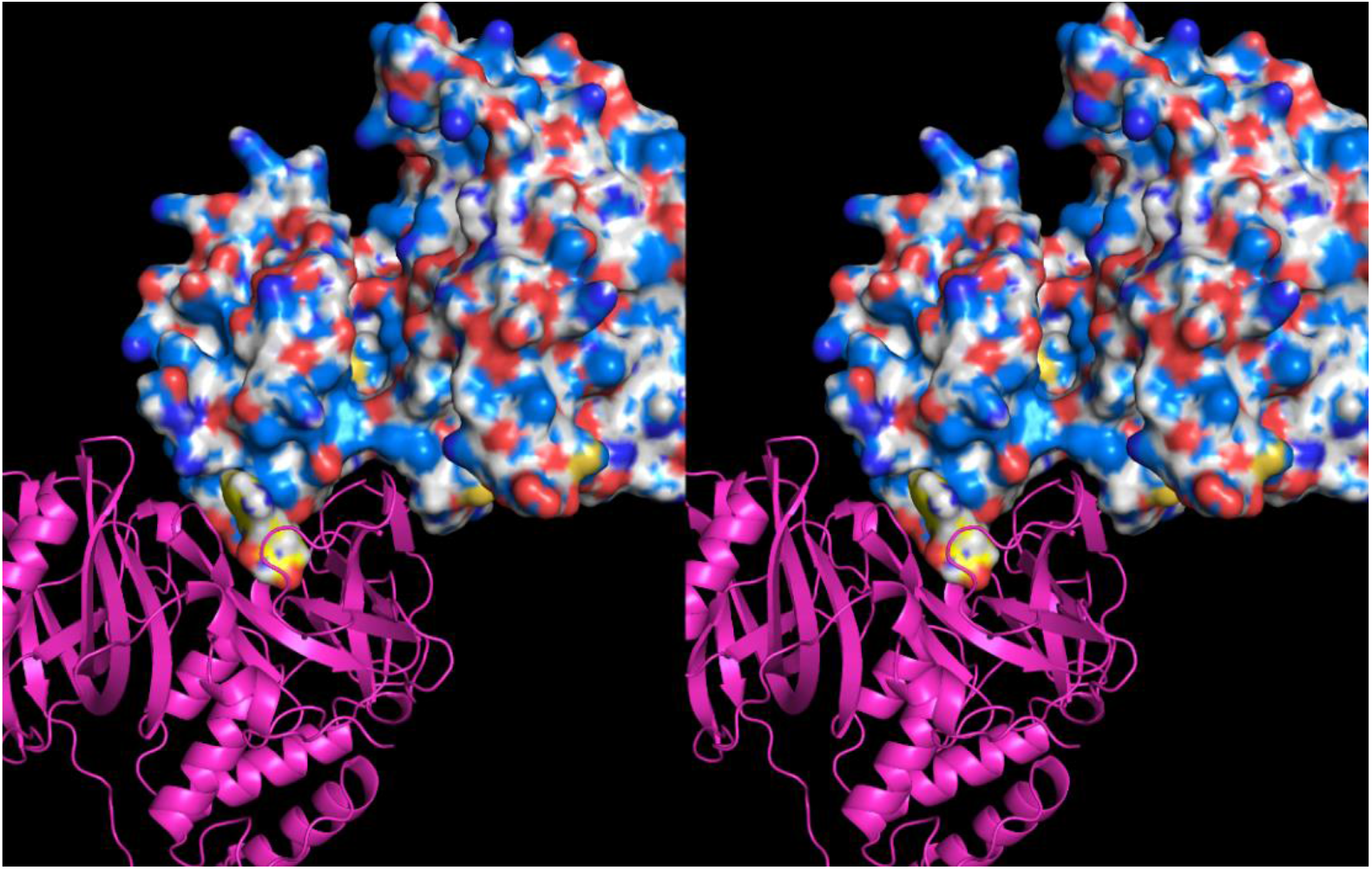
Stereo view of the CK1α-cereblon interface (blue surface and magenta cartoon, respectively) in the ternary structure F. Overlay of structures F and A, looking from CK1α (blue) into pocket L. The small contributions of lenalidomide (yellow) to the H-bond depleted solvation and overall interfacial footprint are inconsistent with the complete absence of substrate binding in the degrader-unbound state of cereblon.

We therefore postulate that association between cereblon and its cognate substrates depends on degrader-mediated stabilization of the substrate binding site, resulting in dramatic speeding of substrate k_1_, as follows:

1) The empty form of the degrader pocket (widely referred to as the Trp-box, and denoted in this work as pocket L) cycles between degrader binding-competent and incompetent states

in our simulations, which correspond to fully formed and collapsed states of the pocket, respectively (noting that the fully formed empty pocket in human cereblon has not been observed experimentally as of this writing).

1) The substrate binding-competent state of cereblon corresponds to the degrader-bound state of the protein, in which key polar groups contributed by Asn305, His311, Arg327, and Trp354 are positioned for H-bonding to the β-hairpin and other residues of CK1α (Figure 22A) (noting that these H-bonds are absent in the starting ternary structure D (Figure 22B), but present in the time-averaged structure (Figure 22C)). We simulated, but did not perform WATMD calculations on the cereblon-palmalidomide-IKZF1 structure (6H0F), which likewise forms interfacial H-bonds during the simulation, including between the β-hairpin and Asn305.

**Figure 22.**
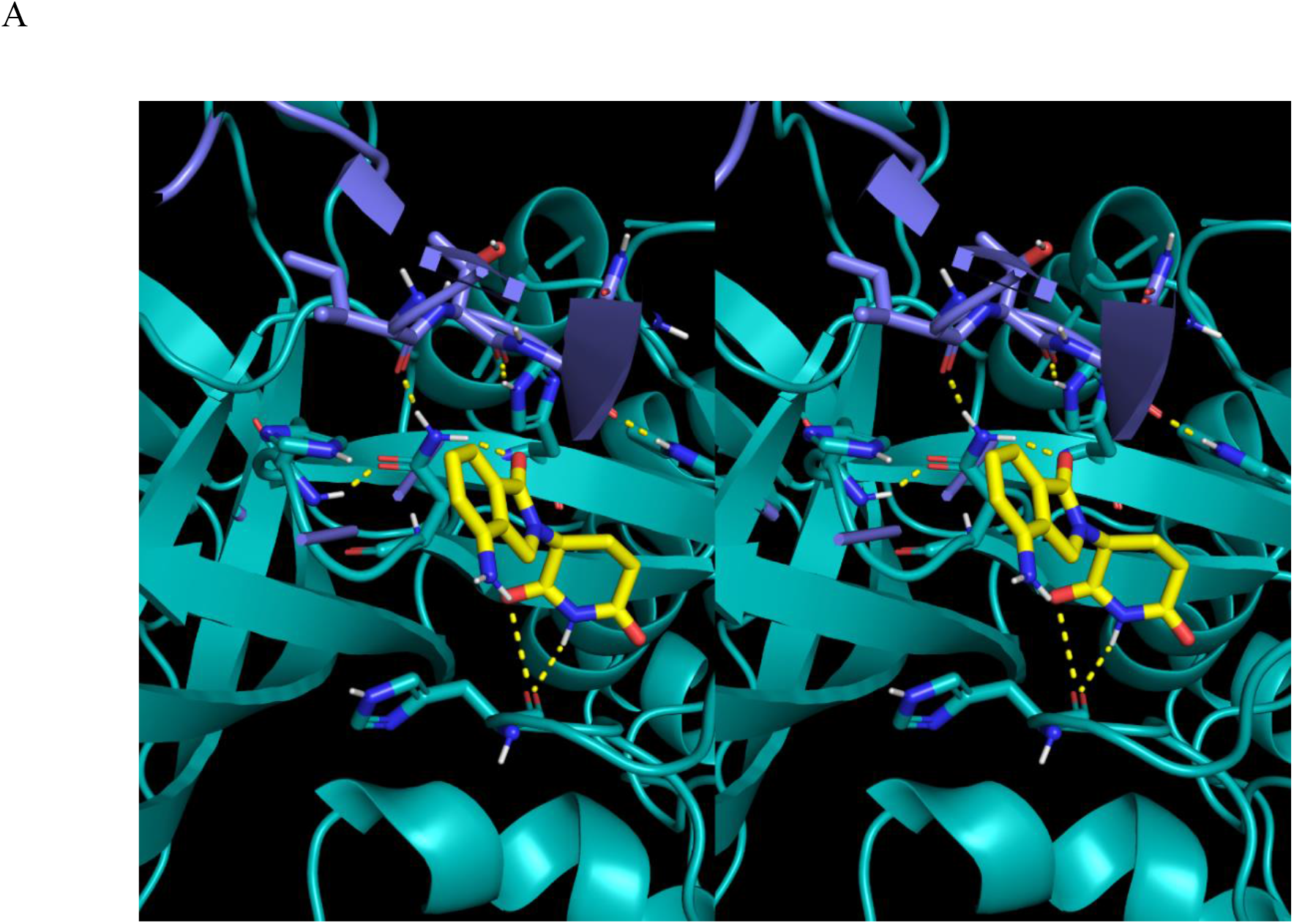
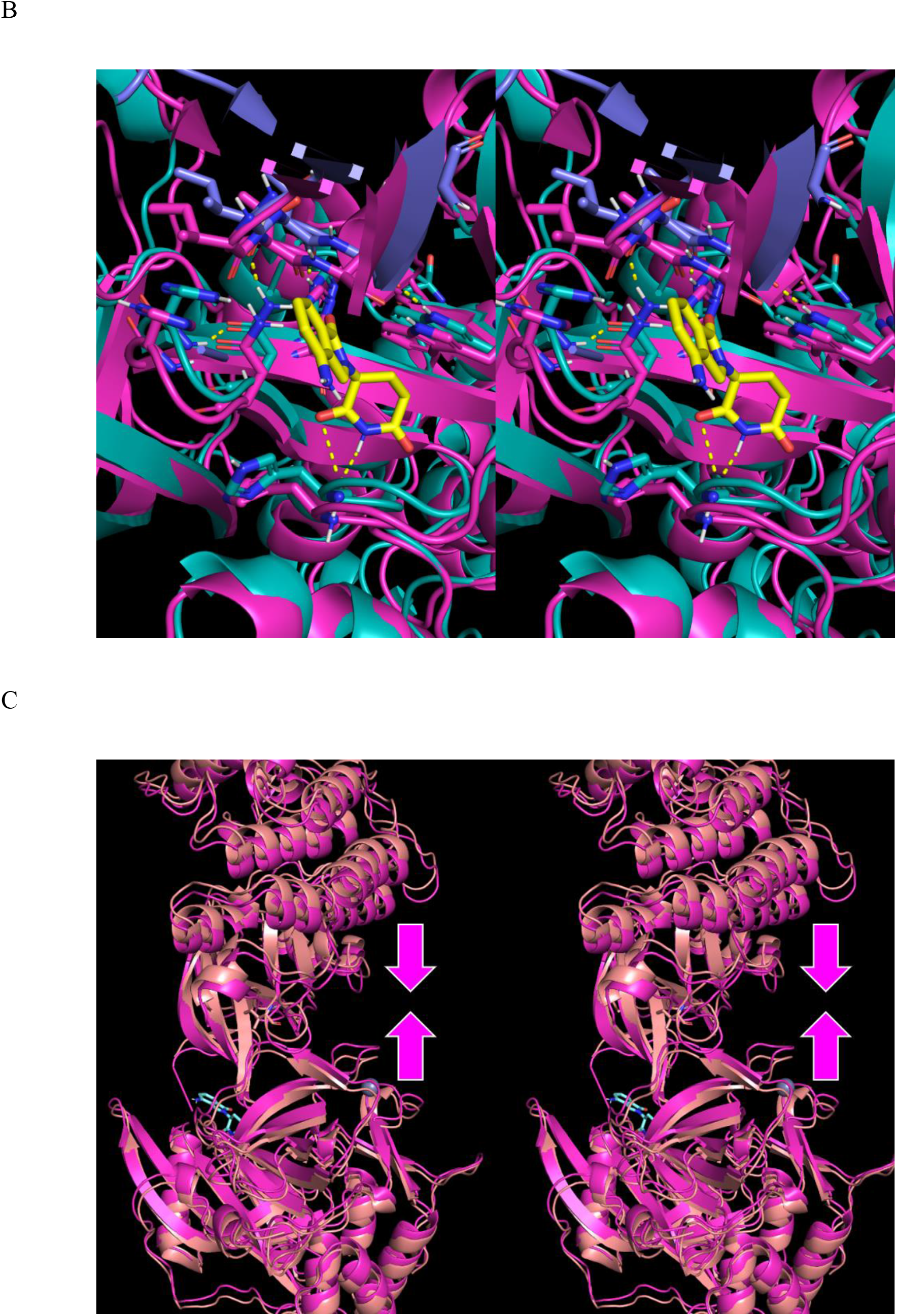
(A) The interfacial H-bonds (dotted yellow lines) between CK1α (purple), cereblon (cyan), and lenalidomide (yellow). Simultaneous alignment of these H-bond groups among the two partners permits close approach of the partners during ternary complex formation (noting that desolvation of H-bond depleted water from the interface, which is located along the protein surfaces, is dependent on tight surface-surface complementarity). Water H-bond replacement partners consist of cereblon His311 (ε-NH) to CK1α Thr419 (backbone O), cereblon Trp354 (indole NH) to CK1α Asn420 (backbone O), cereblon Asn305 (NH_2_) to CK1α Ile418 (backbone O), cereblon Arg327 (guanidium NH) to CK1α Glu423 (COO), lenalidomide phthalimide (O) to Asn305 (NH_2_), lenalidomide glutarimide (NH) to cereblon His332 (backbone O). <u>We hypothesize that complementarity among these H-bond partners depends on the presence of a bound degrader</u>. (B) Overlay of structures C (cyan) and D (magenta), showing the complete absence of interfacial H-bonds in D that are present in the time-averaged structure C. (C) Stereo view of structure F (magenta) overlaid on the initial ternary structure D (copper). The packing between cereblon and CK1α is further optimized in structures F and B (not shown) via an approximately rigid body translation (magenta arrows).

Next, we analyzed the dynamic behavior of the residues lining pocket L by comparing the time-averaged structures A_closed, A_closed’, A_open, C, C’, and H (Table 2). Pocket L is connected to a second transient pocket (denoted as pocket T) in the time-averaged structures A_closed, C, and F, but not C’ or D (in which it is occupied by the side chain of His378). Pockets L and T are lined by both unique and common residues (Figure 23A), and the surface openings reside at nearby regions of the protein (Figure 23B). The pockets and surface openings thereof are exemplifed in Figure 23C for structure A_closed.

**Figure 23.**
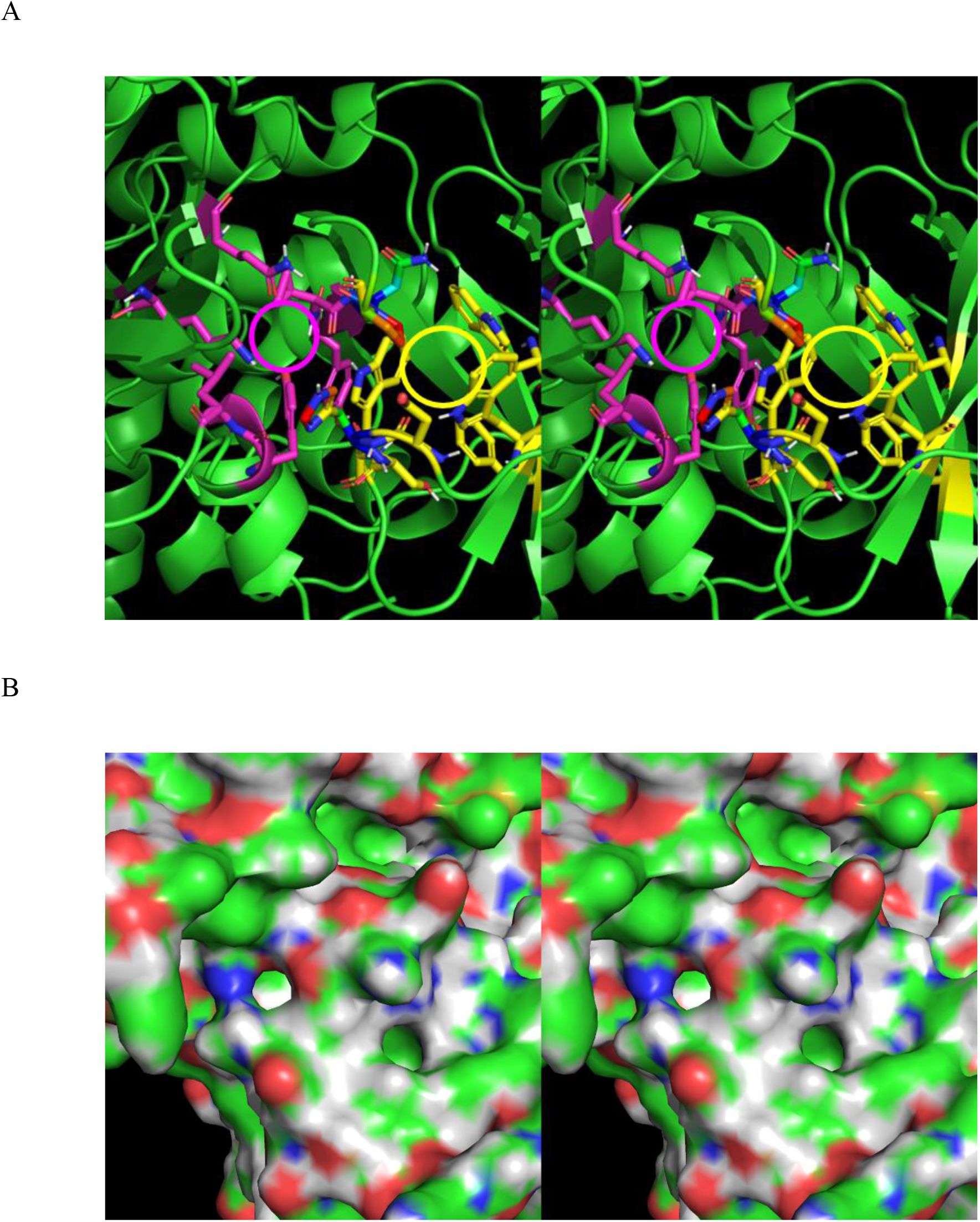
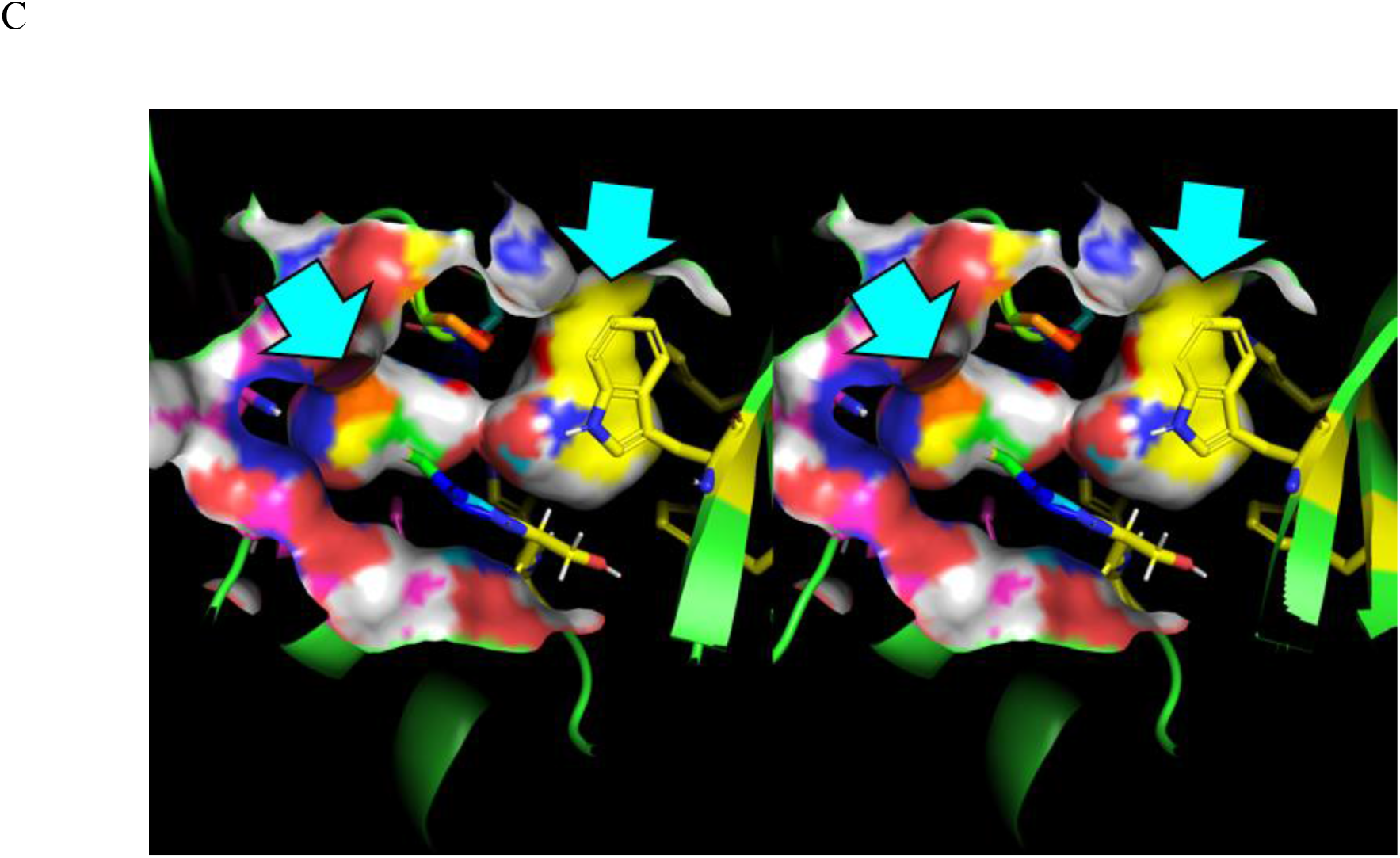
(A) Zoomed-in stereo view of the residues lining pockets L (Glu331, Ser333, Trp334, Trp340, Trp354, and Phe356) (yellow), T (Leu14, Tyr13, Gln54, Lys110, Tyr303, Val304, Pro306, and His332) (magenta), and both L and T (Asn305, Pro306, and His332) (color spectrum) in structure A. (B) Stereo view of the external surface openings of pockets T (northwest) and L (southeast) in structure A. (C) The buried surface of pockets L and T structure A.

We inferred the dynamic behavior of pocket L based on the time-averaged structures as follows (recalling that the starting structures used in our simulations are biased toward the ternary complex in 5FQD):

1) Fully formed in the degrader-occupied structures C, C’, and C” and the empty partially restrained structures H and A_closed” (exemplifed for structure C with lenalidomide shown for reference in Figure 24A).
2) Partially formed in structure A_closed (sampled exclusively in the hypothetical DDB1- unbound state) (Figures 24B), which is caused by counterclockwise shifts in the side chains of Trp334, Asn305, and Pro306 driven by twisting of the semi-circular β-strand 308-316 (Figure 24C).
3) Fully collapsed in structures A_open and A_closed’ (exemplified for structure A_open in Figures 24D and E). The collapsed pocket stems largely from rotation of the Trp340 side chain into the pocket, thereby retaining an H-bond between the indole NH and backbone O of Glu331, whose position shifts away from that in the other structures (e.g., structure F) (Figure 24F). We attribute this rearrangement to overweighting of the H-bond energy contribution in the AMBER force-field, which we postulate is right for the wrong reasons (addressed below).

**Figure 24.**
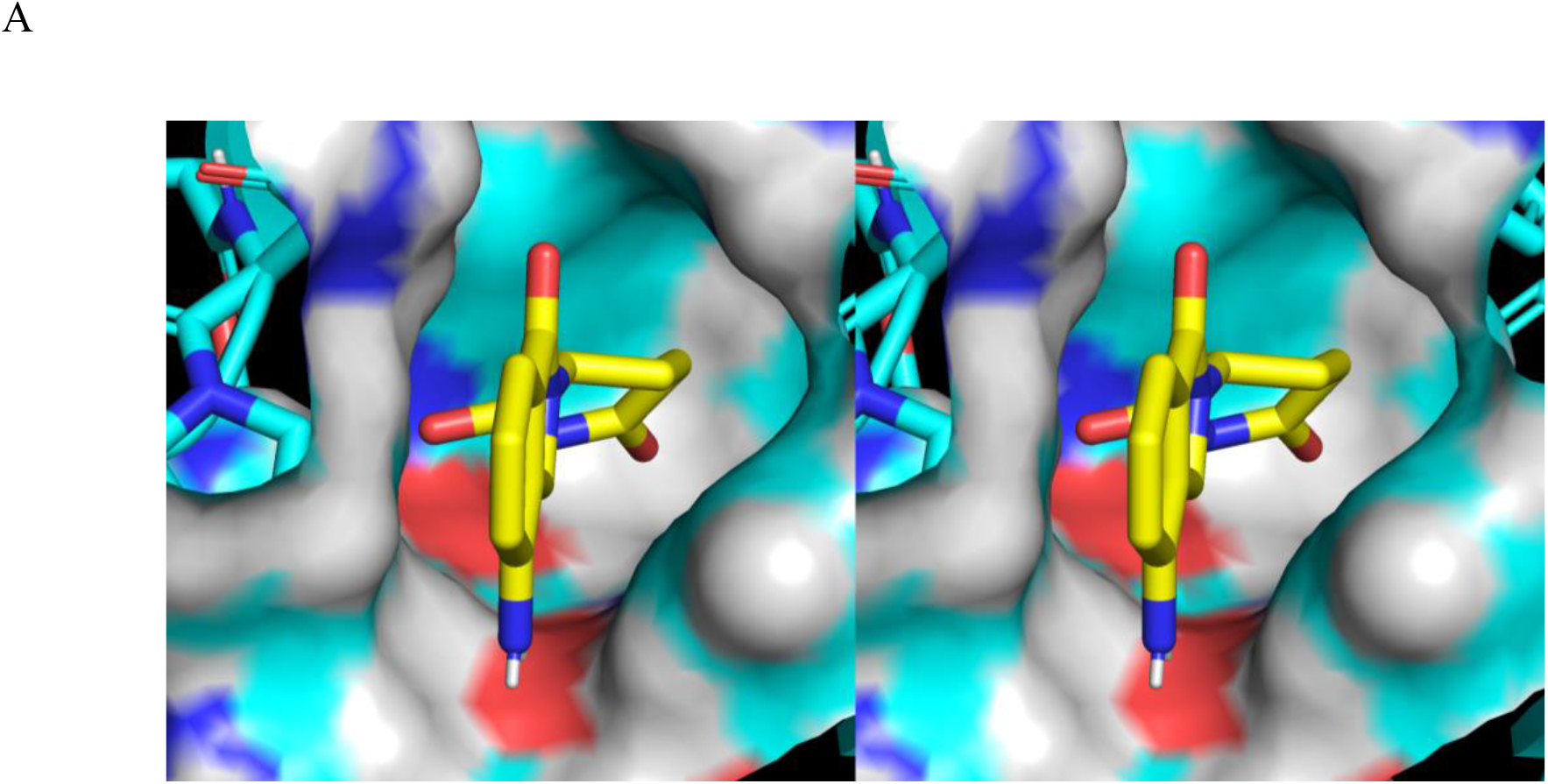
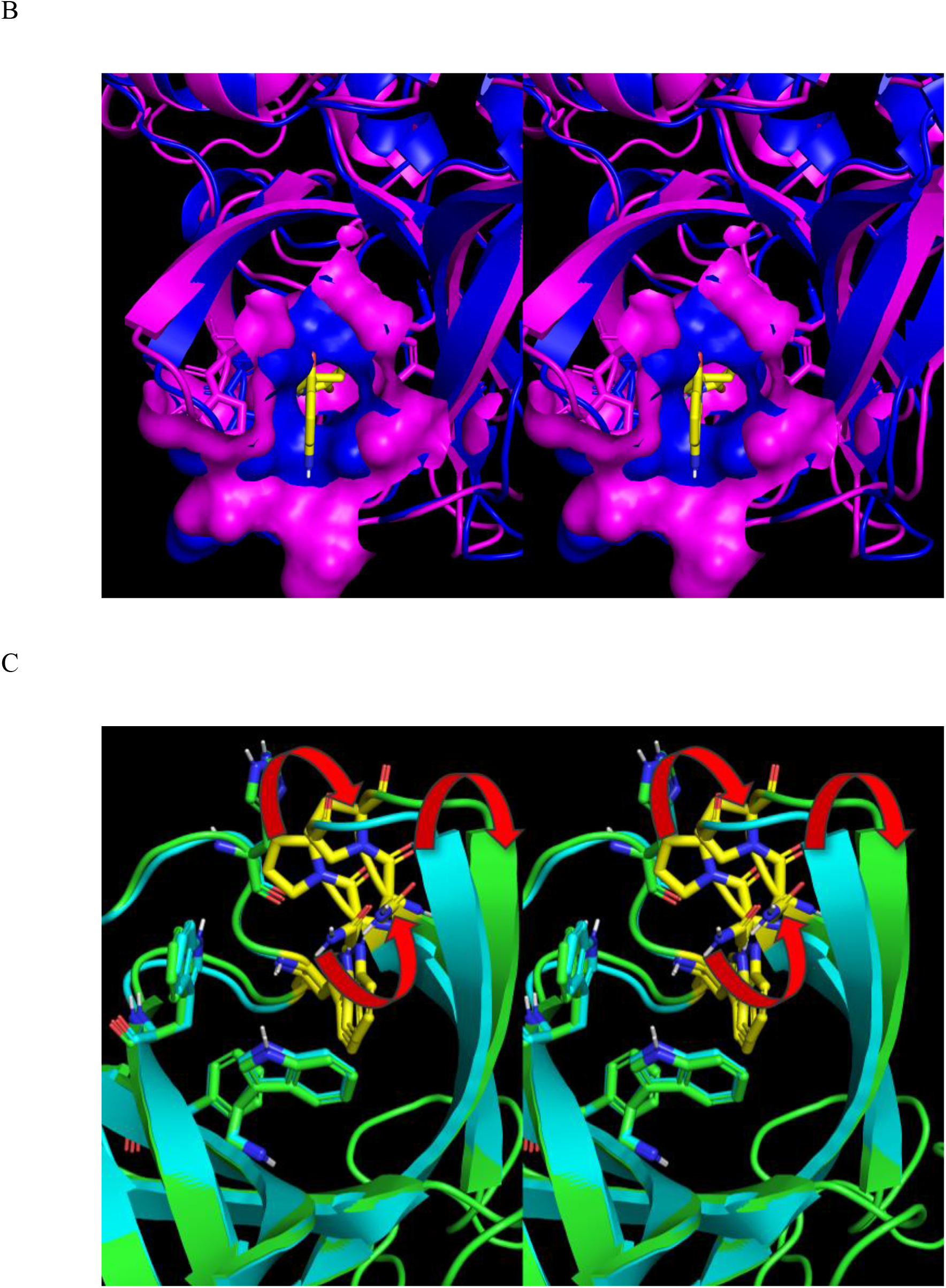
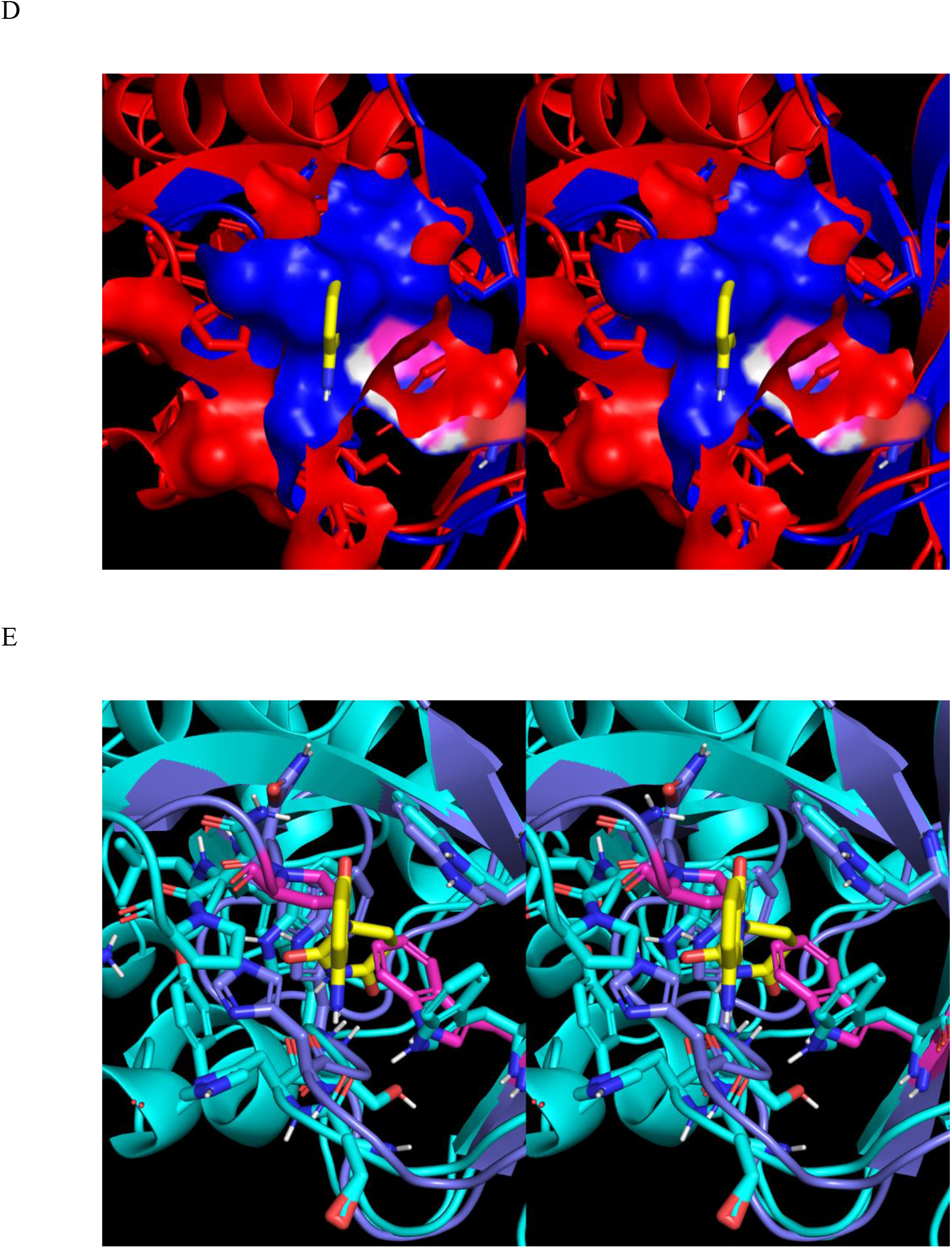
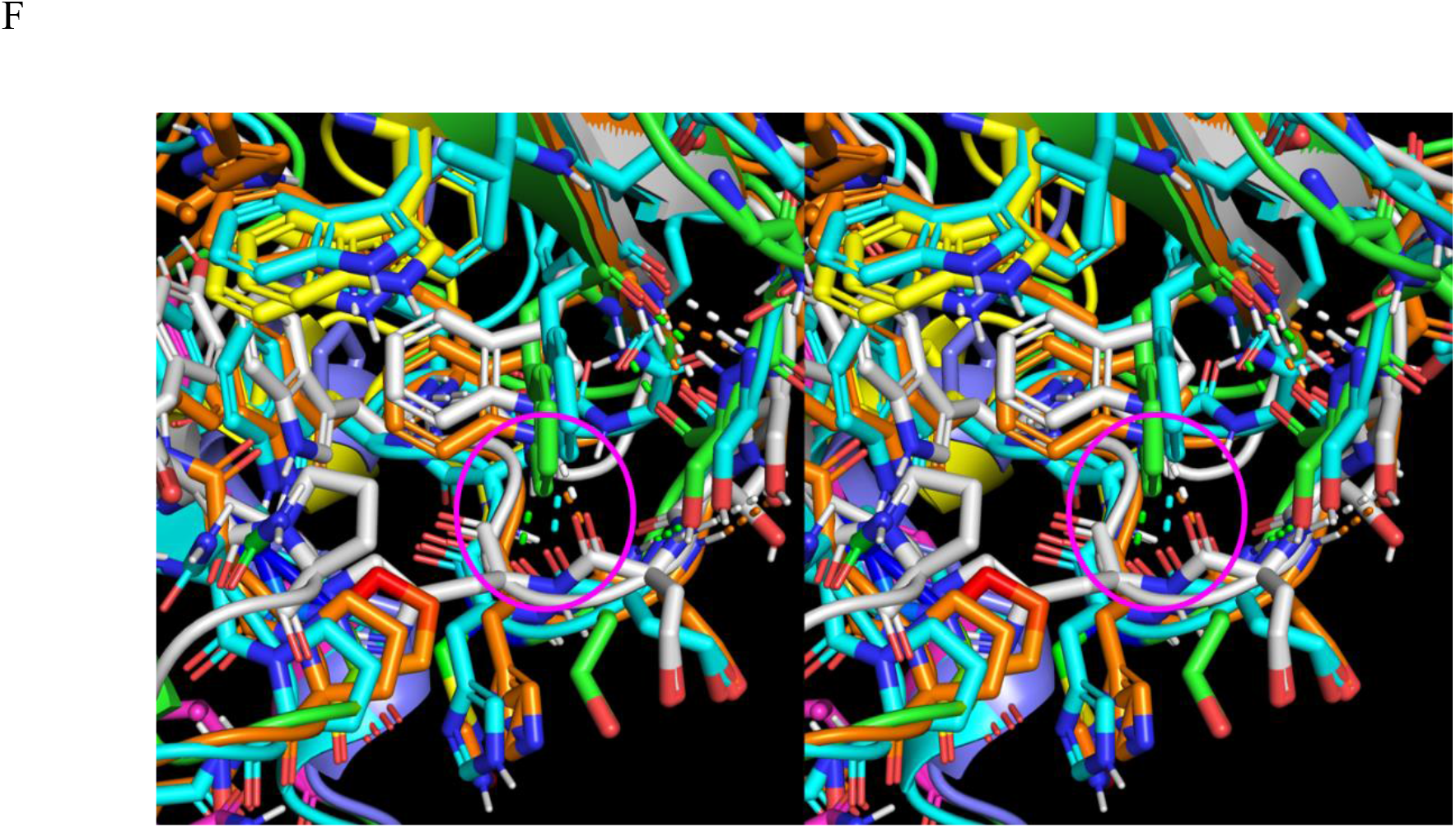
(A) Zoomed-in stereo view of lenalidomide (yellow) bound to pocket L in the binary structure C. (B) Zoomed-in stereo view of the opening of pocket L in structures A (blue) versus C (magenta), showing contraction of the opening in the empty versus lenalidomide-bound state. (C) Zoomed-in stereo view, showing the contracted diameter of the pocket L opening in structure A (green) relative to that of structure C (cyan) due to further twisting of β-strand 308-316 accompanied by shifts in the side chains of Trp334, Asn305, and Pro306 toward the pocket opening. (D) Zoomed-in stereo view of pocket L, which is fully collapsed in structure A_open (blue) relative to structure C (red) due largely to occupation by Pro306 and Trp340. (E) Zoomed-in stereo view, showing the rearranged residues lining pocket L in structure A_open (purple) relative to structure C (cyan). Pro306 and Trp340 are shown in magenta. (F) Zoomed-in stereo view, showing retention of the Trp340-Glu331 H-bond (circled in magenta) via Trp340 rotation in structures A_closed’ (orange) and A_open (white) versus A_closed (green) and C (cyan).

We assume that:

1) A_closed” (derived from 5FQD with partial restraints, as described in Table 1) and mouse cereblon (3WX2 and the time-averaged structure H with partial restraints) are reasonable models for the <u>fully formed pocket L as exists just prior to degrader binding</u>, which is unobserved experimentally (Figure 25).
2) A_closed (from which DDB1 was omitted) is a reasonable model for the <u>partially formed</u> pocket. The pocket and internal solvation in this predicted state are unstable relative to that of the collapsed and fully formed states, which we attribute to the omission of DDB1 in the time-averaged structure (addressed below).
3) A_closed’ (in which DDB1 was included) and A_open (in which DDB1 was likewise included) are reasonable models for the <u>fully collapsed</u> pocket.

**Figure 25.**
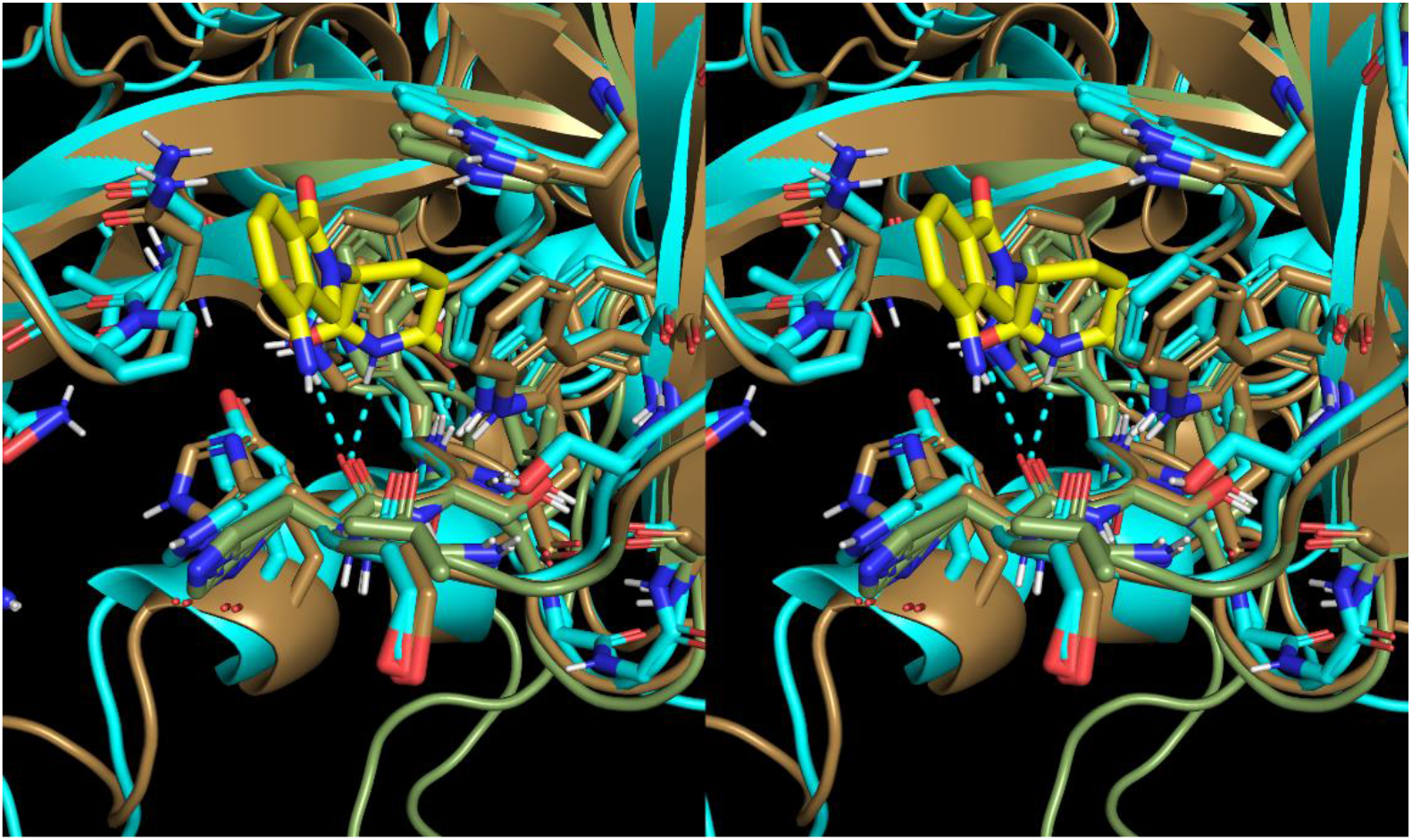
Overlay of the lenalidomide-bound structure C (yellow sticks and cyan cartoon, respectively) on structures A_closed” (tan cartoon) and H (olive green cartoon), which serve as putative models for the empty fully formed pocket L. The H-bonds between lenalidomide and cereblon are shown as cyan dashed lines. Similar structural properties are apparent between the mouse and human orthologs, including high sequence identity among the residues lining the pocket.

Our overall findings are consistent with the following mechanism by which binding between cereblon and its cognate substrates is switched on and off in the presence and absence of bound degraders (Figure 26):

1) The cereblon population is tipped toward the substrate binding-<u>incompetent</u> form in the degrader-unbound state, corresponding to the fully collapsed and partially formed states of the degrader pocket, in which the interfacial H-bond groups needed for low cost desolvation of the interface are <u>misaligned</u> (promoting a non-physiologically slow k_1_) (Figures 27A and B).
2) The cereblon population is tipped toward the substrate binding-<u>competent</u> form in the degrader-bound state, in which the aforementioned H-bond groups are fully aligned (promoting a physiologically meaningful k_1_) (Figures 27C and D).

**Figure 26.**
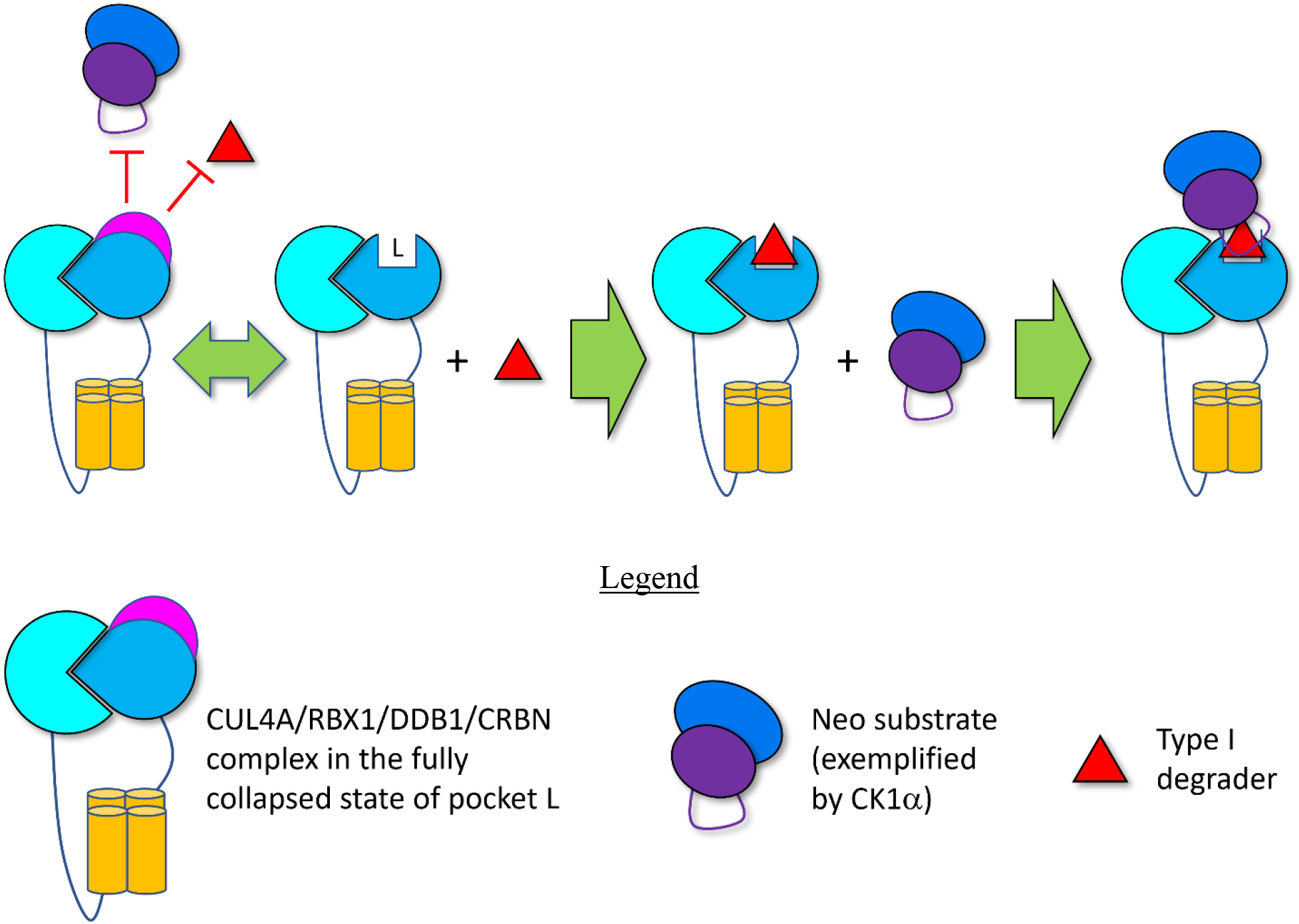
Cartoon depicting cereblon (medium blue tear dropped circle) in the CUL4A/ RBX1/DDB1/CRBN complex (described in the Legend), which we postulate is distributed between the fully collapsed and partially formed states of pocket L (double-headed arrow) in the absence of a bound degrader (red triangle). We hypothesize that the substrate binding surface of cereblon (magenta region) is misaligned for substrate binding in the collapsed degrader-unbound state of pocket L and aligned in the degrader-bound state. We further postulate that the major fraction of cereblon exists in the fully collapsed, substrate binding incompetent state of pocket L, which is shifted to the substrate binding competent form upon degrader binding.

**Figure 27.**
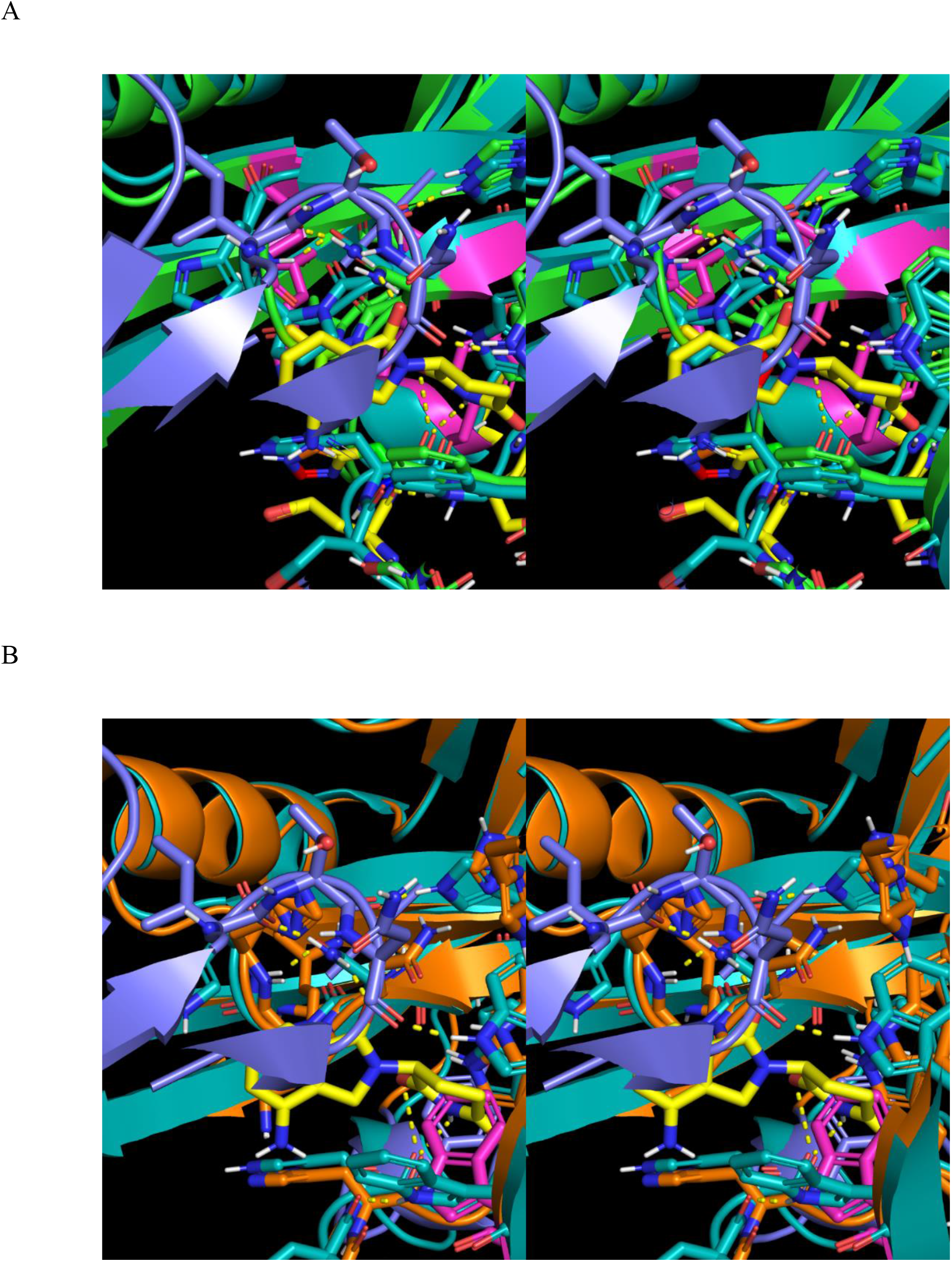
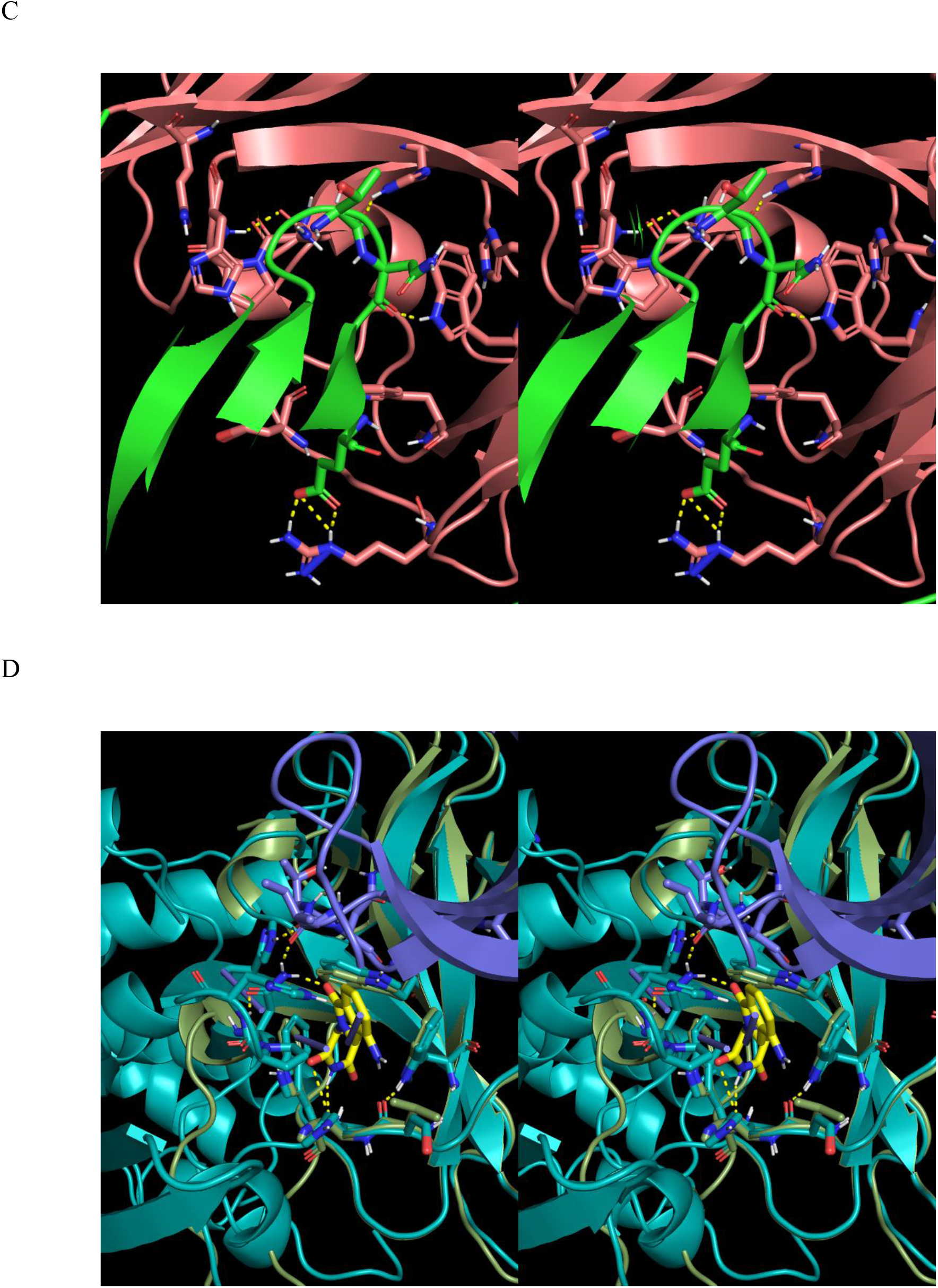
(A) Zoomed-in stereo view of the misaligned interfacial H-bond groups in the degrader- unbound structure A_closed (green) relative to the aligned groups in the degrader-bound structure F (cyan). (B) Zoomed-in stereo view of the misaligned interfacial H-bond groups in A_closed’ (orange) relative to the aligned groups in structure F (cyan). (C) Zoomed-in stereo view of the aligned interfacial H-bond groups in the degrader-unbound structure B. Pre-existing H-bonds in the starting structure (derived from 5FQD) are retained by AMBER, despite the absence of bound lenalidomide, which further supports our claim that the solute H-bond energy contribution is overweighted in the force-field. (D) Zoomed-in stereo view of the interfacial H-bond groups in the degrader-unbound structure H (olive green) and degrader-bound structure C (cyan). The backbone and aromatic side chains lining the degrader pocket of this structure are well-aligned with those in structure C (cyan) but not the substrate H-bond groups.

We hypothesize that the open/rotated CULT domain of cereblon (structures A_open and G) plays a key functional role in substrate ubiquitylation, as follows (Figure 28):

1) Degrader association triggers the release of the <u>N-terminal peptide</u> from the CULT domain that acts as a putative “latch” needed to stabilize the closed form observed in 5FQD. This is followed by dissociation of the CULT and N-terminal domains, thereby forming the open state observed in 6H0F and addressed below (noting that 5FQD may be stabilized in the closed state by crystal contacts). We made no attempt to simulate this process.
2) Substrate association ensues (noting that substrate binding prior to CULT rotation cannot be ruled out, and the two possibilities are non-mutually exclusive).
3) Ubiquitylation of the growing poly-Ub chain is facilitated by dynamic rotation of the ternary CULT-degrader-substrate complex.

Pocket T exists solely in the time-averaged structures C, F, and A_closed (DDB1 omitted), as well as A_closed’ (DDB1 included), but is occupied by His332 in structures D, C’ (DDB1 included) and C” (N-terminal peptide and DDB1 omitted) (Figure 29). We postulate that the closed → open state transition (captured in 6H0F and structure A_open) depends on degrader-induced release of the N-terminal peptide via desolvation of pocket T by His332.

**Figure 28.**
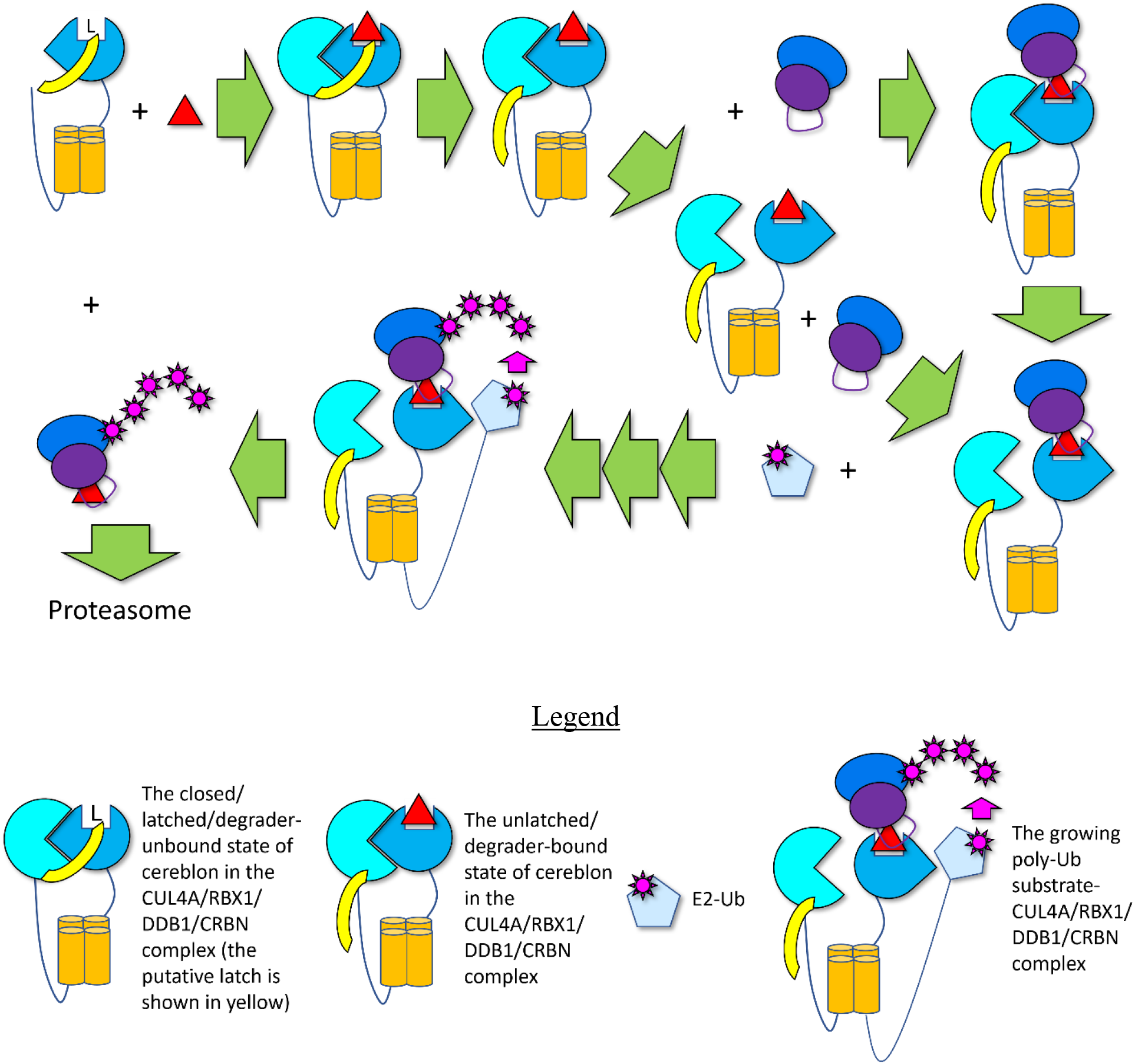
Cartoon depicting the putative CUL4A/RBX1/DDB1/CRBN enzyme cycle (described in the Legend), in which the degrader (red triangle) binds to the closed form of cereblon (the cyan and blue shapes and orange helical bundle), resulting in the release of the N-terminal peptide “latch” (yellow shape), followed by dissociation of the N-terminal and CULT domains. Substrates may associate to the closed and/or open forms of cereblon, which are then poly-ubiquitylated (magenta shapes) by the E2 subunit (light blue pentagon), followed by release of the poly- ubiquitylated product.

**Figure 29.**
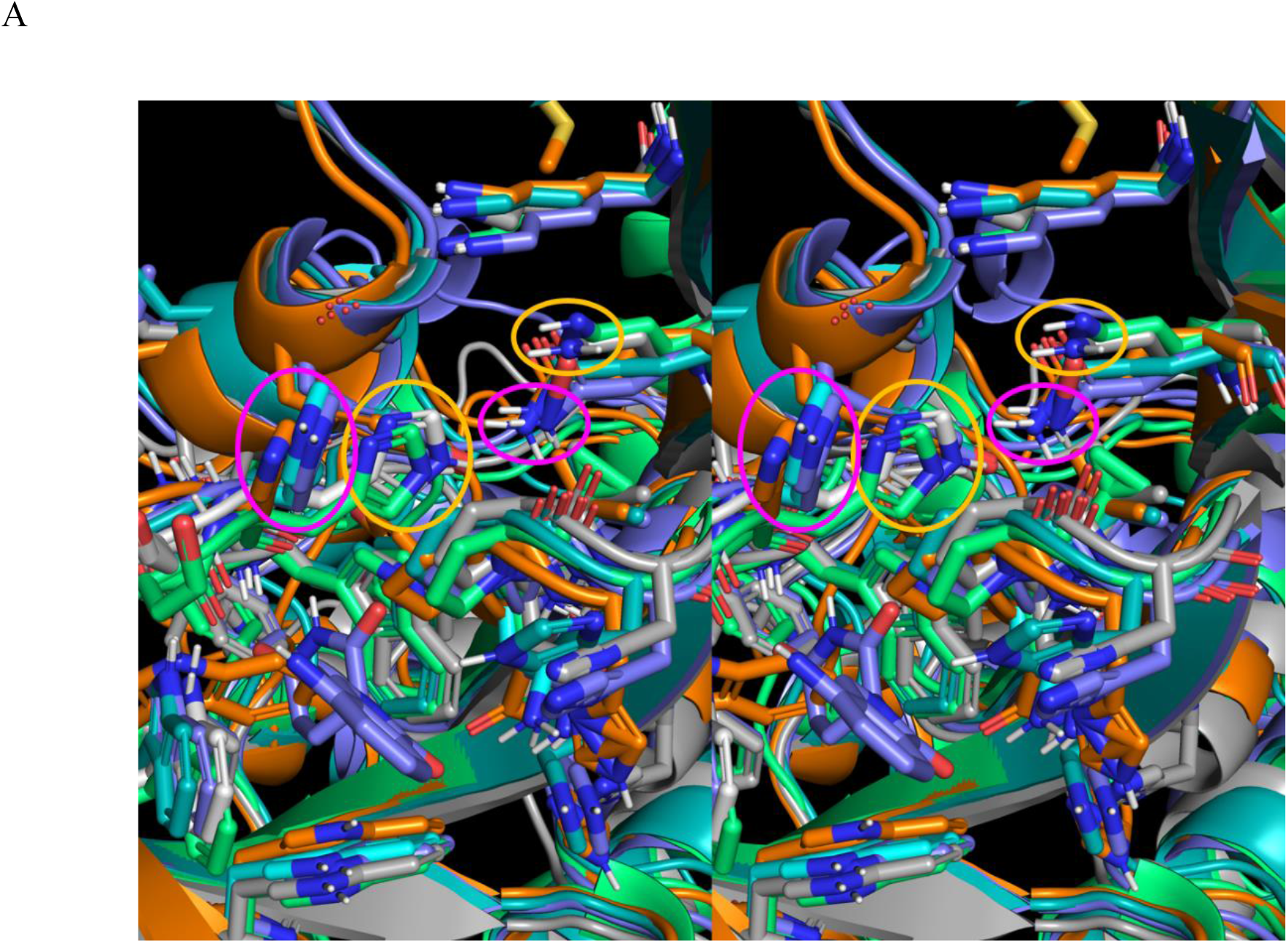
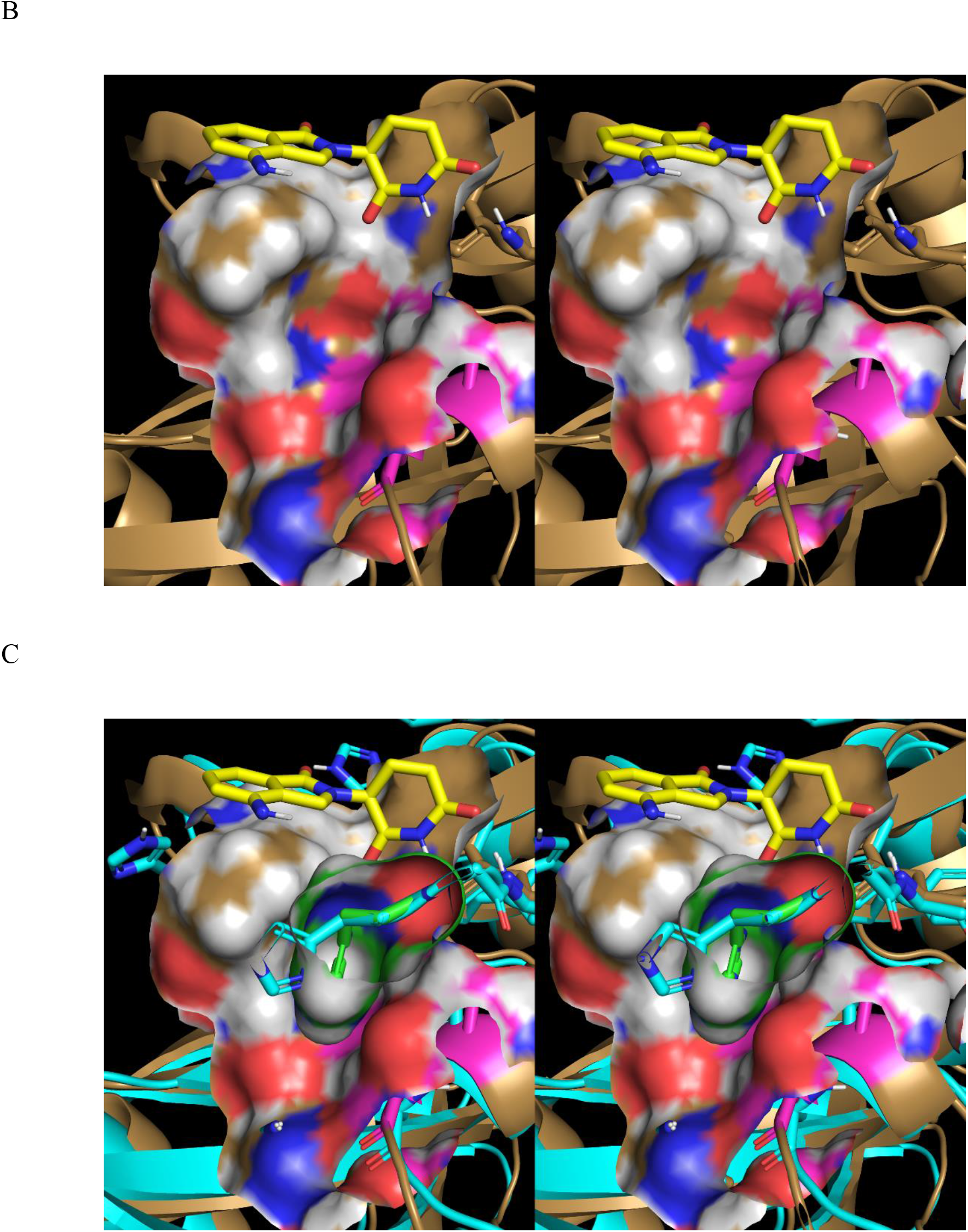
(A) Stereo view of His332, which transitions from the interior of pocket T in structures C’ (mint green cartoon) and C” (white cartoon) to the exterior in structures C (purple cartoon) and F (cyan cartoon). Occluding versus non-occluding His332 side chain positions are circled in yellow and magenta, respectively (where the pocket overlaps with the yellow circled His side chain). The His332 side chain position is correlated with that of Gln54 located on the N-terminal domain (the position correlated with the His332-occupied form is circled in yellow, and that with His332 unoccupied form is circled in magenta). (B) Stereo view of pocket T in the His332- occupied state in structure A_closed” with the His332 side chain omitted for clarity and lenalidomide from structure F shown for reference (yellow). Pocket T resides at the three-way interface between the N-terminal peptide (magenta cartoon and surface), N-terminal domain, and CULT domain. (C) Same as B, except in the His332-occupied form of pocket T (green sticks and surface). Lenalidomide (yellow sticks) and His332 (cyan sticks) in pocket L of structure F are shown for reference. We hypothesize that the interface is disrupted when pocket T is vacated by His332 in response to degrader binding, which is driven by desolvation of the pocket by the southern glutarimide O of the degrader.

### The solvation-powered intra- and intermolecular dynamics of cereblon

Next, we studied the solvation fields powering the intra- and intermolecular dynamics of cereblon, and specifically:

1) The solvation free energy contributions to the substrate and degrader association and dissociation barriers underlying k_1_, k_-1_ and k_on_, k_off_, respectively, in the context of the time-averaged structures and solvation fields of cereblon complexed with DDB1:

a) Modeled unbound cereblon (structures A_closed, A_closed’, and A_open, and H) and CK1α (structure E).
b) The hypothetical binary cereblon-CK1α (structure B).
c) Modeled binary cereblon-lenalidomide (structures C and C’).
d) The ternary cereblon-lenalidomide-CK1α (structure F) complexes (5FQ1).
2) Complementarity between the solvation field of the degrader pocket (structures A_closed and H) and those of thalidomide, lenalidomide, and uridine.
3) The putative mechanism by which dissociation of the N-terminal peptide (observed in 6H0F) is powered by the degrader-induced release of solvation free energy stored within pocket T.
4) The internal/buried solvation within cereblon and its role in facilitating and powering the intramolecular state transitions of the protein (explained in general terms in the Introduction).

According to Biodynamics theory, association barriers, in general, are contributed principally by the <u>desolvation costs</u> of the <u>associating</u> partners, whereas dissociation barriers are contributed principally by the <u>resolvation costs</u> of the <u>dissociating</u> partners (as explained in previous sections of this work and in our previous works^2–4)^. Fast association depends on high binding partner concentrations and/or fast k_1_, which in turn, depends on optimal H-bond replacements of H-bond enriched interfacial solvating water by inter-partner H-bonds. Slow dissociation depends on significant desolvation of H-bond depleted solvation during association (i.e., depending on the existence of such solvation on either partner, and the extent to which such solvation is sterically overlapped in the bound state). We assume that the cognate substrates of each specific cereblon- degrader complex:

1) Are sterically matched and tightly packed so as to access k_-1_-slowing H-bond <u>depleted</u> solvation residing at the protein surface, deep within the solvation field.
2) Contain sufficient amounts of H-bond depleted solvation that is mutually expelled from the binding interface (conveying a physiologically meaningful t_1/2_).
3) Project spatially matched polar groups into the interface.

We calculated the solvation fields within the binding interface of CK1α and cereblon separately, along with those of the hypothetical binary cereblon-CK1α (structure B) and ternary (structure F) complexes using WATMD. Replacement of water in the <u>ultra-low occupancy voxels</u> (ULOVs) of CK1α by non-polar groups contributed by the cereblon-lenalidomide complex and of cereblon by non-polar groups contributed by CK1α is shown in Figures 30A and B, respectively. H-bonds formed among the aligned partners replace those of their solvating water, thereby lowering the desolvation cost of the interface (Figure 30C).

**Figure 30.**
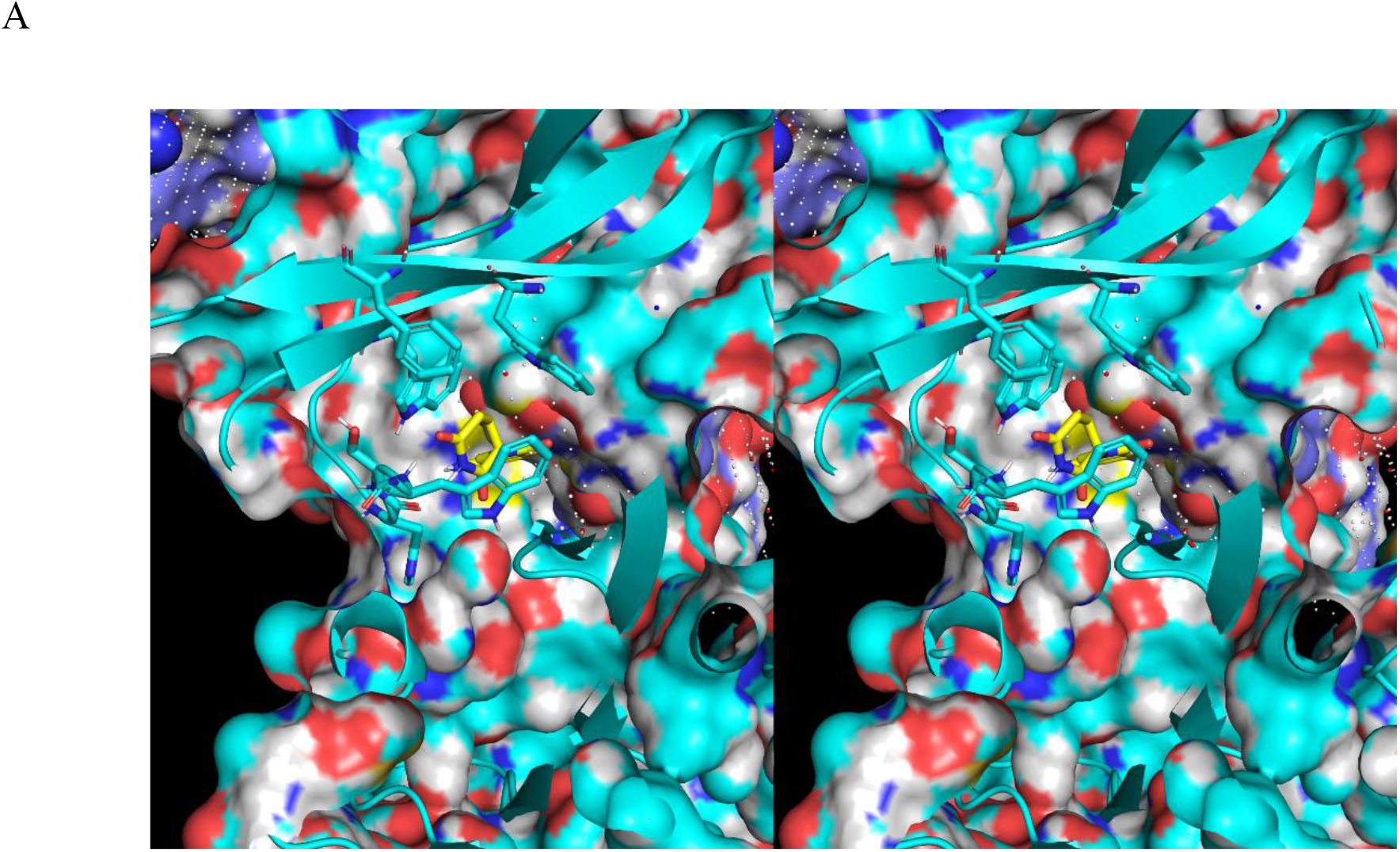
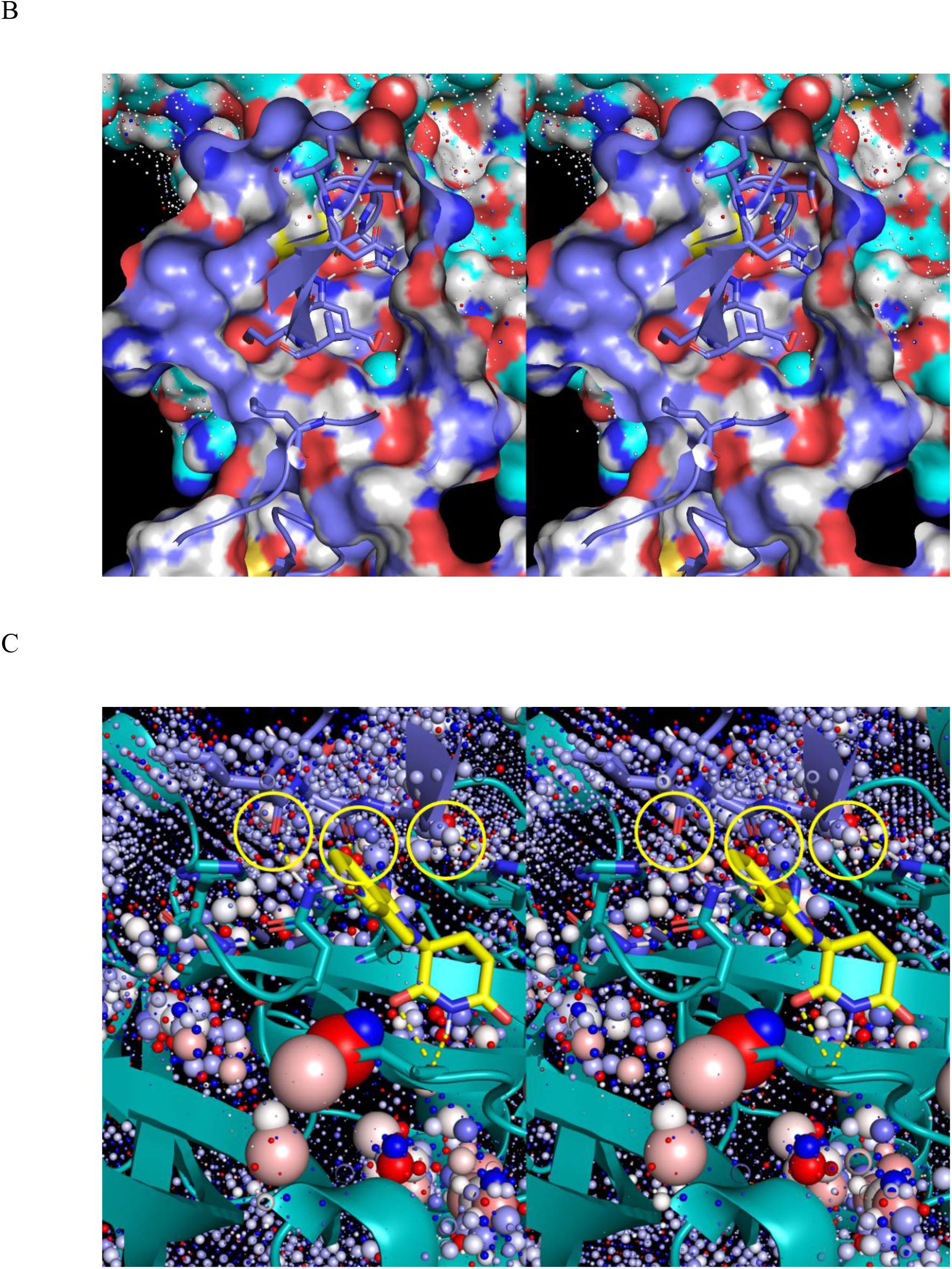
(A) Stereo view of the ultra-low occupancy voxels (ULOVs: small white spheres) on the surface of CK1α (blue cartoon) corresponding to H-bond depleted solvation that is replaced by non-polar groups during cereblon-lenalidomide (cyan cartoon) association (viewed from the interior of the cereblon-lenalidomide complex looking toward the external surface of CK1α). (B) Same as A, except the ULOVs on the surface of the cereblon-lenalidomide complex corresponding to H-bond depleted solvation replaced by non-polar CK1α groups (viewed from the interior of CK1α, looking toward the external surface of cereblon-lenalidomide). (C) Stereo view of high occupancy voxels (HOVs: blue-white-red color spectrum coded according to donor-no preference- acceptor preference) corresponding to H-bond enriched solvation of cereblon Asn305, His311, and Trp354 desolvated by CK1α (circled in yellow) (noting that the intra-cereblon and cereblon- lenalidomide H-bonds were necessarily desolvated prior to CK1α binding). The small voxel sizes corresponding to this water reflect relatively mild H-bond enrichment (which is potentially underestimated by WATMD), suggesting that the interfacial desolvation cost is relatively low and k_1_ is therefore relatively fast. The H-bonds of this water are replaced by the cereblon-CK1α H- bonds shown in Figure 22A.

It then follows that the substrate binding-competent state of cereblon is stabilized by degrader-induced desolvation of pocket L, and further stabilized by two remote H-bond enriched solvation positions (corresponding to the HOVs in Figures 31A and B), one of which persists in the CK1α- bound state, that likely further stabilizes that state (Figure 31C).

**Figure 31.**
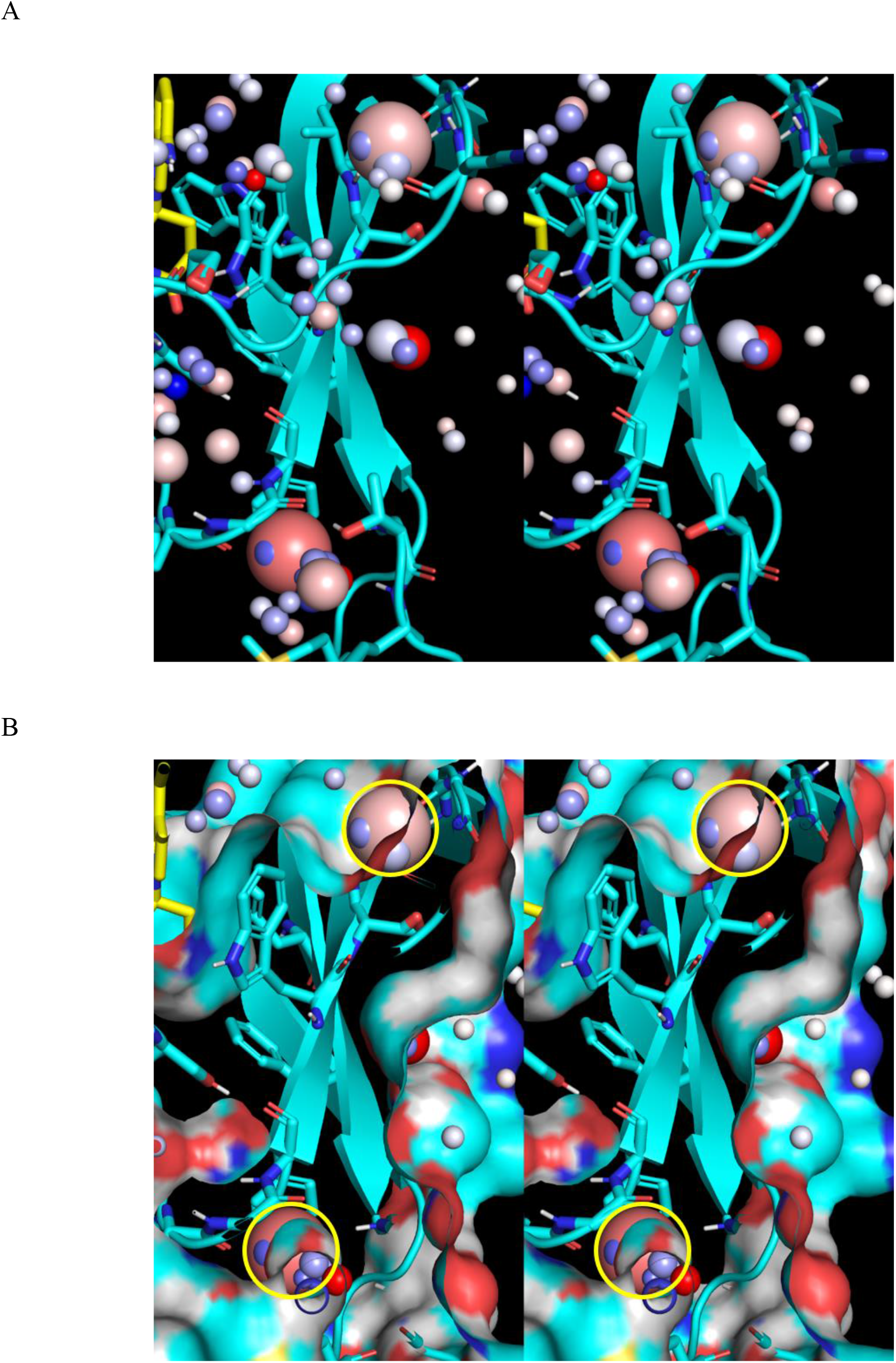
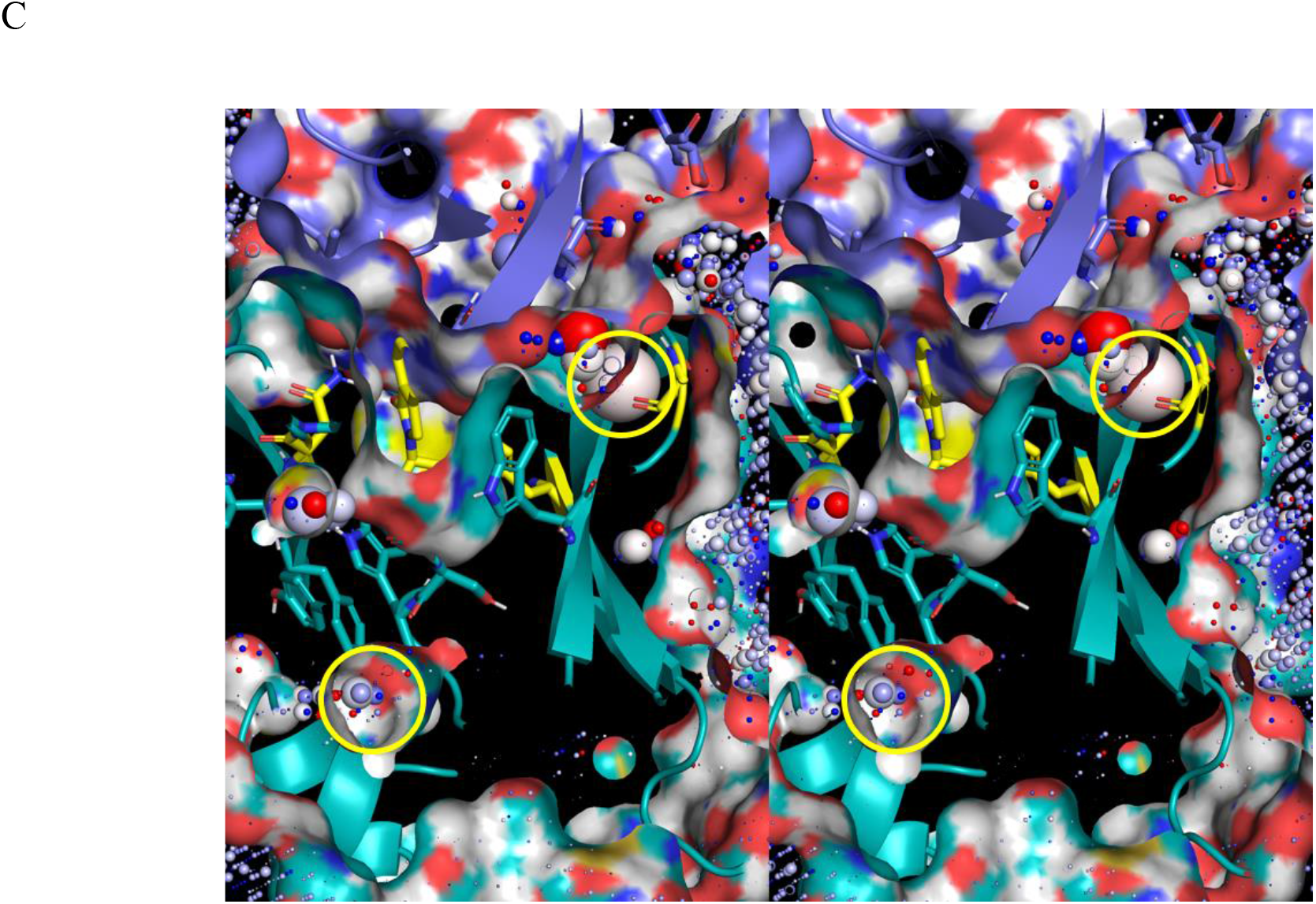
(A) Zoomed-in stereo view of two HOVs (circled in yellow) present in structure C, but not structure A. This water likely helps to stabilize the degrader-bound state of the protein. (B) Same as A, except showing the protein surface. The two HOVs reside at the protein surface, and therefore represent H-bond enriched, rather than buried/trapped water. (C) Same as A, except for the ternary structure F. The upper voxel resides within the CK1α-cereblon interface (purple and cyan surfaces, respectively), whereas the lower one is entirely absent.

Next, we analyzed the WATMD results for the empty fully formed state of pocket L (structures A_closed” and H). The ability of pocket L to switch between the substrate binding-competent and incompetent forms of cereblon depends on instability of the <u>empty pocket</u> and substrate binding surface, corresponding to disrupted shape and interfacial H-bond group complementarity vis-à-vis cognate substrate(s). We postulate that the solvation of pocket L (Figure 32A) resides within a Goldilocks zone of unfavorability (both enthalpically and entropically), such that the population is distributed dynamically between the collapsed/auto-desolvated and fully formed/solvated states (states 1 and 2, respectively), as follows:

1) State 1 (the degrader and substrate binding-<u>incompetent</u> state): desolvation of the pocket via inward rotation of Trp340 (which is favorable) is coupled directly to the <u>loss</u> of H-bond enriched solvation in the peri-pocket region (which is unfavorable) (Figure 32B).
2) State 2 (the degrader and substrate binding-<u>competent</u> state): resolvation of the pocket via outward rotation of Trp340 (which is unfavorable) is driven by the accompanying <u>gain</u> of H-bond enriched solvation in the peri-pocket region (which is favorable). As such, <u>the substrate-binding competent/substrate-bound populations build on the coattails of degrader association</u>.

**Figure 32.**
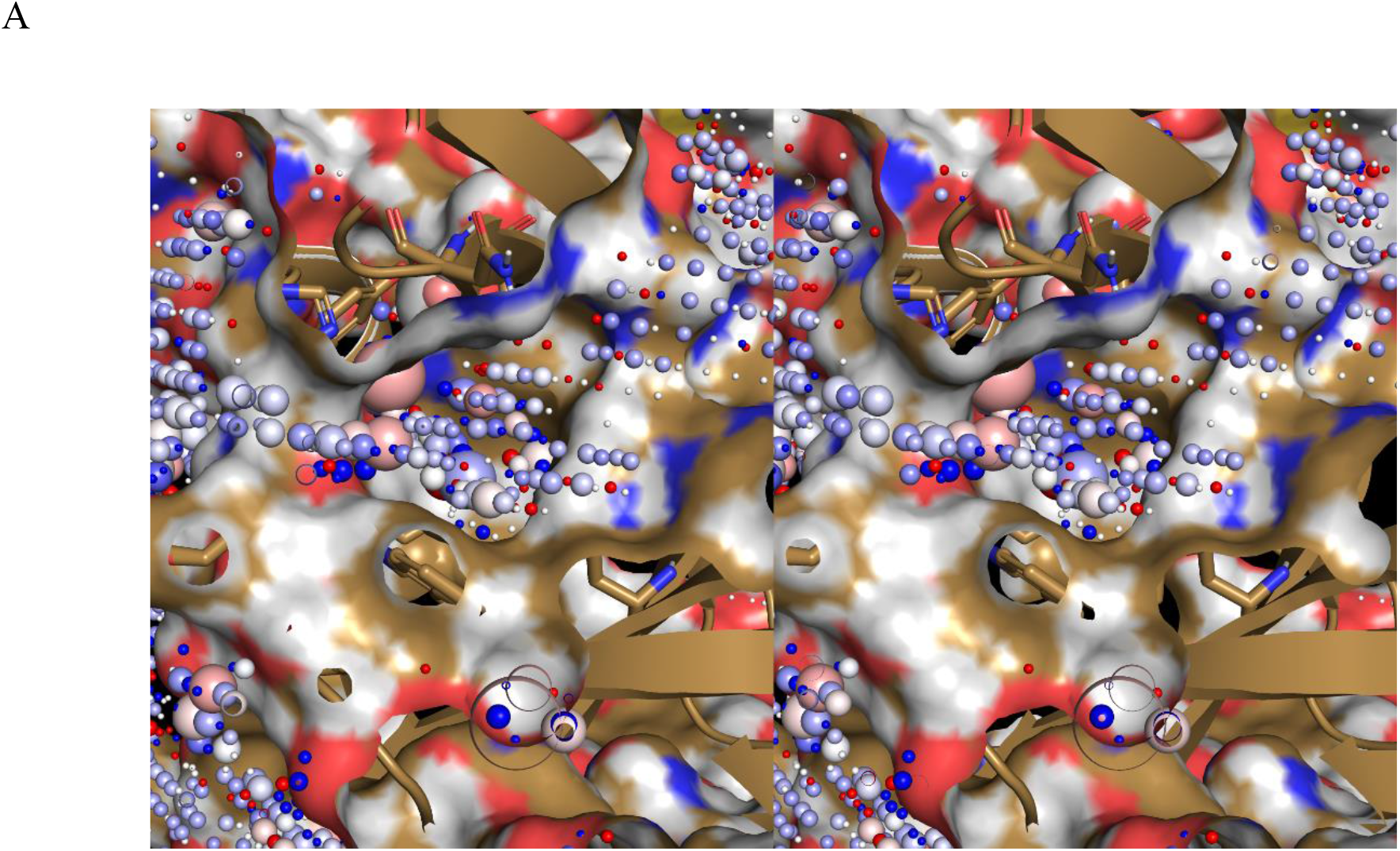
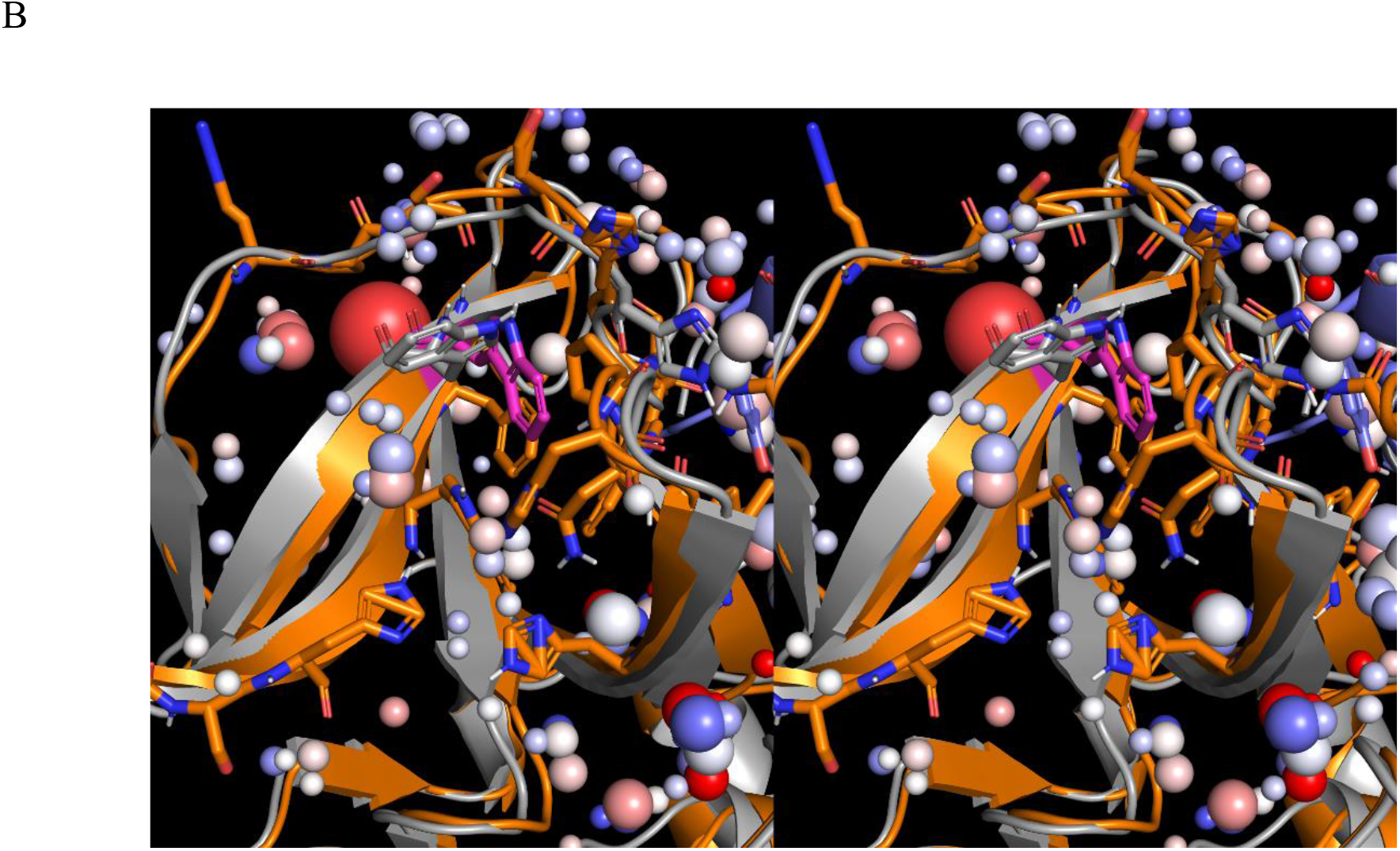
(A) Stereo view of the ULOVs and HOVs within the fully formed pocket L of structure A_closed’. This solvation exhibits a range of H-bond depletion (small to large spheres) relative to bulk solvent (see text). (B) Favorable auto-desolvation of pocket L in structure A_closed’ (in which pocket L is fully collapsed) is achieved via transient rotation of Trp340 into the pocket (white stick from structure C versus magenta stick from structure A_closed’). This rearrangement is putatively counter-balanced by the unfavorable loss of H-bond enriched solvation from the voxel positioned directly behind the Trp340 backbone (red sphere), resulting in conformational “frustration”.

A significant number of ULOVs reside within the fully formed (Figures 33A and 34A) and partially collapsed states of pocket L (structures A_closed” and A_closed, respectively), which is consistent with the highly aromatic composition of the pocket. The higher occupancy voxels within the pocket (Figures 33B and 34B) correspond to solvating water that is H-bonded to a small number of protein H-bond groups (Figure 33C) (well below the enrichment threshold, recalling that bulk solvent contains ∼3.5 moderately strong H-bonds per water at zero entropic cost). The voxel configuration is consistent with fast k_on_ and slow k_off_ of lenalidomide under native conditions (corresponding to low desolvation and high resolvation costs, respectively), the correct measurement of which may be hampered by the hypothesized non-continuous availability of the pocket. Fewer HOVs are present in structure H, likely due to the shallower nature of the pocket (Figures 34B and C).

**Figure 33.**
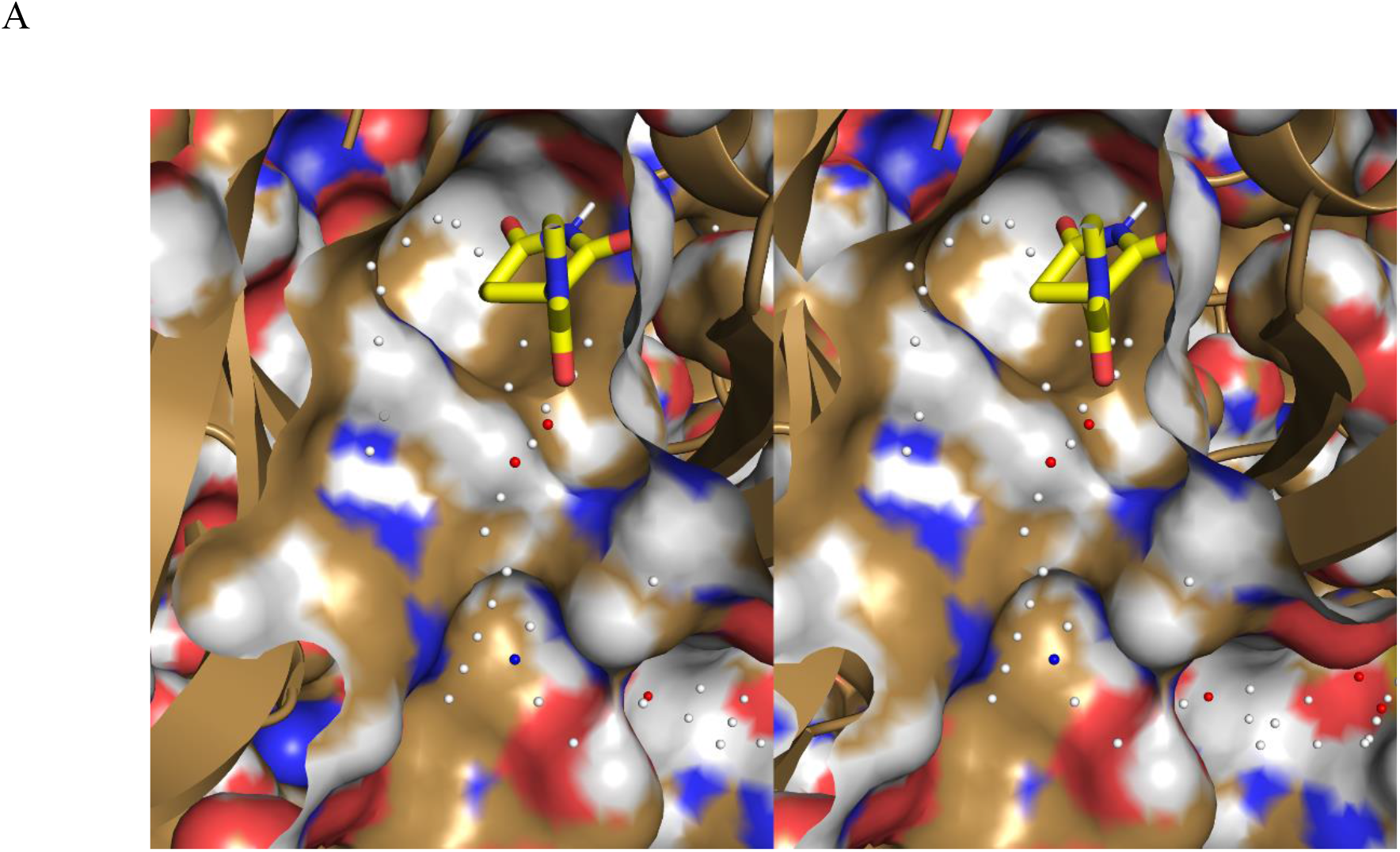
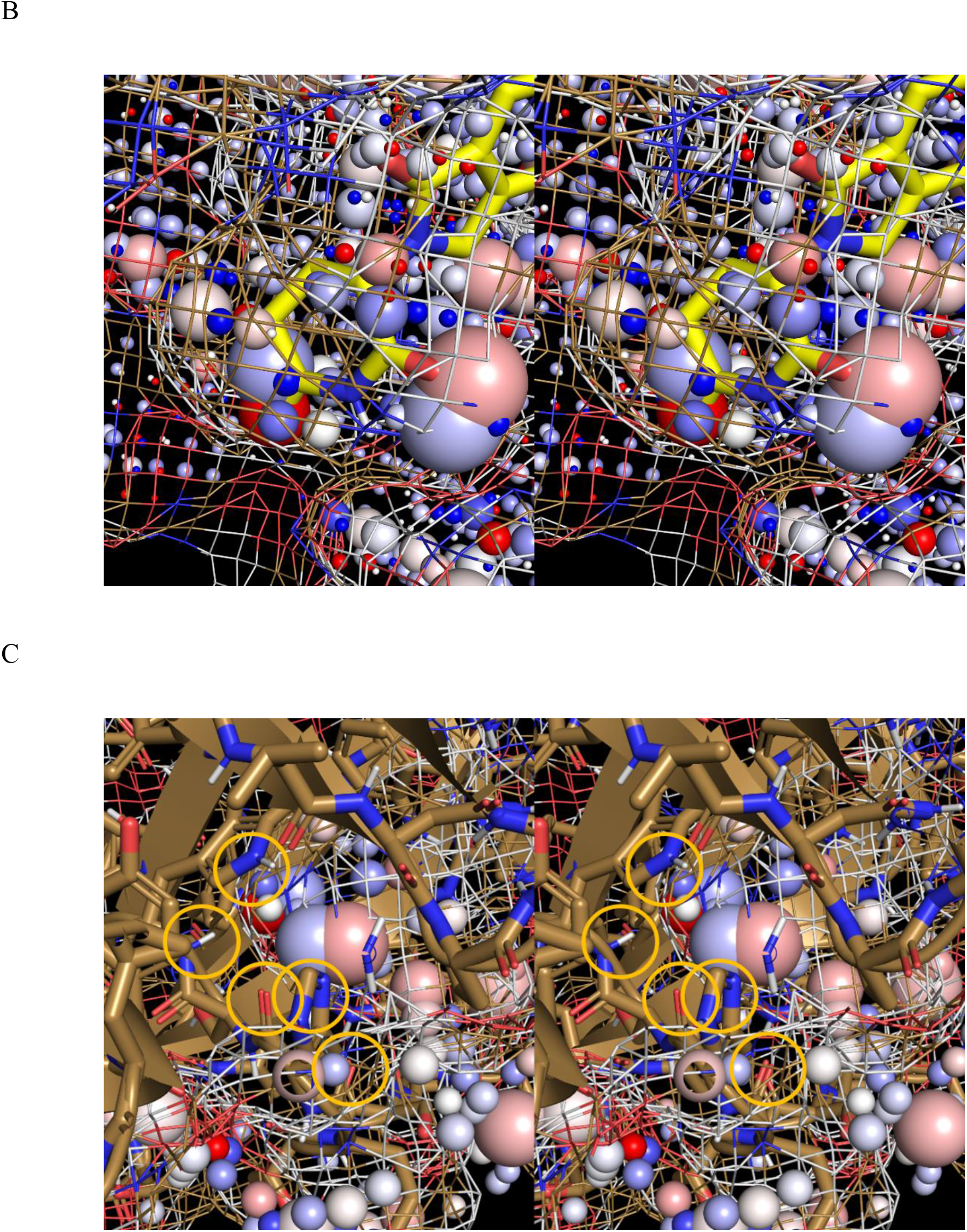
Stereo views of the voxels within the modeled fully formed pocket L in structure A_closed” (tan solid surface) (shown as spheres described in Materials and methods). (A) Zoomed-in view of the ULOVs in the pocket overlaid on lenalidomide from structure C (yellow sticks). (B) Same as A, except showing the higher occupancy voxels which are organized in a ring-like fashion that mirrors the glutarimide ring of the degrader (such that the glutarimidine oxygens overlap the two large voxel clusters, with the NH group residing between). Favorable desolvation of HOVs/H-bond enriched solvation is conveyed strictly by polar atoms, such as the glutarimide moiety of the ring, which is consistent with the lack of H-bond enriched solvation at these positions. (C) Same as B, except zooomed into the HOVs at the base of the pocket, showing the H-bond groups that partially stabilize the solvating water at these positions (circled in yellow).

**Figure 34.**
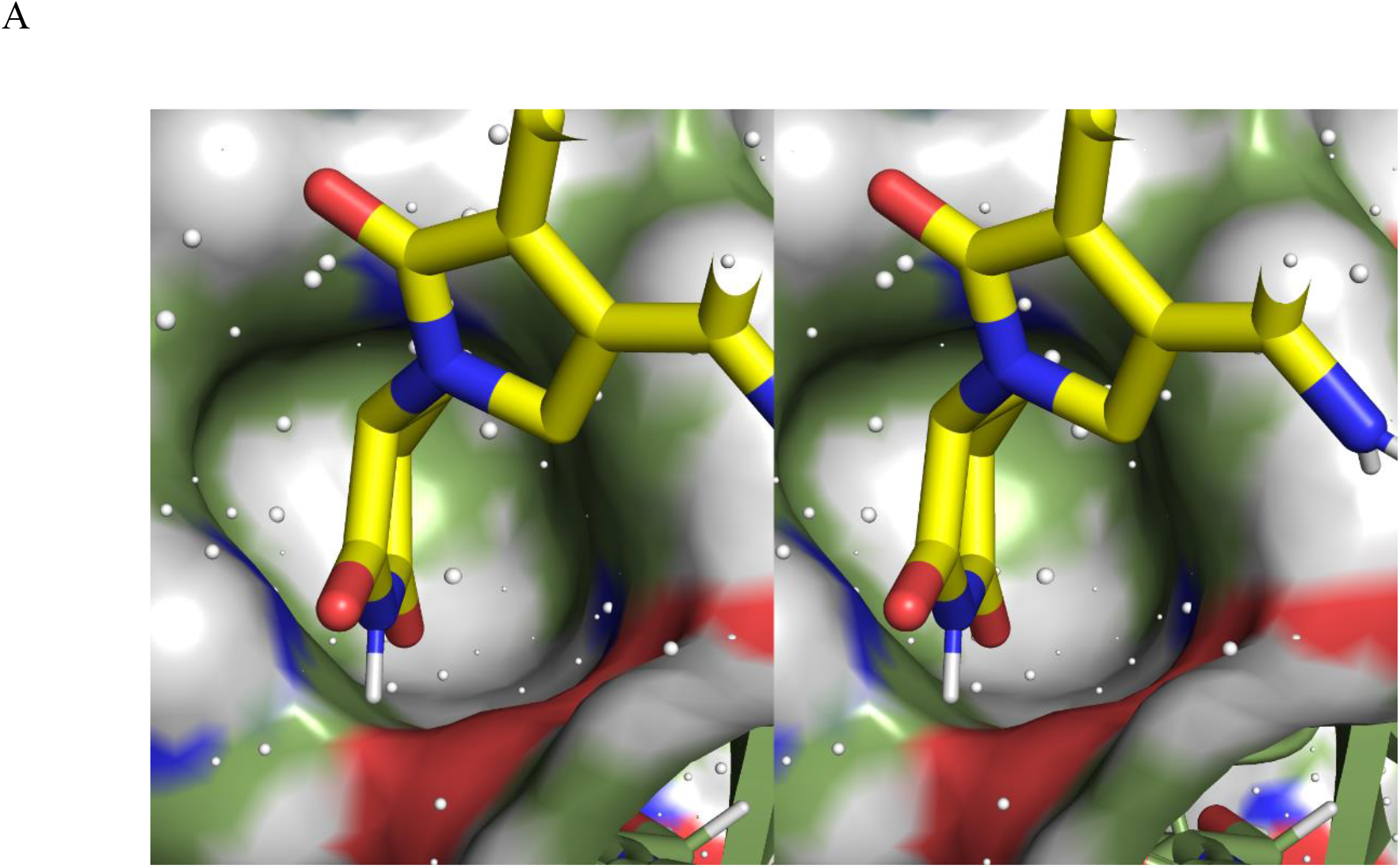
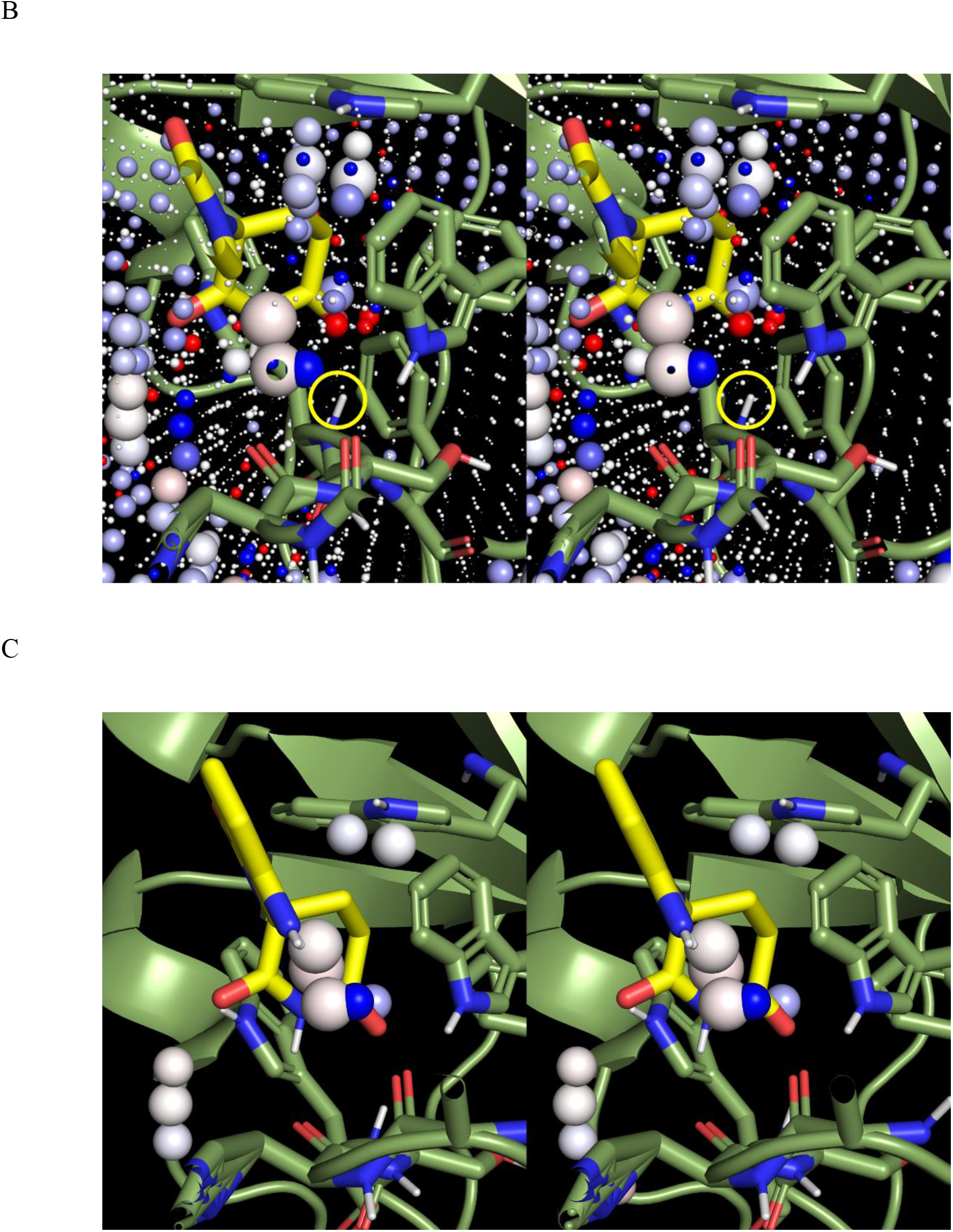
Zoomed-in stereo views of the voxels within the fully available pocket L of structure H. Lenalidomide (yellow) from structure C is shown for reference. (A) ULOVs corresponding to H-bond depleted solvation (white spheres) residing along the pocket surface. (B) Moderate occupancy voxels corresponding to slightly enriched solvation arising from H-bonding with the backbone NH of Trp64 (circled in yellow). The voxels are overlapped by one glutarimide O of lenalidomide, consistent with replacement of the occupying solvation by this group during association. (C) HOVs corresponding to H-bond enriched solvation arising from H-bonding with the backbone Os of Val61 and His62. The voxels are approximately overlapped by the exocyclic NH_2_ of lenalidomide, consistent with replacement of the occupying solvation by this group during association.

Next, we analyzed the LMW WATMD results for lenalidomide (Figure 35), thalidomide (Figure 36), and uridine (based on a docked model)) (Figure 37) and assessed the voxel complementarity with those of the fully formed pocket L in structure A_closed” (the results are described in the figure captions).

**Figure 35.**
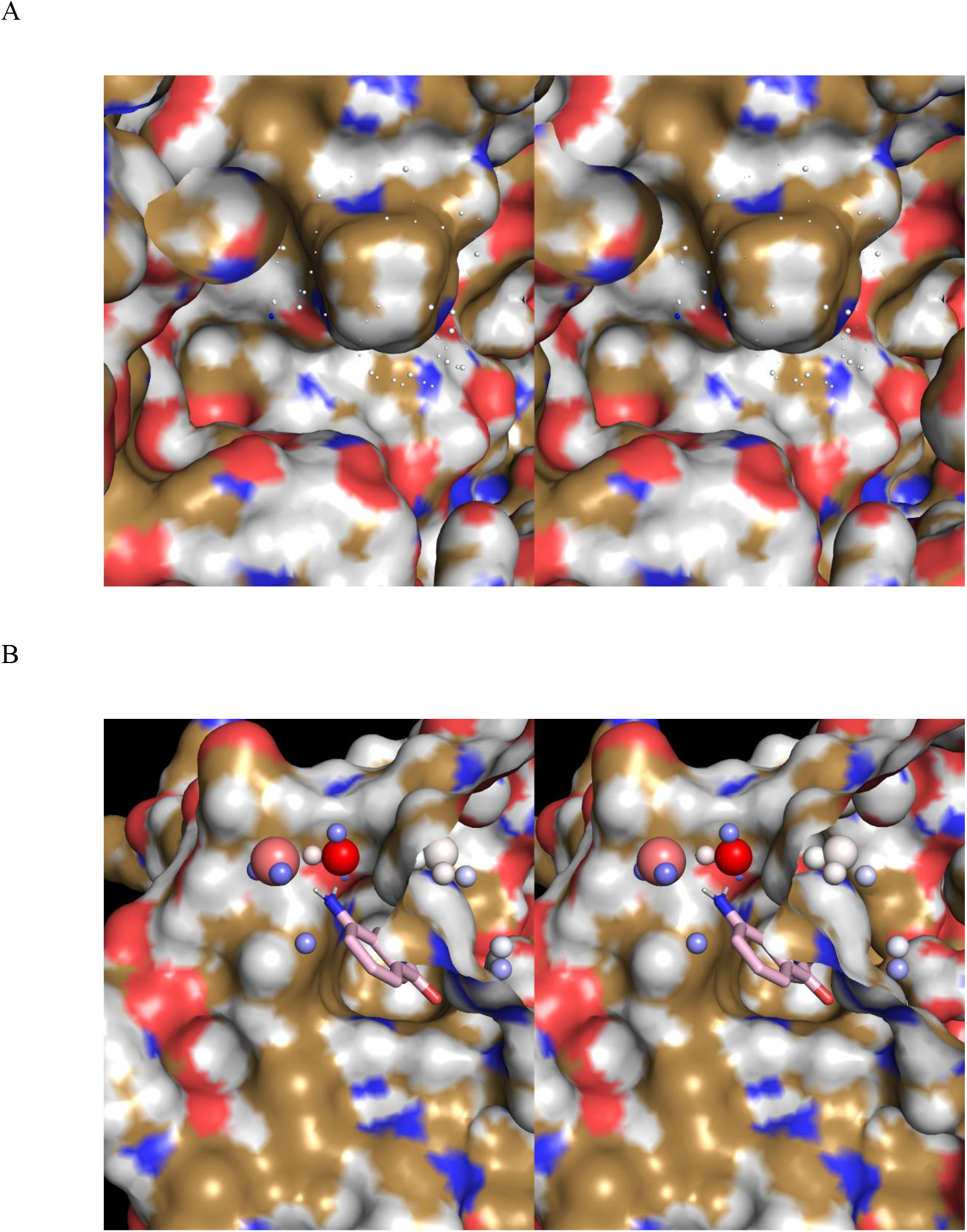
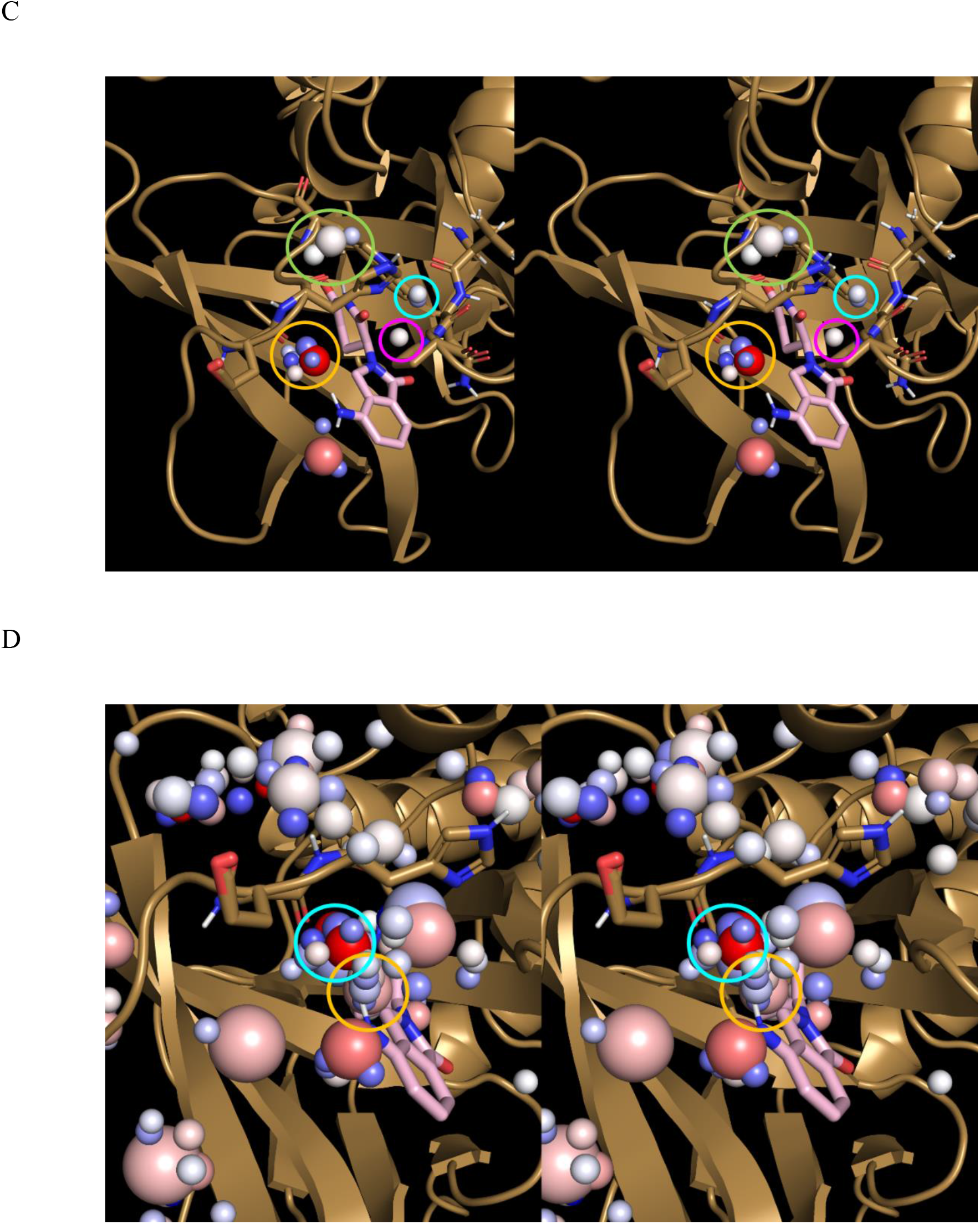
(A) Stereo view of the the LMW time-averaged lenalidomide structure (not visible) and ULOVs (white spheres) overlaid on structure A_closed”. The interior surface of structure A_closed” (looking from the DDB1 interface toward pocket L) is visible in the center of the figure. The solvation in these voxels is necessarily expelled by cereblon during lenalidomide association, contributing to slowed k_off_ of the degrader. (B) Stereo view of the LMW time-averaged lenalidomide structure (pink sticks) and HOVs (colored using a blue-white-red spectrum, as described in Materials and methods) overlaid on structure A_closed”. All but the large red voxel in the northeast corner are expelled during lenalidomide association, which putatively governs the degrader k_on_. (C) Stereo view of the LMW time-averaged lenalidomide structure and HOVs thereof, showing desolvation of those positions by the backbone O of Glu331 (circled in yellow), side chain of Trp334 and backbone O of His332 (circled in light green), side chain of Pro306 (circled in magenta), and side chain of Trp334 (circled in cyan). (D) Same as C, except including the HOVs around the backbone O of Glu331 of structure A_closed” (circled in yellow), together with those around the exocyclic amine of lenalidomide (circled in cyan). These positions are mutually desolvated by the protein and degrader during association.

**Figure 36.**
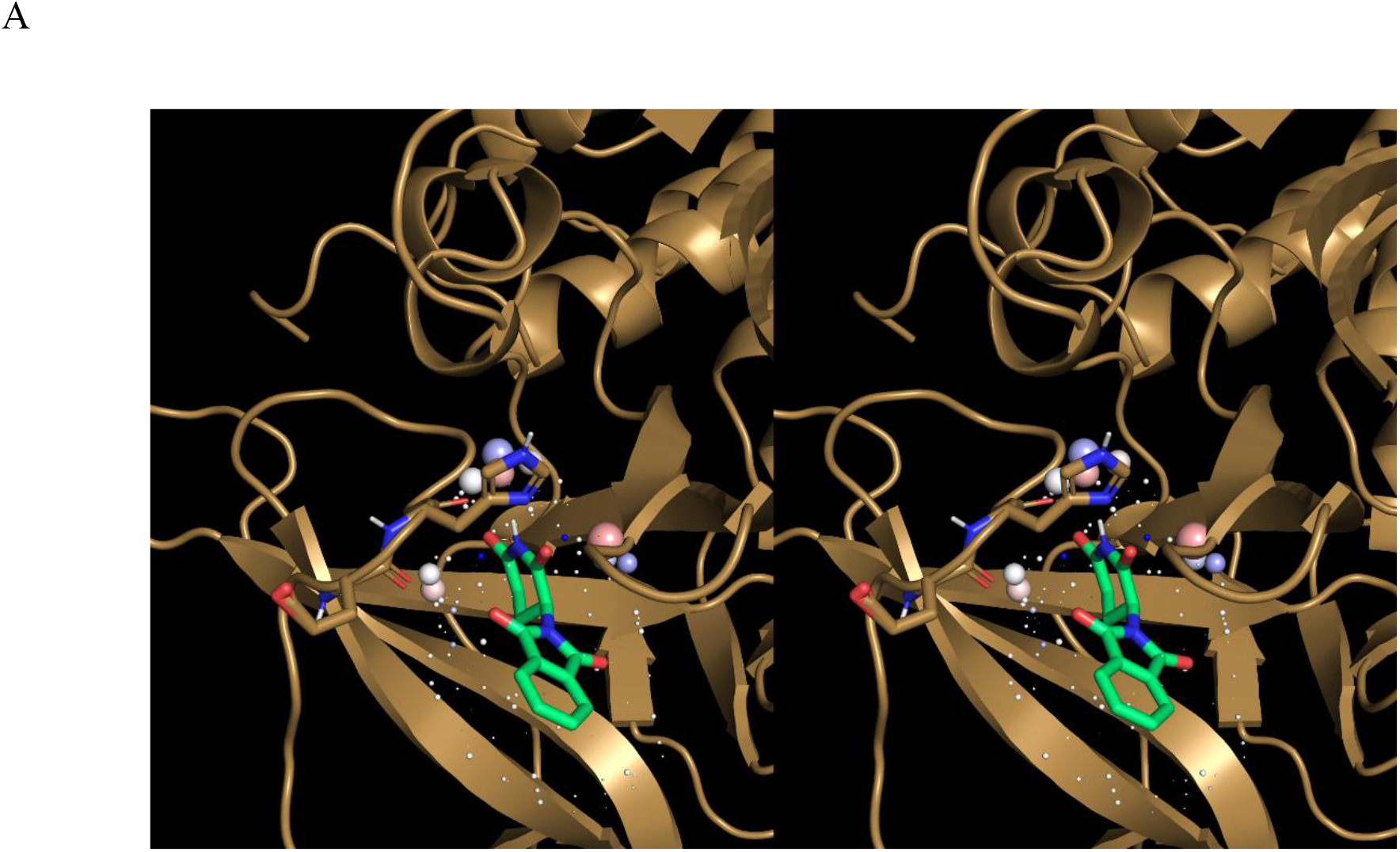
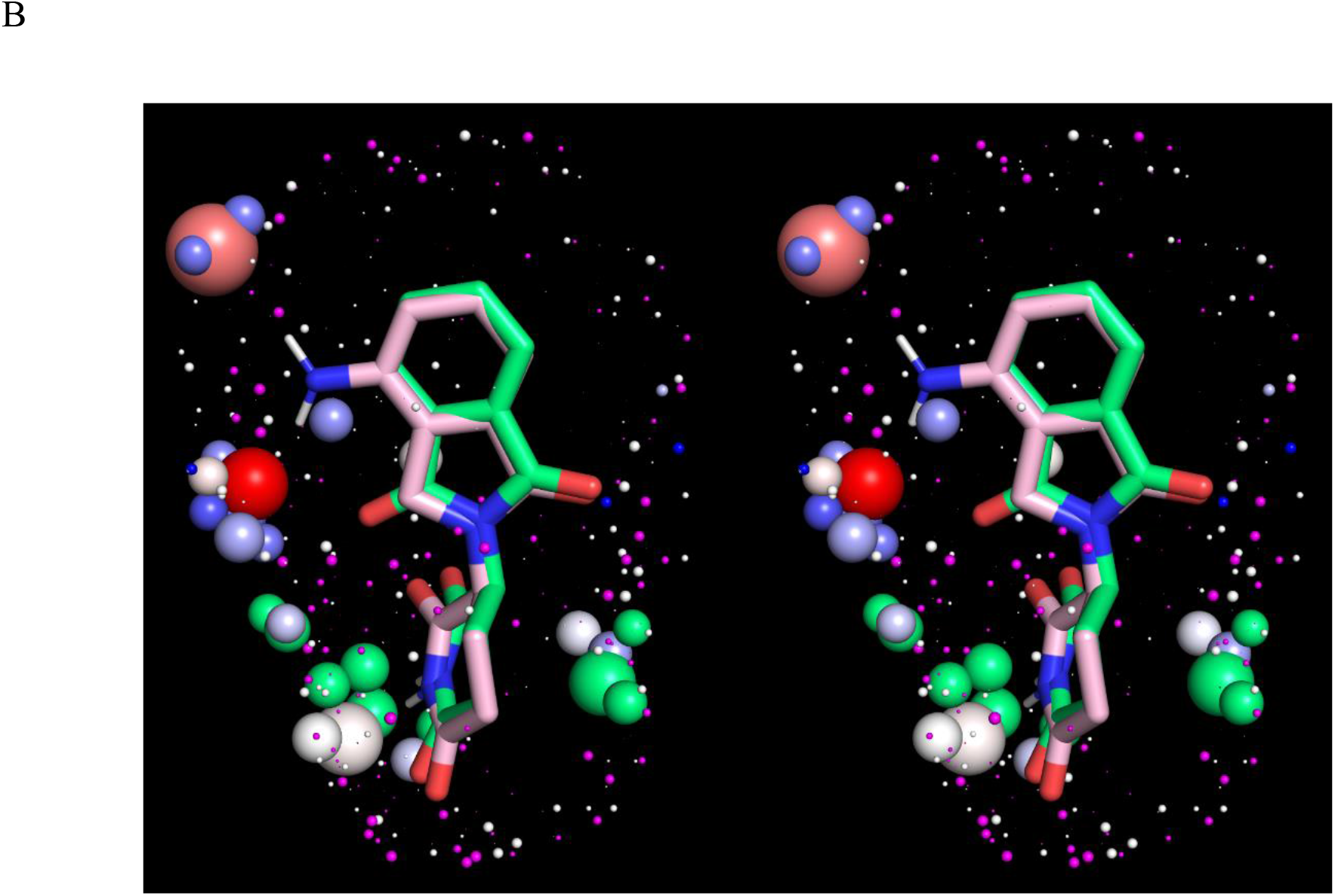
(A) Stereo view of the LMW time-averaged lenalidomide structure and the ULOVs and HOVs overlaid on structure A_closed”. (B) Stereo view of the time-averaged lenalidomide structure (pink sticks) and voxels (colored as in A) overlaid on the voxels (HOVs and ULOVs colored in mint green and magenta, respectively) and time-averaged thalidomide structure (mint green sticks). The voxels of the two degraders differ mainly around the exocyclic amine position.

**Figure 37.**
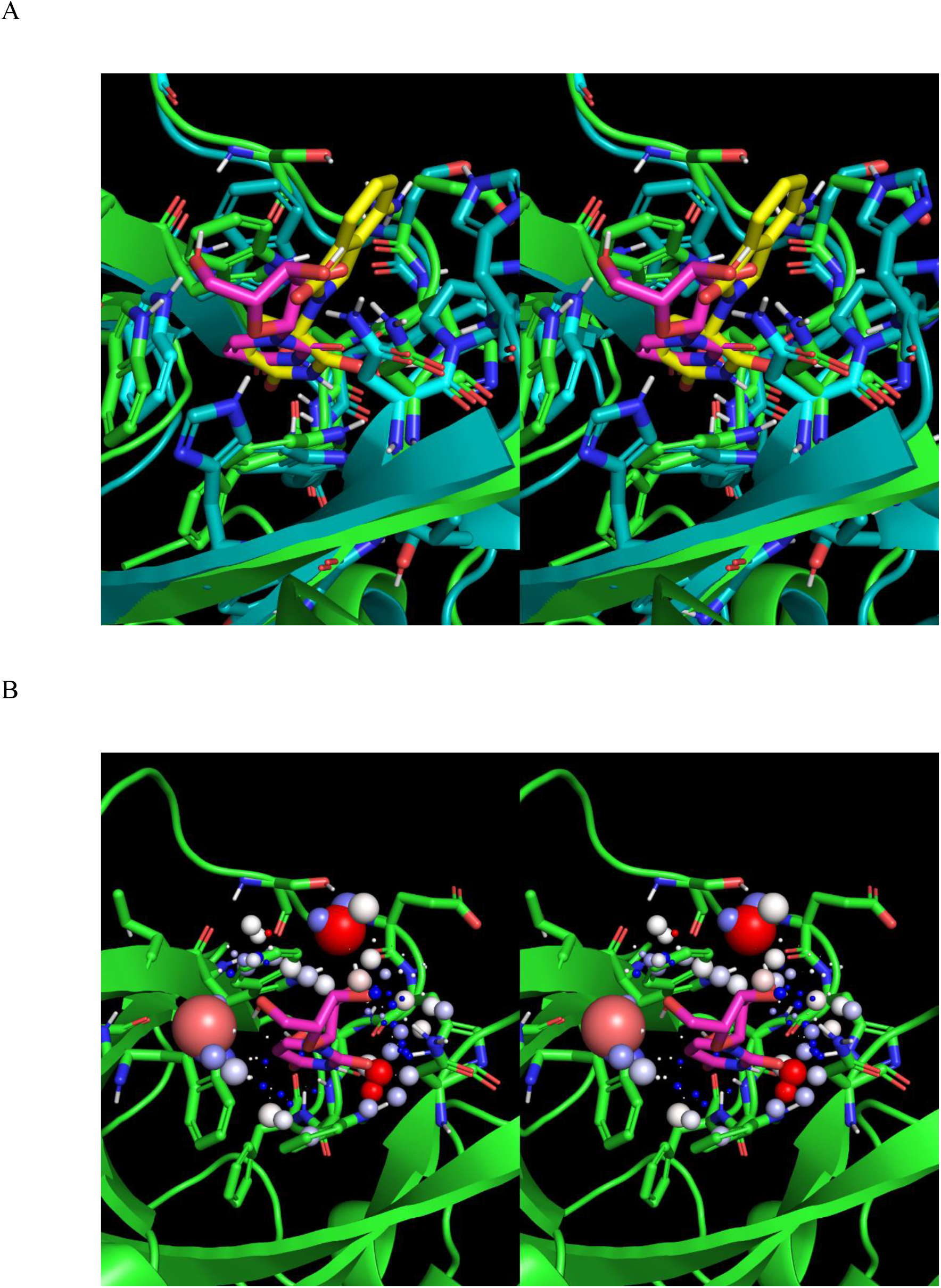
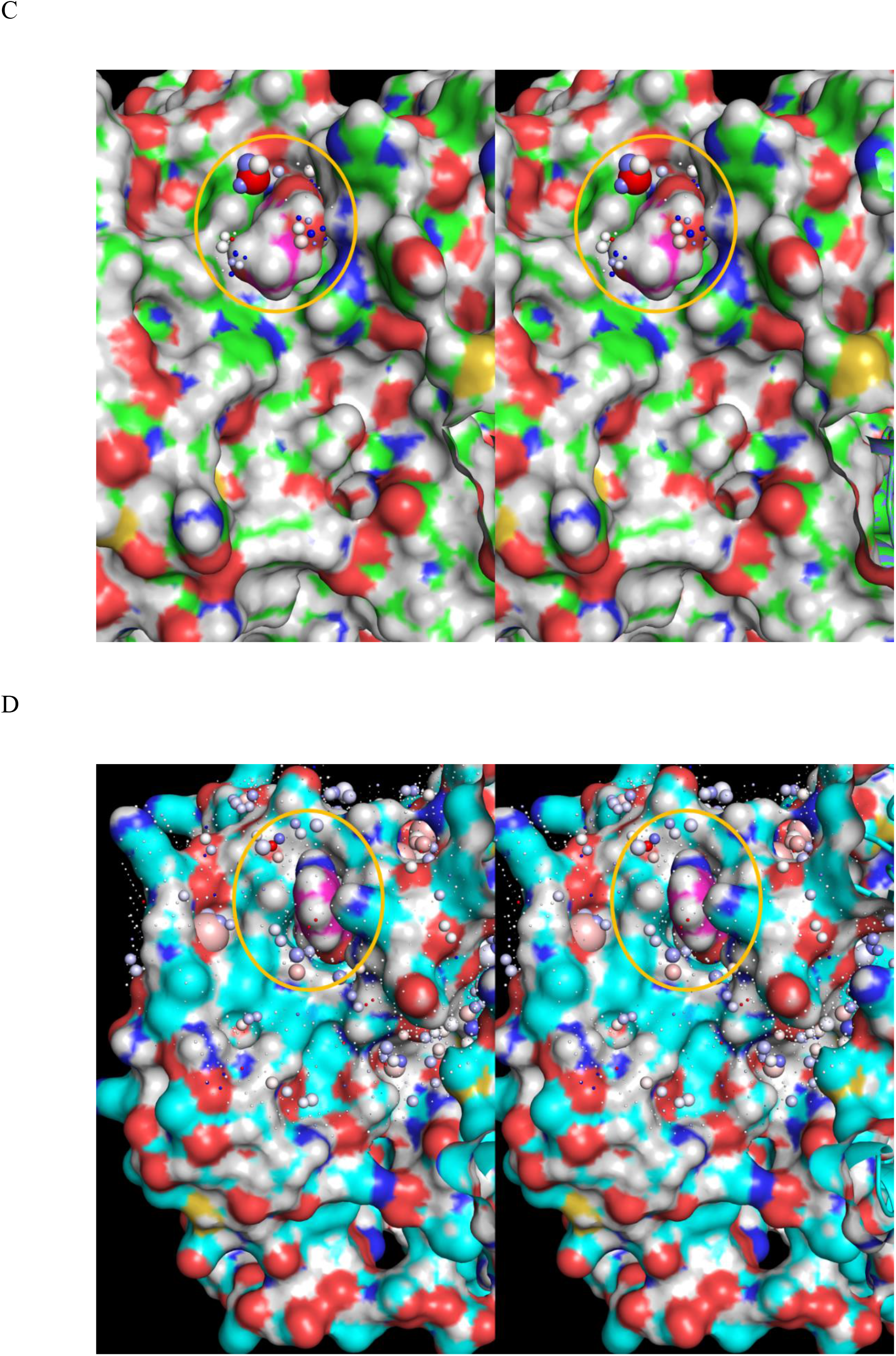
(A) Overlay of structure F (cyan cartoon) with lenalidomide (yellow sticks) and structure K (green cartoon) with uridine (magenta sticks) docked in 5FQD and subjected to a 10 ns MD simulation. (B) Same as A, except showing the HOVs and ULOVs of uridine (magenta sticks). (C) Same as A, except showing the exposed ribose moiety on the substrate binding surface (circled in yellow), together with residual ULOVs and HOVs. The substrate binding surface differs greatly from that of phthalimide-type degraders, suggesting that the native substrate(s) of uridine- bound cereblon differ greatly from neo substrates like CK1α. We postulate that native substrates bind to an extended interface encompassing the deep butterfly-shaped depression visible in the figure (with uridine residing in the top left “wing”). (D) Same as C, except showing the lenalidomide-bound substrate binding surface (circled in yellow) of structure C (magenta and cyan surfaces, respectively) and the residual ULOVs and HOVs.

We docked uridine into the pocket L of structure D as described in Materials and methods, followed by a 10 ns SD simulation (resulting in time-averaged structure K). The uracil moiety of the docked degrader is well-overlaid on the glutarimide ring of lenalidomide, with the ribose ring projecting outward toward the binding surface of the substrate (Figures 37A and B). CK1α, and likely all β-hairpin containing substrates, appear to be highly incompatible with the binary cereblon-uridine complex (the lenalidomide-bound surface of cereblon is shown in Figure 37C for comparison).

Next, we analyzed the partially formed (structure A_closed) state of pocket L, which contains significantly larger voxels than the fully formed state (structure A_closed”) (Figure 38) (noting that this state is observed in our simulations solely in the DDB1-unbound form of cereblon, in which the free protein population is likely tipped further toward the degrader binding-incompetent state). This hypothesis is consistent with the reported weak −5.2 kcal/mol ΔH ^44^ for lenalidomide binding to the DDB1-unbound state.

**Figure 38.**
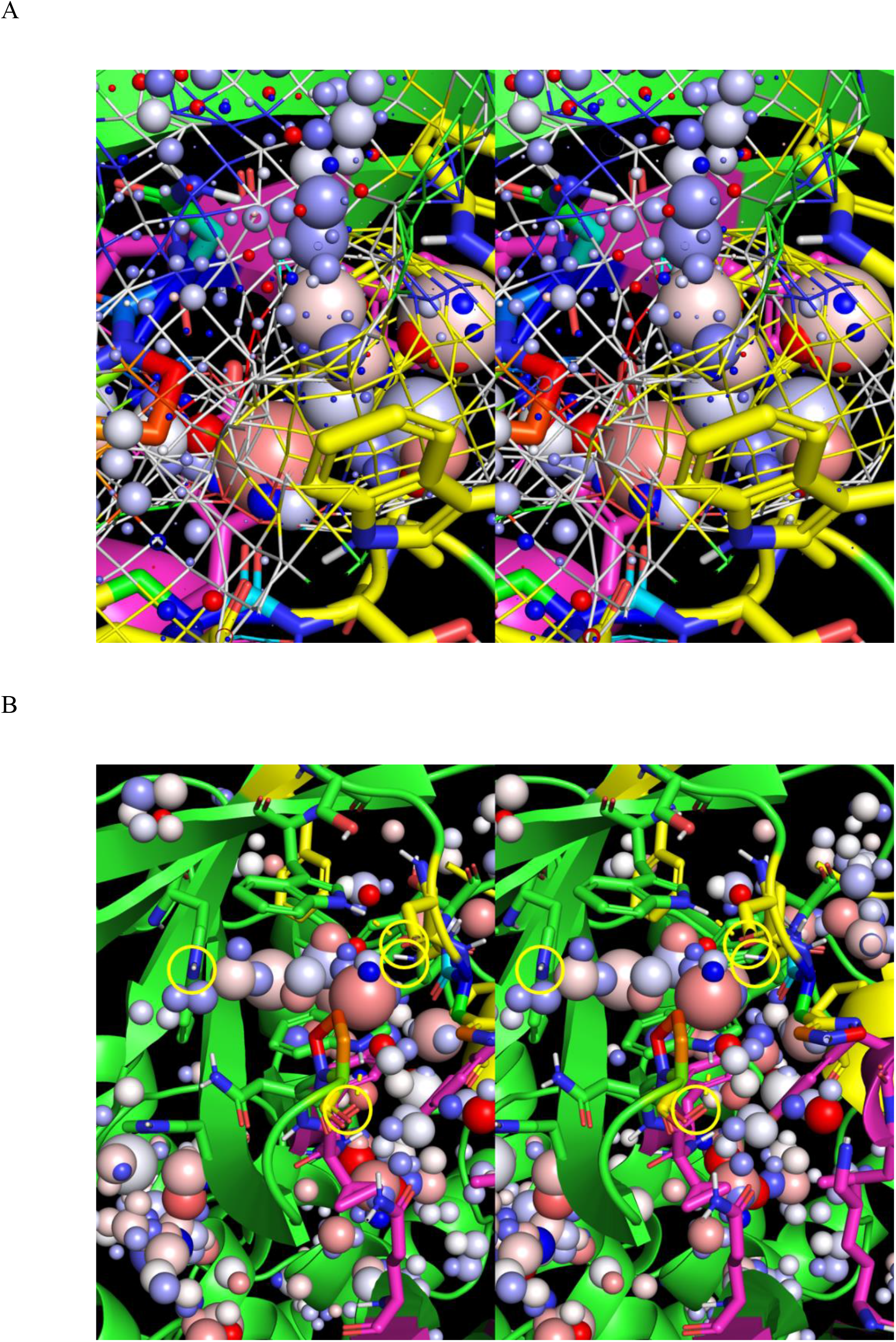
Stereo views of the voxels within the partially formed state of pocket L and fully formed state of pocket T in structure A_closed. Lenalidomide (yellow) from structure C is overlaid for reference. We postulate that the pocket is destabilized by impeded solvation caused by the narrowed opening, manifesting as non-H-bond enriched HOVs. (A) Zoomed-out view of the voxels, highlighting the restricted flow of water to/from the pocket. (B) The solvating water within pocket L putatively resides at the upper end of a free energy Goldilocks zone, in which the aromatic/non-polar pocket lining is partially counter-balanced by protein H-bond groups (circled in yellow).

Pocket T is most prominent in structure A_closed (left arrow in Figure 23C), and is also present in structures A_closed’, C, and F (but is fully absent in structures A_closed”, C’, and C”). This pocket resides at the three-way N-terminal domain-N-terminal peptide-CULT domain interface and is occupied by His332 in the degrader-unbound state. We postulate that His332 serves as a linchpin between the aforementioned substructures, which is expelled from pocket T during degrader association via desolvation of an H-bond enriched HOV (Figure 39A). Solvation of pocket T (exchanged for His332) weakens the three-way interface, resulting in dissociation of the N-terminal peptide and rotation of the CULT domain (Figure 39B).

**Figure 39.**
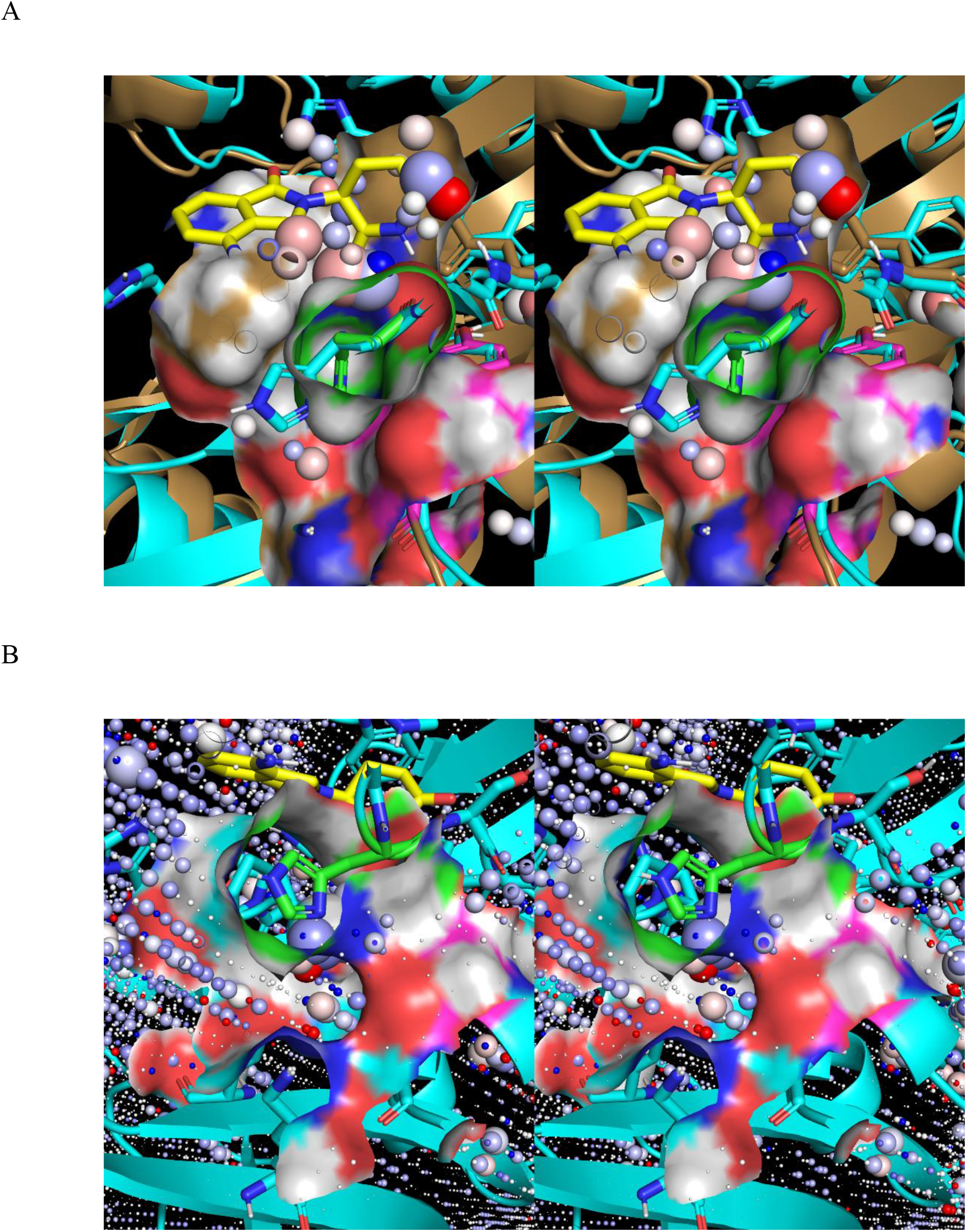
(A) Same as Figure 29C, except showing the vosels within the empty pocket L, including a cluster of HOVs corresponding to H-bond enriched solvation that bridges between pockets L and T, and overlaps the southern O of the glutarimide moiety of lenalidomide (overlaid for reference, and shown as yellow sticks). Optimal H-bond replacement of this water by the glutarimide O putatively results in low cost expulsion thereof via pocket T, accompanied by displacement of His332 from the pocket. (B) The degrader-bound state of structure F, in which His332 (green sticks) has been displaced from pocket T by lenalidomide (yellow sticks). Weakening of the three-way interface between the N-terminal domain, N-terminal peptide (magenta surface), and CULT domain by His332 displacement putatively results in dissociation of these substructures, followed by rotation of the CULT domain to the open state captured in 6H0F and structure A_open (not shown). We postulate that the closed state is adapted for degrader binding (and possibly substratre binding as well), whereas the open state is adapted for substrate ubiquitylation. the closed state present in the starting crystal structure (5FQD) and time-averaged degrader-bound structures is due to crystal packing.

Lastly, we used WATMD to study the solvation properties of the CULT domain within the peri- N-terminal peptide binding interface (the putative “latch”) (Figures 28 and 40A). The N-terminal peptide is stabilized in the CULT-bound state by a large number of ULOVs corresponding to H-bond depleted solvation (small white spheres). We postulate that the peptide dissociates in response to degrader binding via displacement of His332 from pocket T and transient solvation thereof, which weakens the binding interface (Figure 40B). Dissociation of the N-terminal and CULT domains is putatively promoted by dissociation of this peptide, which we postulate facilitates processive substrate poly-ubiquitylation.

**Figure 40.**
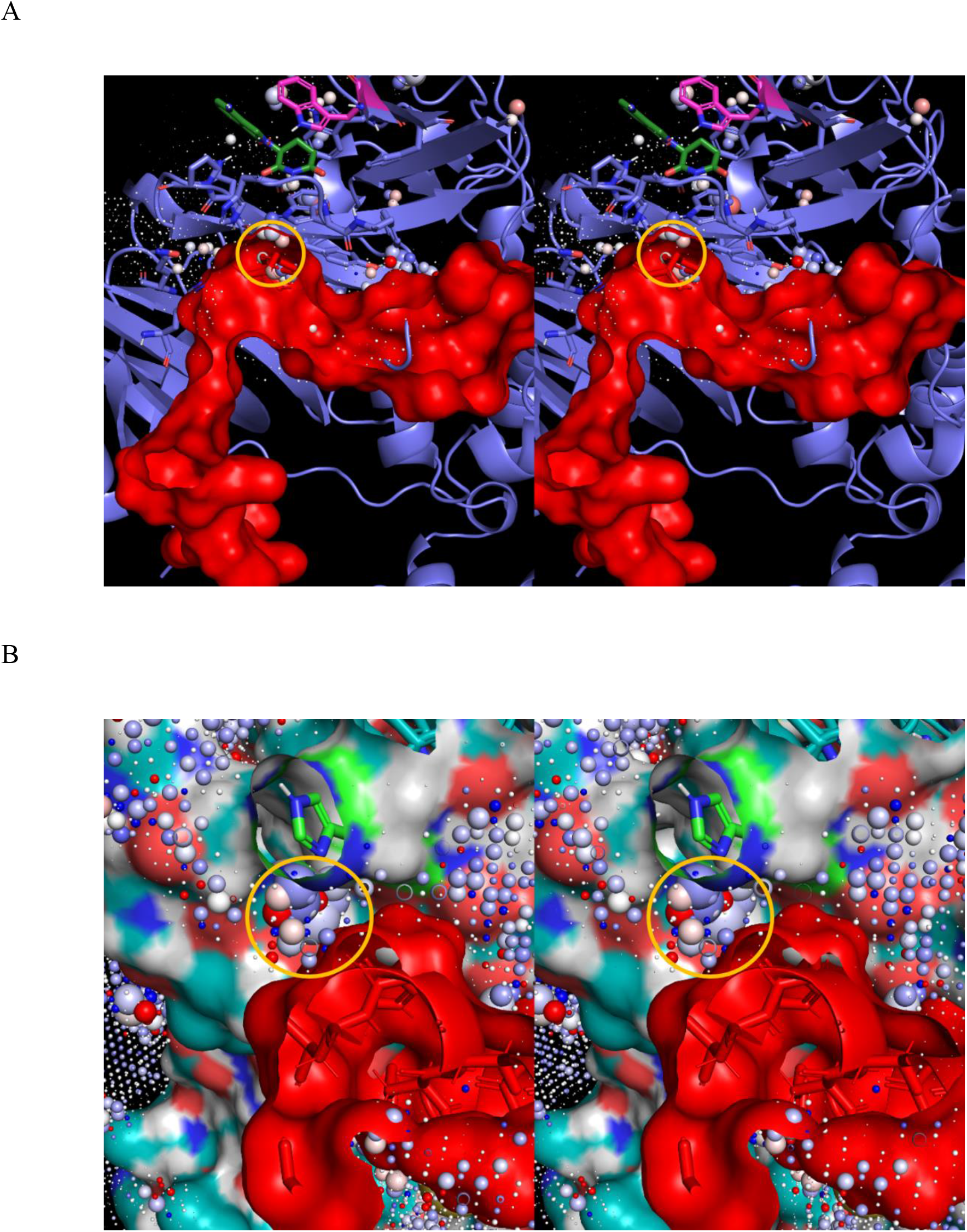
(A) Stereo view of the isolated N-terminal peptide extracted from C (red surface) overlaid on the ULOVs and HOVs of C” (blue cartoon) within the footprint of the N-terminal peptide. The entrance to pocket T is visible at the upper left horizontal section of the peptide (circled in yellow). (B) Close-up stereo view of the entrance to pocket T (circled in yellow) prior to degrader-induced peptide (red surface) dissociation. His332 (green sticks), which is displaced from the pocket, is shown for reference.

## Discussion

### The non-covalent rearrangements of aqueous solutes are powered by solvation free energy

We proposed more than a decade ago that non-covalent intra- and intermolecular rearrangements among the sterically-accessible states of aqueous biomolecules and multi-molecular systems under native cellular conditions are powered principally by solvation free energy.^23^ More recently, we expanded this concept into a general first principles multi-scale theory on the means by which cellular functions are conveyed by networked molecular fluxes (“physical equations”) operating in the NLD regime.^1, 3, 4, 46^ According to our theory, molecular fluxes are powered in a source-sink fashion by covalent free energy and undergo interdependent solvation-powered non-covalent and enzyme-catalyzed state transitions manifesting as cellular states and sub-states (the instantaneous solutions of the multiplexed flux “equations”). In this work, we further extend and exemplify the solvation dynamics concept, in which enthalpic and entropic potential energy is dynamically stored and released in the form of:

1) Impeded or trapped water within dynamically rearrangeable internal channels and cavities, in which exchanges between such water and bulk solvent (the reference state of liquid water) are slowed to varying degrees. Trapped/impeded high energy water manifests as HOVs in our WATMD calculations (not to be confused with HOVs corresponding to low energy solvation at external surfaces).
2) Solvating water that is “repelled” from H-bond depleted solute surfaces to bulk solvent. Such water manifests as ULOVs in our calculations.

High potential energy solvation (relative to that of bulk solvent) resides within a Goldilocks zone, where:

1) Structural states above the Goldilocks zone of free energy form slowly or not at all.
2) Structural states below the Goldilocks zone of free energy are inert (i.e., are stuck in such states).

**Potential energy is released via relaxation of high energy H-bond fields, in which H-bond depleted solvation is expelled (at zero cost) and replaced by non-polar/weakly polar intra- and/or inter-solute groups that likewise exhibit low or no desolvation cost**. Rearrangements necessarily begin and end with the respective storage and release of positive solvation potential energy, including exchanges between:

1) Unfavorable states i and j, in which H-bond depleted, impeded, or trapped solvation expelled from one or more positions in state i appears elsewhere in state j (noting that restrained SD simulations result in artificially exaggerated impedance).
2) Transitions to/from states containing counter-balanced favorable-unfavorable solvation configurations to/from states in which the unfavorable contribution is desolvated at the expense of the favorable contribution (e.g., Trp340 gating in cereblon depicted in Figure 32B).

Molecular populations are distributed across their sterically accessible states under equilibrium conditions according to the free energy of each state relative to the global free energy minimum, or between any states of interest (ΔG_k_). Whereas 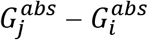 (denoted as scenario 1 (Figure 9A) and 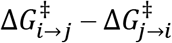 (denoted as scenario 2) (Figure 9B)) are both valid representations of ΔG under equilibrium conditions, the interpretation of measured ΔG or prediction of ΔG based implicitly or explicitly on 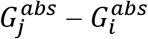 is, in practice, precluded for the following reasons:

1) *G^abs^* resides largely in solvation dynamics (*G^abs^* ≈ *G^bulk^* − (*G^enriched^* + *G^depleted^*) = *δG*), rather than solute structure per se. However, *G^abs^* is implicitly assumed in force-field (FF)- and quantum mechanics (QM)-based computational approaches to reside principally in solute interatomic contacts. As such, ΔG (or ΔΔG among closely related molecules) cannot be predicted reliably using such methods in the absence of data fitting (e.g., free energy perturbation).
2) Water-water, water-solute and solute-solute H-bond contributions to free energy (principally internal energy) are weighted similarly in FF-based methods, and solute-solute contacts are overweighted relative to water-water and water-solute H-bond free energy contributions (noting that *ab initio* QM methods are impractical for explicitly solvated systems). We have provided examples of inter- and intramolecular cereblon rearrangements that are ostensibly driven by overweighted H-bond force-field energies, including rotation of Trp340 into pocket L and retention of the interfacial H-bonds across the hypothetical binary cereblon-CK1α interface (time-averaged structure B). Whereas Trp340 rotation makes sense in the context of pocket L desolvation (to which the AMBER force-field is largely ambivalent), the resemblance of the time-averaged structure B to the starting ternary structure D does not. As such, structural rearrangements predicted by MD simulations are more likely to be right for the wrong reasons (or entirely wrong), unlike SD simulations that are aimed at predicting the properties of H-bond-dominated solvation fields.
3) All of the aforementioned contributions to *G^abs^* depend on <u>dynamic water exchanges</u> between bulk solvent and solvation that are neglected in both FF- and QM-based computational methods, which are limited to static solute structures and solvation configurations. Our WATMD results suggest that the H-bonds of solvating water switch continually among co-localized solute acceptors and donors, either via rotational/translational rearrangements (“smearing”) or during separate exchanges with bulk solvent, manifesting as HOV clusters that, in many cases, exhibit a range of water H and O preferences. H-bond switching reduces entropic losses incurred when solvating water is constrained rigidly in concurrent multi-H-bond interactions (noting that water participates in 3.5 H-bonds/molecule in bulk solvent via rapid partner-partner switching, in which entropic losses are avoided^3^).

Scenario 2 applies to both the biochemically relevant equilibrium and native cell relevant non-equilibrium regimes, whereas scenario 1 is limited strictly to the former regime. Furthermore, the qualitative interpretation of measured ΔG or the prediction of ΔG based on 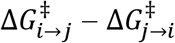 within the Biodynamics framework is possible for the following reasons:

1) An explicit relationship is assumed between 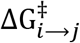 and desolvation and 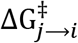 and resolvation during the i → j and j → i state transitions, respectively.
2) SD simulations exploit the fact that water-water H-bonds dominate the free energy landscape in aqueous systems, manifesting as high sensitivity of solvating water to H-bond gains and losses relative to bulk solvent (capable of resolving a single carbon atom difference among solutes^3^).
3) *δG*^*enriched*^ = (*G*^*bulk*^ − *G*^*enriched*^) and *δG*^*depleted*^ = (*G*^*bulk*^ − *G*^*depleted*^) are assumed to be reflected qualitatively in the time-averaged voxel occupancies calculated by WATMD.

The overall non-linear dynamic behavior of solutes and their solvation fields is treated implicitly in SD simulations and WATMD, but not in free energy calculations in which <u>static</u> solute-solvation configurations are assumed. Solvation contributions to predicted structural state preferences are therefore further underestimated relative to the intra- and inter-solute contributions by these methods due to their heavy bias toward maximization of van der Waals contacts, H-bonds, electrostatic contacts, ring-stacking, etc.

In our previous work, we used Biodynamics and WATMD to study the putative intramolecular and observed intermolecular rearrangements of COVID M^pro^ in the non-equilibrium context.^4^ Here, we have used these methods to probe the structure and function of the binary and ternary complexes of the cereblon subunit of the CUL4A/RBX1/DDB1/CRBN E3 ligase, LMW degraders, and the neo substrate CK1α, likewise in the non-equilibrium context. Substrate binding to both proteins, degrader binding to cereblon, and M^pro^ dimerization are powered by <u>external</u> solvation, whereas the up/down state transitions of M^pro^ and release of the N-terminal peptide of cereblon are powered by <u>internal</u> solvation similar to that which we postulated for M^pro^.^4^ We have further postulated that both M^pro^ and cereblon are activated by substrate-induced dimerization and degrader binding, respectively. We propose that HMW-HMW (e.g., protein-protein) and HMW- LMW binding are, in general, powered predominantly by desolvation and resolvation costs underlying k_on_ (or k_1_) and k_off_ (or k_-1_), respectively, which may be regulated by additional inputs (e.g., allosteric activators/inhibitors) governing interfacial surface shape and H-bond group complementarity among the partners. We further propose that the solvation fields across the full complement of aqueous biomolecules constitute a new omics class that we refer to as the “solvome” (form follows function follows free energy).

### Cereblon is switched by degraders from binding incompetent to competent forms

Our findings challenge the widely held view that cereblon is “glued” to its substrates via LMW degraders (i.e., acting as a switch between fast k_-1_ in their absence and slow k_-1_ in their presence). The relatively small fraction of ULOVs contributed by lenalidomide in the cereblon-lenalidomide complex is apparent in Figure 30B (the yellow region of the surface above lenalidomide), consistent with incremental, rather than radical k_-1_ slowing in ternary complex formation. **Furthermore, impeded substrate turnover in the binary complex due to fast k_-1_ relative to k_cat_, rather than slowed k_1_ (where k_1_ remains similar to that of the ternary complex), would result in substrate inhibition of cereblon**. Instead, Type I degraders far more plausibly speed k_1_ (i.e., by lowering the association barrier) versus slow k_-1_ (i.e., by raising the dissociation barrier), noting that slowing k_-1_ from the ternary complex is well-explained by the association-induced release of H-bond depleted water present in the overall cereblon-degrader-CK1α interface. **Therefore, we propose that Type I degraders activate cereblon by aligning the surface shape and key exposed H-bond groups with those of its cognate substrates** (Table 3). The substrate binding surface of binary cereblon-Type I degrader complexes consists of both cereblon and the exposed phthalimide ring of bound synthetic degraders (Figures 41A and B) or the ribose ring of uridine (Figure 41 C), which putatively speed k_1_ (and to a lesser extent, slow k_-1_), and further contribute to substrate selectivity/complementarity. The smaller footprint of BRD4 on the corresponding cereblon-degrader complex (Figure 15C) relative to that of CK1α (Figure 15A) is consistent with insufficient desolvation of H-bond depleted solvation to achieve a physiologically meaningful k_-1_. As such, Type II degraders are expected to both speed k_1_ and slow k_-1_ via expansion of the interface deeper within the target substrate (acting as both “glues” and “activators”).

**Figure 41.**
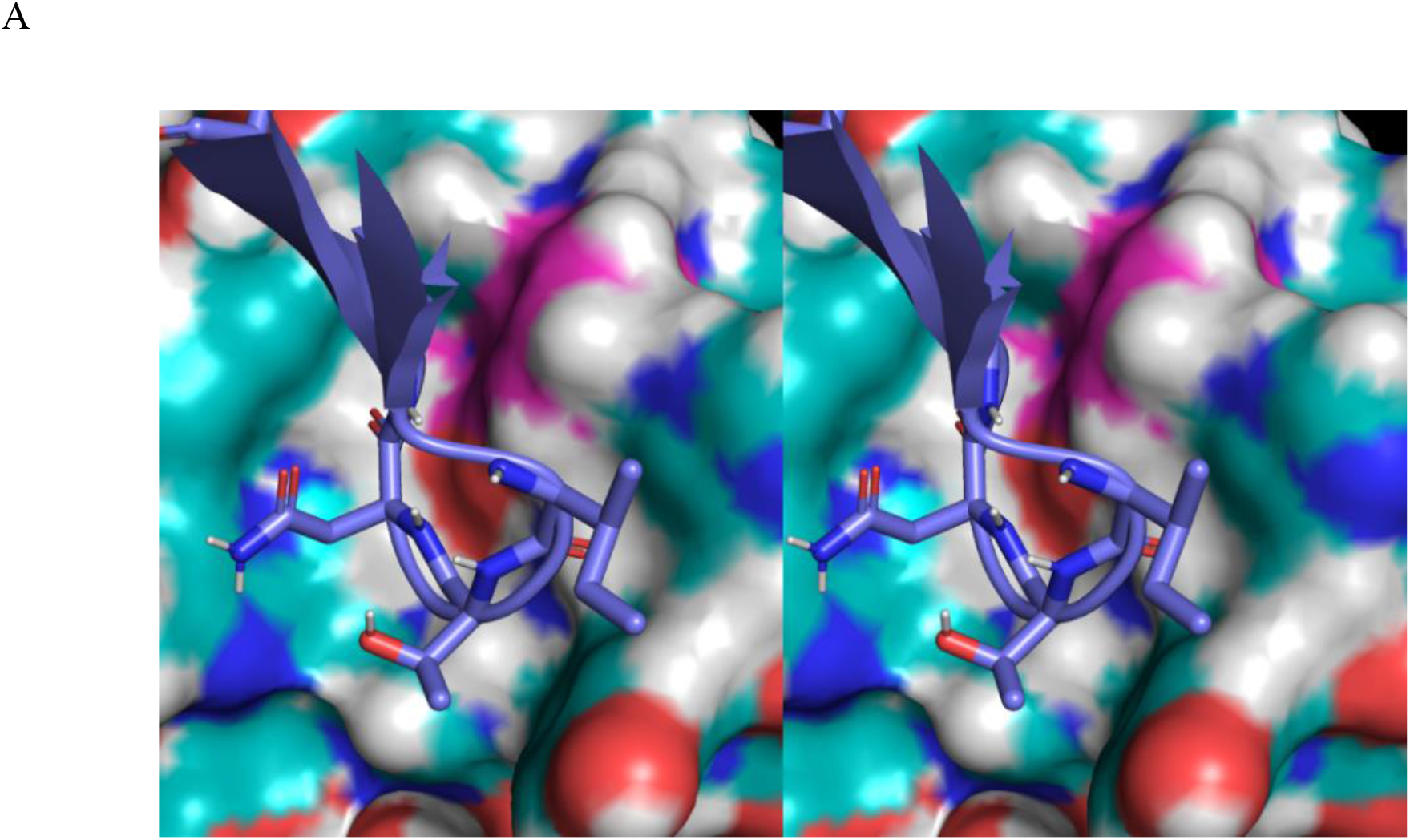
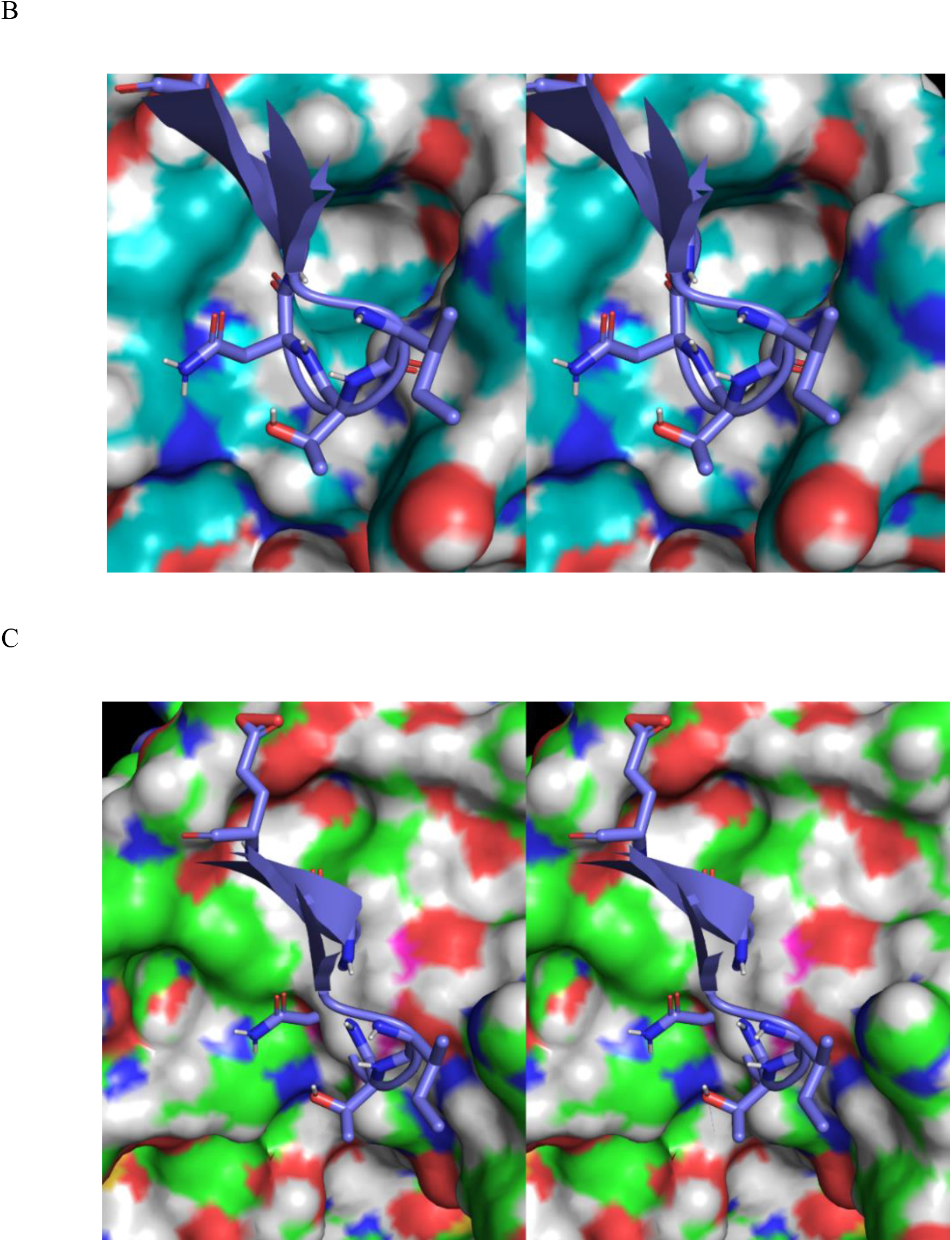
(A) Stereo view of the blended surfaces of cereblon plus lenalidomide (blue and magenta, respectively), together with CK1α (blue cartoon) (structure F). High complementarity between the binary surface and the substrate is apparent. (B) Same as A, except showing the cereblon surface in the absence of the degrader contribution. (C) Same as A, except for docked uridine (structure K) with CK1α from structure F overlaid for reference. The CK1α pocket is completely absent in this structure, consistent with the lack of CK1α binding.

**Table 3.** Implications of “glue” versus “accelerator/activator” mechanisms for substrate-cereblon binding kinetics in the absence (*k*^′^, *k*^′^_1_) and presence (*k*_1_, *k*_−1_) of Type I and II degraders (noting that, based on our rationale, Type I and II degraders are designated as “activator” and “glues”, respectively).

|  |  | Ratio of substrate association and dissociation rate constants in the absence (primed) and presence (unprimed) of degraders |  |
| --- | --- | --- | --- |
| Type | Scenario | $k'_1/k_1$ | $k'_{-1}/k_{-1}$ |
| II | Glue (slows $k_{-1}$) | $\sim 1$ | $\gg 1$ |
| I | Activator (speeds $k_1$) | $\ll 1$ | $\gtrsim 1$ |

## Conclusion

We have extended our previous work on molecular fluxes,^1, 4, 5^ focusing on the general means by which:

1) Cellular behaviors are conveyed by multiplexed multi-flux systems operating in the non-linear dynamics regime via a molecular form of analog computing. Fluxes undergo switchable covalent and non-covalent intra- and intermolecular state transitions governed by the overall system behavior.
2) Switching is powered by desolvation of H-bond depleted water from internal/buried and external surfaces.
3) Unfavorable and favorable free energy is stored in the H-bond network of water solvating both HMW and LMW species (referred to as the “solvation field”), equating to entry and exit barriers underlying the intrinsic rates) to/from non-covalent states.
4) Solvation fields are governed by solute composition and structure (i.e., structure-solvation field relationships).

We exemplified the aforementioned principles via cereblon, the substrate-recruiting subunit of the CUL4A/RBX1/DDB1/CRBN E3 ligase complex, which serves as the sink for both native and neo substrates. In particular, we explored the:

1) Relationships between the extrinsic and intrinsic rates of cereblon binding to LMW Type I and II degraders and substrates (assuming that the partners build and decay over time) using the symbolic mathematical model provided in the Supplementary Information, **focusing on the optimal rates for achieving the maximum dynamic degrader occupancy at the lowest possible degrader concentration (based on the “kinetic tuning” concept that we advanced previously**^1^). We postulate that for Type I degraders, the substrate degradation rate (k_cat(4)_) is optimally >> the rate of downstream substrate-mediated effector activation (k_16_), and the rate of binary cereblon-degrader formation (k_9_) is optimally ≈ the rate of cereblon buildup (k_7_). We postulate that for Type II degraders the rate of degrader-substrate binding (k_20_) is ideally > the rate of degrader-cereblon binding (k_9_), which simultaneously promotes <u>substrate inhibition and degradation</u>, while limiting cereblon inhibition.
2) Qualitative structure-solvation field relationships of cereblon and its binding partners to deduce the solvation-powered dynamic binding mechanism (analogous to our previous work on COVID M^pro^).^4^ We postulate that occupancy of the degrader pocket of cereblon shifts the population toward the substrate binding-competent state, which is otherwise populated predominantly in degrader and substrate binding-incompetent states. We further postulate that degrader binding is powered by the release of high potential energy stored as H-bond depleted solvation within the empty degrader pocket (counter-balanced by co- localized H-bond enriched solvation), which is transduced into alignment between the interfacial H-bond groups of cereblon with those of cognate substrates, followed by release and rotation of the N-terminal peptide from the CULT domain. We proposed that Trp340 and His332 play key roles in “gating” the empty degrader pocket L and the adjacent water-filled pocket T. Last but not least, we showed that the solvation fields of uridine, lenalidomide, and thalidomide are highly complementary to that of the degrader pocket, translating to low (but specific) desolvation costs of the associating degrader species.

