## Supplemental File for "Solvation dynamics-powered structure and function of multi-molecular cellular systems exemplified by non-equilibrium cereblon-degrader-CK1α ternary complex formation"

**Reaction scheme for Type I degraders that bind in order of cereblon-degrader + substrate:**

source ⟶ CUL4A/RBX1/DDB1

source ⟶ native E3

source ⟶ DUB (deubiquitylase) ⟶ sink

source ⟶ substrate (sink consists of proteasomal degradation)

source ⟶ substrate_effector (downstream partner of the substrate) ⟶ sink

source ⟶ cereblon ⟶ sink

source ⟶ degrader ⟶ sink (PK curve)

CUL4A/RBX1/DDB1 + cereblon ⇌ CUL4A/RBX1/DDB1-cereblon

CUL4A/RBX1/DDB1-cereblon + degrader ⇌ CUL4A/RBX1/DDB1-cereblon-degrader

CUL4A/RBX1/DDB1-cereblon-degrader + substrate ⇌ CUL4A/RBX1/DDB1-cereblon-degrader-substrate ⟶ CUL4A/RBX1/DDB1-cereblon-degrader-substrate-$(Ub)_n$

CUL4A/RBX1/DDB1-cereblon-degrader-substrate-$(Ub)_n$ ⟶ CUL4A/RBX1/DDB1-cereblon-degrader + substrate-$(Ub)_n$

substrate-$(Ub)_n$ + DUB ⇌ substrate-$(Ub)_n$-DUB

substrate-$(Ub)_n$-DUB ⟶ substrate + DUB

substrate-$(Ub)_n$ ⟶ proteasome

substrate + substrate_effector ⇌ substrate-substrate_effector

substrate + native E3 ⇌ substrate-native E3 ⟶ substrate-$(Ub)_n$ + native E3

**Ordinary differential equations for Type I degraders:**

$d[\text{CUL4A/RBX1/DDB1}](t)/dt = -k_0 \cdot [\text{CUL4A/RBX1/DDB1}_{source}](t)$

$d[\text{DUB}](t)/dt = -k_1 \cdot [\text{DUB}_{source}](t) - k_2 \cdot [\text{DUB}](t)$

$d[\text{substrate}](t)/dt = -k_3 \cdot [\text{substrate}_{\text{source}}](t) + k_{cat(4)} \cdot [\text{DUB-substrate-(Ub)}_n](t)$

$d[\text{cereblon}](t)/dt = -k_4 \cdot [\text{cereblon}_{\text{source}}](t) - k_5 \cdot [\text{cereblon}](t) - k_7 \cdot [\text{CUL4A/RBX1/DDB1}](t) \cdot [\text{cereblon}](t) + k_8 \cdot [\text{CUL4A/RBX1/DDB1-cereblon}](t)$

$d[\text{degrader}](t)/dt = -k_6 \cdot [\text{degrader}_{\text{source}}](t) - k_{cat(1)} \cdot [\text{degrader}](t)$ (absorption/clearance)

$d[\text{CUL4A/RBX1/DDB1-cereblon}](t)/dt = k_7 \cdot [\text{CUL4A/RBX1/DDB1}](t) \cdot [\text{cereblon}](t) - k_8 \cdot [\text{CUL4A/RBX1/DDB1-cereblon}](t)$

$d[\text{CUL4A/RBX1/DDB1-cereblon-degrader}](t)/dt = k_9 \cdot [\text{CUL4A/RBX1/DDB1-cereblon}](t) \cdot [\text{degrader}](t) - k_{10} \cdot [\text{CUL4A/RBX1/DDB1-cereblon-degrader}](t)$

$d[\text{CUL4A/RBX1/DDB1-cereblon-degrader-substrate-(Ub)}_n](t)/dt = k_{11} \cdot [\text{CUL4A/RBX1/DDB1-cereblon-degrader}](t) \cdot [\text{substrate}](t) - k_{12} \cdot [\text{CUL4A/RBX1/DDB1-cereblon-degrader-substrate}](t) - k_{cat(2)} \cdot [\text{CUL4A/RBX1/DDB1-cereblon-degrader-substrate}](t)$

$d[\text{degraded-substrate-(Ub)}_n](t)/dt = -k_{13} \cdot [\text{proteasome}](t) \cdot [\text{substrate-(Ub)}_n](t) + k_{cat(3)} \cdot [\text{substrate-(Ub)}_n](t)$

$d[\text{DUB-substrate-(Ub)}_n](t)/dt = k_{14} \cdot [\text{DUB}](t) \cdot [\text{substrate-(Ub)}_n](t) - k_{15} \cdot [\text{DUB-substrate-(Ub)}_n](t)$

$d[\text{substrate-substrate\_effector}](t)/dt = k_{16} \cdot [\text{substrate}](t) \cdot [\text{substrate\_effector}](t) - k_{17} \cdot [\text{substrate-substrate\_effector}](t)$

$d[\text{substrate-native E3}](t)/dt = k_{18} \cdot [\text{substrate}](t) \cdot [\text{native E3}](t) - k_{19} \cdot [\text{substrate-native E3}](t)$

**Reaction scheme for Type II degraders that bind in order of substrate-degrader + cereblon and cereblon-degrader + substrate (in addition to Type I):**

degrader + substrate $\rightleftharpoons$ degrader-substrate

CUL4A/RBX1/DDB1-cereblon + degrader-substrate $\rightleftharpoons$ CUL4A/RBX1/DDB1-cereblon-degrader-substrate

**Ordinary differential equations for Type II degraders:**

d[degrader-substrate](t)/dt = $k_{20}$ · [degrader](t) · [substrate](t) – $k_{21}$ · [degrader-substrate](t)

– $k_{22}$ · [CUL4A/RBX1/DDB1-cereblon](t) · [degrader-substrate](t) + $k_{23}$ · [CUL4A/RBX1/DDB1-cereblon-degrader-substrate](t)

d[CUL4A/RBX1/DDB1-cereblon-degrader-substrate](t)/dt = $k_{22}$ ·

[CUL4A/RBX1/DDB1-cereblon](t) · [degrader-substrate](t) – $k_{23}$ ·

[CUL4A/RBX1/DDB1-cereblon-degrader-substrate](t)
